# Towards understanding interactions in a complex world: Design and analysis of multi-species functional response experiments

**DOI:** 10.1101/2023.12.19.571428

**Authors:** Benjamin Rosenbaum, Jingyi Li, Myriam R. Hirt, Remo Ryser, Ulrich Brose

## Abstract

1. The functional response describes feeding rates of consumers as a function of resource density. While models for feeding on a single resource species are well studied and supported by a large body of empirical research, consumers feeding on multiple resource species are ubiquitous in nature. However, laboratory experiments designed for parameterizing multi-species functional responses (MSFR) are extremely rare, mainly due to logistical challenges and the non-trivial nature of their statistical analysis.
2. Here, we describe how these models can be fitted to empirical data in a Bayesian framework. Specifically, we address the problem of prey depletion during experiments, which can be accounted for through dynamical modeling. In a comprehensive simulation study, we test the effects of experimental design, sample size and noise level on the identifiability of four distinct MSFR models. Additionally, we demonstrate the method’s versatility by applying it to a list of empirical datasets.
3. We identify experimental designs for feeding trials that produce the most accurate parameter estimates in two- and three-prey scenarios. Although noise introduces systematic bias in parameter estimates, model selection performs surprisingly well for the four MSFRs, almost always identifying the correct model even for small datasets.
4. This flexible framework allows the simultaneous analysis of feeding experiments from both single- and multi-prey scenarios, either with or without prey depletion. This will help to elucidate mechanisms such as prey selectivity, prey switching and their implications for food web stability and biodiversity. Our approach equips researchers with the appropriate statistical tools to improve the understanding of feeding interactions in complex ecosystems.

## Introduction

In ecological communities, species engage in diverse interactions, forming complex networks that underpin the functioning and resilience of ecosystems. In the realm of food web ecology, these interactions are characterized by trophic links between species. Understanding the constraints on the strength of these links, which determine the flow of energy and matter through ecological systems, is crucial for predicting community structure and ecosystem functioning.

Feeding rates are commonly captured by the “functional response”, which describes consumption of prey based on their density. Since its introduction (Solomon 1949), numerous models have been proposed (Jeschke, Kopp & Tollrian 2002), with a sustained interest until today (DeLong 2021; Abrams 2022). The most prevalent versions of the functional response are the Holling type 2 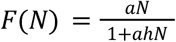 and the Holling type 3 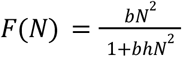 (Holling 1959), where *a* is an attack rate (the fraction of habitat cleared from prey per time), *h* is handling time (the time the predator needs to consume and digest the prey), and *b* is a constant. The type 3 form arises from the type 2, when attack rates themselves are density-dependent, i.e. the attack rate of a predator increases with prey density as *a* = *bN*, but also a generalized formulation *a* = *bN*^*q*^ with a flexible exponent *q* > 0 exists (Real 1977). The occurrence of sigmoidal type 3 functional responses has been explained by learning behavior of the predator when confronted with a prey species more frequently, or switching behavior between different prey species (Murdoch & Oaten 1975). The difference between the functional response types has gained particular importance due to their effects on population and food web stability (Kalinkat *et al*. 2023).

Although classical functional response models measure interactions of a predator with a single prey species, nature is more complex. Predators typically have a variety of prey options, and their feeding habits (and thus interaction strengths) are influenced not just by the abundance of one prey species but by the collective presence and density of all available prey (Abrams 2022). For instance, the GATEWAy database of 290 food webs (Brose *et al*. 2019) contains 2,917 predators, of which 92% are reported to feed on more than one prey species. Feeding rates on a focal resource are generally expected to decrease in the presence of alternative resources. Their mathematical descriptions *F*_*i*_ (*N*_1_,…, *N*_*m*_) of feeding on a species *i* depend not only density *N*_*i*_, but also on other prey densities *N*_*j*_ (*j* ≠ *i*), leading to multi-species functional responses (MSFR), (Gentleman *et al*. 2003; Koen-Alonso 2007). MSFRs are capable of modeling important concepts such as prey preference (Chesson 1983), switching (Murdoch 1969) or optimal foraging (Holt 1983), with implications for species coexistence and ecosystem stability. A plethora of MSFR models covering these topics have been proposed, with examples in (Gentleman *et al*. 2003; Morozov & Petrovskii 2013; Vallina *et al*. 2014; Ryabov, Morozov & Blasius 2015; Baudrot *et al*. 2016; Stouffer & Novak 2021). To our knowledge, none of these simultaneously included (1) sigmoidal behavior in single-prey scenarios, (2) active prey switching (which cannot be predicted from single-prey information) in multi-prey scenarios, and (3) prey species-specific maximum feeding rates (e.g. due to differences in body size). Here, we developed a new flexible but still tractable MSFR with these features that generalizes some of previous models.

To empirically measure and parameterize these mathematical descriptions of functional responses, feeding trials in highly controlled laboratory experiments are commonly used (Griffen 2021). In single-prey scenarios, typically one or more predator individuals are allowed to feed on a given prey density for a fixed period of time and numbers of eaten prey are recorded. Experiments are replicated along a gradient of prey densities and functional response models are fitted to the observed data of eaten vs. offered prey abundances. This allows parameterizing models, testing model formulations, and tackling questions of e.g. allometric scaling or temperature dependence (Daugaard, Petchey & Pennekamp 2019; Uszko, Diehl & Wickman 2020; Sohlström *et al*. 2021; Kratina *et al*. 2022). Numerous single-prey experiments have been conducted in recent decades. The FoRAGE database compiles more than 2,500 taxonomically diverse datasets of functional responses from 70 years covering terrestrial, marine and freshwater habitats (Uiterwaal *et al*. 2022).

In multi-prey scenarios, accurate parameterization of MSFRs is required, not least because they are the backbone of widely food web models, which often include multi-species extensions of the Holling type 3 response (Martinez 2020; Gauzens *et al*. 2023). Here, population dynamics may be sensitive to parameters such as the exponent *q*, with drastic consequences for biodiversity (Williams & Martinez 2004). However, empirical evidence for MSFR models remains extremely rare, see e.g. (Stouffer & Novak 2021) and (Coblentz 2020) for non-exhaustive lists of 30 and 20 multi-species datasets, respectively. While the collection and analysis of field data can be accomplished, e.g., (Smout *et al*. 2010; Novak *et al*. 2017; Preston *et al*. 2018; Beardsell *et al*. 2022; Uiterwaal & DeLong 2024) and features its own difficulties, even experiments in highly controlled laboratory settings come with logistical and statistical challenges.

First, these models tend to be data-hungry with potentially hundred(s) of individual feeding trials, each requiring the counting of up to hundreds of prey individuals of different species. Second, after data collection researchers are required to perform some sort of nonlinear model fitting which might exceed a classical statistics curriculum (but see, e.g., (Smout *et al*. 2010; Baudrot *et al*. 2016; Smith & Smith 2020) as examples of Bayesian inference). Here, two factors have been widely disregarded in the MSFR literature: experimental design and prey depletion. First, whereas response-surface designs have been used for estimating species interactions (Inouye 2001; Okuyama & Bolker 2013; Terry, Morris & Bonsall 2017; Uszko *et al*. 2020; Novak & Stouffer 2021a; Kleinhesselink *et al*. 2022), their potential effects on the accuracy of MSFR parameter estimation have not been explored yet. Second, we know from single-prey experiments that model fitting without correcting for prey depletion leads to biased estimates (Rosenbaum & Rall 2018). If eaten prey are not replaced instantly, prey abundances and hence feeding rates are not constant over the course of an experiment, but decrease. In single-prey experiments, statistical methods accounting for prey depletion are available (Rogers 1972; Bolker 2008; Okuyama & Ruyle 2011; Pritchard *et al*. 2017; Rosenbaum & Rall 2018; Uszko *et al*. 2020). Closed-form solutions exist for simple models such as the type 2 response. More complex formulations (e.g. the flexible type 3 response using *a* = *bN* ^*q*^) are treated by predicting prey density dynamically (i.e. time-dependent *N*(*t*)) via ordinary differential equations (ODEs). The rate of change 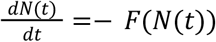 is expressed by the functional response, and the equation can be solved for *N*(*t*) numerically (by a computer algorithm).

In multi-prey scenarios, biases might be exacerbated, since both prey species are depleted by the predator in a nonlinear and interdependent way over the course of an experiment, unless eaten prey items (of the correct species) are constantly replaced to keep densities fixed (which increases logistical effort). Here, we propose to fit ODE-based predictions of eaten prey to observed numbers of both prey species in a two-prey scenario, extending the approach from, e.g., (Rosenbaum & Rall 2018) to the multi-prey case. Feeding rates on both species decline dynamically due to decreasing prey availability. Any density-dependent MSFR model can be included in a coupled set of differential equations

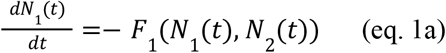

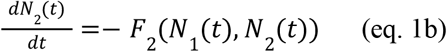

which can be solved numerically to predict the number of prey *N*_1_ (*T*), *N*_2_ (*T*) remaining at the end of the experiment with duration *T*, given the initial prey densities *N* _1_ (0), *N*_2_ (0). Their difference yields predictions of eaten prey (*N*_*E*,1_, *N*_*E*,2_), which are fitted to the observed data to estimate model parameters such as attack rates, handling times and preferences.

We here integrate previous approaches, show how MSFR models can be fitted to empirical data and assess the method’s accuracy. We conducted a simulation study following a “virtual ecologist” approach (Zurell *et al*. 2010), by generating realistic datasets of experiments *in silico* with a predator feeding on two prey species without prey replacement, to answer two questions: First, whether MSFR model parameters can be estimated correctly by fitting ODE-based model predictions and, second, through which experimental design we can obtain the best parameter estimates. As test cases we used multi-prey extensions of the Holling type 2, type 3 and a prey switching model. In addition, we developed a new, more flexible MSFR that generalizes these models (Fig. 1). In a full factorial approach, we tested (1) these four MSFR models of increasing complexity on (2) six experimental designs with (3) five levels of sample size. (4) Data was simulated with three levels of noise. Each of these combinations was replicated 500 times. We used Markov Chain Monte Carlo sampling (MCMC) to fit ODE-based predictions to each of the resulting 180,000 datasets in R in a Bayesian framework with Stan (Stan Development Team 2023a), which has a built-in numerical ODE solver. Additionally, we investigated scenarios with a predator feeding on three prey species in 48,000 datasets. We complement the simulation study with test cases using published experimental datasets.

**Figure 1:**
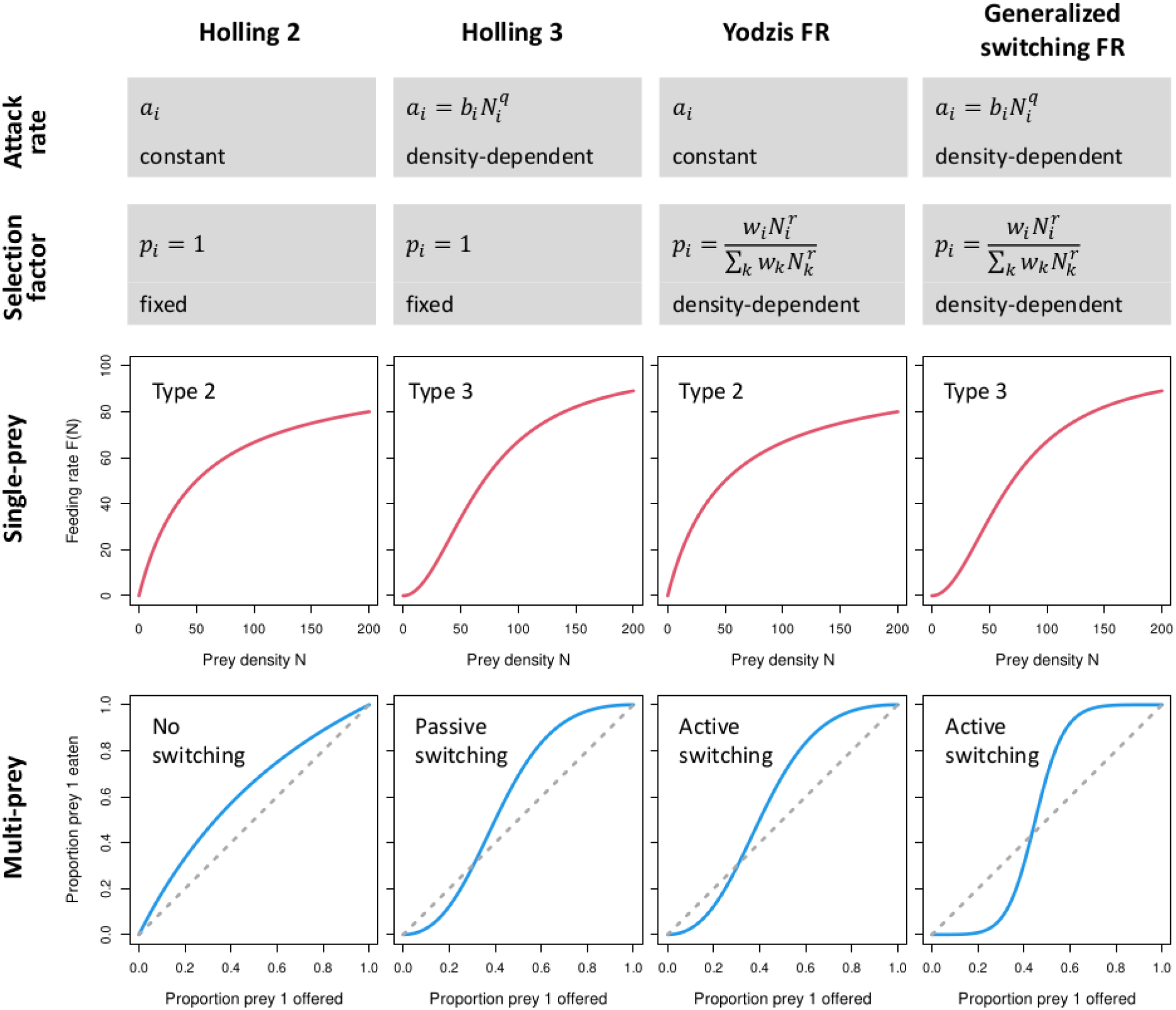
Four MSFR models of the general formulation as in eq. 2 with attack rates *a*_*i*_ and selection factors *p*_*i*_ either constant or density-dependent. In single-prey scenarios, they collapse to a type 2 or type 3 FR. In multi-prey scenarios, prey preference and switching can be passive (as predicted from single-prey scenarios) or active (distinct or more pronounced than in single-prey scenarios).

We find that parameters of our tested MSFR models are estimated accurately in low-noise scenarios. In high-noise scenarios, however, most parameter estimates are systematically biased, but model selection still identifies the correct model even in small datasets. We provide practical suggestions for experimental designs of two- and three-prey feeding trials, that yield the most accurate estimates across all scenarios. Additionally, we provide a detailed tutorial and ready-to-use R code to facilitate the method’s application.

## Materials and Methods

### Multi-species functional response models

(Koen-Alonso 2007) mechanistically derived a generalized form of the MSFR by dividing a predator’s total foraging time into searching time and time spent handling prey individuals of *m* different species, following (Holling 1959). Then, by expressing the capture rate (number of prey eaten divided by searching time) as *C*_*i*_ = *p*_*i*_ *a*_*i*_ *N*_*i*_, where *p*_*i*_ is a dimensionless selection factor (possibly depending on all resource densities *N*_1_,…, *N*_*m*_) and *a*_*i*_ is the attack rate (*a*_*i*_ *N*_*i*_ depending on *N*_*i*_ only), the feeding rate on species *i* reads

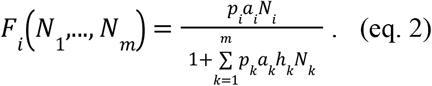

Keeping *p*_*i*_ and *a*_*i*_ as constants or making either of them density-dependent produces different classes of MSFRs, which collapse to either type 2 or type 3 responses in single-species scenarios, and can exhibit different types of prey switching in multi-species scenarios (Fig. 1).

Prey switching is defined through preferences 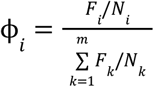 and their pairwise ratio 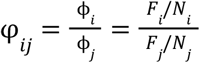 (Murdoch 1969; Gentleman *et al*. 2003; Koen-Alonso 2007). If φ_*i,j*_ is a constant and not density-dependent, then the composition of a predator’s diet 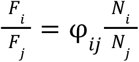 is relative to the proportion of available prey 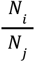. E.g., φ _*i,j*_> 1 would mean that species *i* is always preferred over species *j* and consumed more often than its relative abundance in the environment would predict. If, however, φ_*i,j*_ is density-dependent, a transition from φ_*i,j*_ < 1 to φ_*i,j*_ > 1 is possible and a switch from relative avoidance to relative preference of species *i* can occur when the composition of available prey changes. Preferences and switching are called passive when they can be explained by single-species information alone (e.g. differences in species-specific attack rates). Active switching, on the other hand, requires multi-species information and cannot be predicted from single-species data alone (Gentleman *et al*. 2003).

#### Holling type 2 MSFR

In its simplest expression of eq. (2), we assume constant attack rates *a*_*i*_ and let all *p*_*i*_ = 1 fixed, which produces a multi-species Holling type 2 response (Fig. 1a)

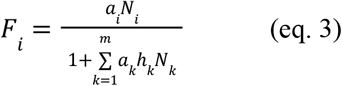

e.g. (Gentleman *et al*. 2003). Note that we could not estimate free constants *p*_*i*_ and *a*_*i*_ both from data since they only appear as a product *a*_*i*_ *p*_*i*_ in eq. (2). Estimates would be perfectly negatively correlated and therefore unidentifiable. In single-prey scenarios (all *N*_*j*_ = 0, *j* ≠ *i*), the response collapses to the hyperbolic Holling-2 FR 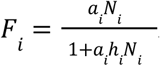. This MSFR does not produce prey switching, because relative prey preferences 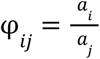 are constant and the ratio of attack rates determines relative prey preference or avoidance. Consequently, this MSFR is determined by single-prey parameters *a*_*i*_ and *h*_*i*_, and could theoretically be estimated from single-prey feeding trials alone.

#### Holling type 3 MSFR

Allowing attack rates 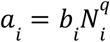 to be density-dependent with a coefficient *b*_*i*_ and a positive exponent *q*, while still keeping all *p* _*i*_= 1 fixed, generates a multi-species Holling type 3 response (Fig. 1b)

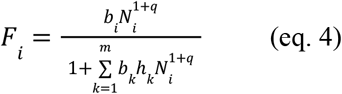

e.g. (Gentleman *et al*. 2003). For *q* = 0 it reduces to the Holling-2 MSFR (replacing *b*_*i*_ by *a*_*i*_). In single-prey scenarios, the response collapses to a sigmoidal Holling-3 FR 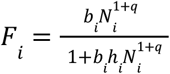. Relative prey preferences 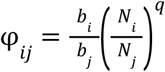 are density-dependent for all *q* > 0 and can produce prey switching. Since all parameters (*b*_*i*_, *h*_*i*_, *q*) could theoretically be estimated from single-prey data, switching is passive.

#### Yodzis MSFR

For another MSFR (Koen-Alonso & Yodzis 2005), named Yodzis functional response in (Koen-Alonso 2007), selection factors 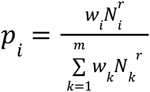 are made density-dependent (for positive *r* > 0), while attack rates *a*_*i*_ are constants. This modifies capture rates *C*_*i*_ = *p*_*i*_ *a*_*i*_ *N*_*i*_ depending on species composition. The denominator ensures 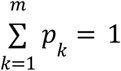 independent of prey composition, and dimensionless preference weights are required to satisfy 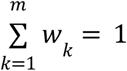. The model can be written as

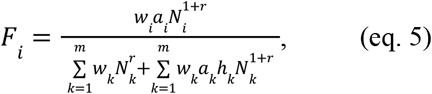

see Fig. 1c. For *r* = 0 it reduces to a Holling-2 MSFR (replacing *w*_*i*_ *a*_*i*_ by *a*_*i*_). In single-prey scenarios, *p*_*i*_ reduces to 1 (only species *i* present) and the model collapses to a Holling-2 FR even for positive *r* > 0, which cancels out just like the weights *w*_*i*_. In multi-prey scenarios however, prey switching can occur because the relative preference 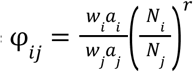 is density-dependent, with exponent *r* controlling the abruptness of switching. Single-prey data does not hold any information on parameters *w*_*i*_ and *r*. Therefore, information from multi-prey scenarios is required to fully determine this MSFR, making it an active-switching response.

#### Generalized switching MSFR

Finally, we can make both the attack rates 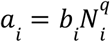 and the selection factors 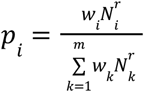 density-dependent, also making capture rates *C*_*i*_ = *p*_*i*_ *a* _*i*_*N*_*i*_ dependent on prey composition. This new MSFR (Fig. 1d)

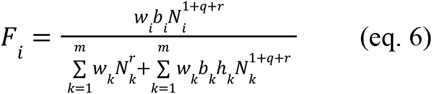

generalizes the three previous models. For *q* = *r* = 0 it reduces to a Holling-2 MSFR (replacing *w* _*i*_*b*_*i*_ with *a* _*i*_), for *q* > 0 and *r* = 0 it reduces to a Holling-3 MSFR (replacing *w*_*i*_ *b*_*i*_ by *b*_*i*_), and for *q* = 0 and *r* > 0 it reduces to the Yodzis MSFR (replacing *b*_*i*_ by *a*_*i*_). It collapses to a type 3 FR in single-prey scenarios, but unlike the Holling-3 MSFR, the switching is active: relative preference 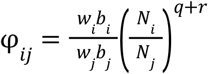 is density-dependent and parameters *w*_*i*_ and *r* can only be informed with multi-prey data. While *q* > 0 controls the sigmoidal shape in single-prey scenarios, switching can be even more pronounced in multi-prey scenarios via the exponent *q* + *r*.

### Experimental design

We tested six experimental designs for combining initial abundances of two prey species (Fig. 2). The maximum abundance of each species was 200, but we assume that researchers choose a system-specific maximum for their experiments, so that feeding rates show a certain level of saturation. Abundances were always rounded to the nearest integer. For each design, we tested numbers of *n* = 64, 128, 256, 512 and 1024 feeding trials. Although our highest sample sizes could, in practice, only be achieved with unrealistically high effort, we wanted to include these scenarios to quantify the theoretical limits of estimation accuracy.

**Figure 2:**
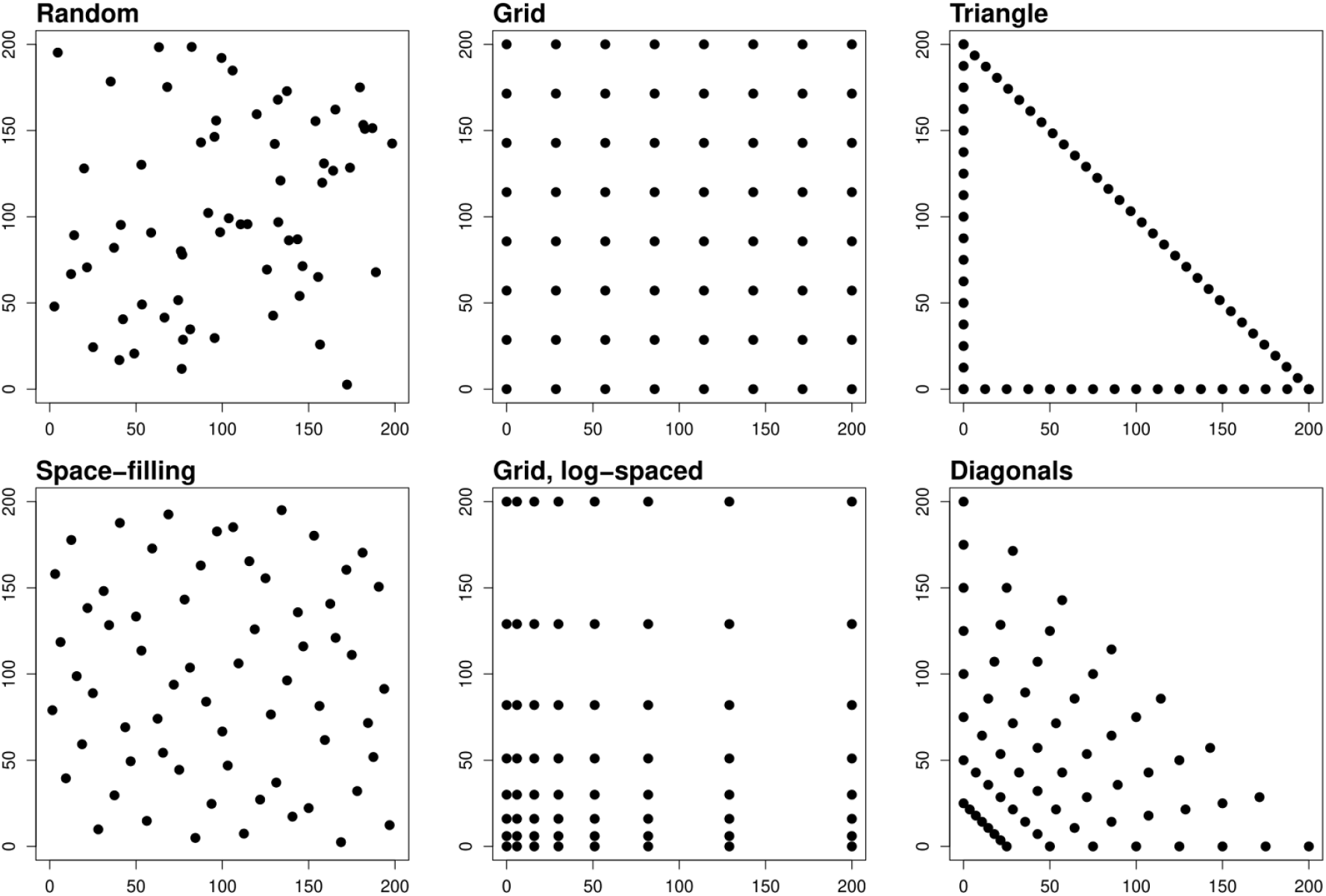
Six experimental designs for offered numbers of prey (*N*_1_, *N*_2_) for two-prey experiments with a maximum initial abundance of 200. Sample size *n*=64, each.

1. Random design. For each feeding trial, abundances *N*_0,1_ and *N*_0,2_ are each drawn from a uniform distribution *U*(0, 200).
2. Space-filling design. We use a Halton sequence, a quasi-random number sequence that covers a rectangle more evenly than a purely random distribution (R-package “randtoolbox” (Dutang & Savicky 2023)).
3. Grid. Abundances are placed equidistantly on an 8x8 factorial grid. 1, 2, 4, 8 or 16 replicates of each combination were used to achieve given sample sizes *n*.
4. Log-spaced grid. Distances between grid points are not identical, but increase by factor 1.5 with increasing abundance. Thus, more experiments are distributed at lower abundance levels. Again, we used 1, 2, 4, 8 or 16 replicates of each combination.
5. Triangle. Feeding trials are conducted in single-prey scenarios for each species (*n*/4 trials each, equidistant), and on a diagonal line in multi-prey scenarios (*n*/2 trials, equidistant), where the total number of initially available prey *N*_0,1_ + *N* _0,2_ = 200 is constant.
6. Diagonals. Experiments are placed equidistantly along 8 diagonals, each representing a different constant level of total prey *N*_0,1_ + *N*_0,2_ (*n*/8 experiments each).

### Data simulation

For each simulated dataset, model parameters were drawn from random distributions to represent a unique combination of a predator and two prey species. Parameter ranges below were chosen to result in predator saturation levels 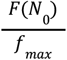 in single-prey experiments at the generic, highest available densities *N*_0_ = 200 of 78% (Holling-2 and Yodzis-FR) and 91% (Holling-3 and Generalized-FR) on average with a standard deviation of 5%, each, emulating a researcher’s choice for a sufficiently large maximum offered abundance.

**First**, we sampled maximum feeding rates *f* _*max,i*_ ∼*U*(50, 100) and half-saturation densities *N*_*half,i*_ ∼*U*(25, 75) of each species (*i* = 1, 2) from uniform distributions. This determines handling times 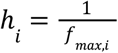 and attack rates 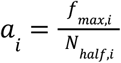 (Holling-2 and Yodzis-FR), or attack coefficients 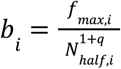 (Holling-3 and Generalized-FR), see e.g. (Rosenbaum & Rall 2018). Exponents *q* (Holling-3 and Generalized-FR) and *r* (Yodzis-FR and Generalized-FR) were also drawn from uniform distributions *U*(0. 5, 1. 5). Their means correspond to linear density-dependences of attack rates (*q* = 1, classic Holling type 3 FR with fixed exponent) and preferences (*r* = 1). Preference weights (*w*_1_, *w*_2_) were sampled from a joint Dirichlet distribution satisfying *w*_1_ + *w*_2_ = 1, with its shape parameters (3, 3) describing the level of concentration towards to their mean (0. 5, 0. 5).

**Second**, for each feeding trial *j* = 1,…, *n* of a given experimental design, we deterministically predicted the mean density of eaten prey 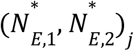. We numerically solved the coupled differential equations (eq. 1) with initial densities (*N*_0,1_, *N* _0,2_)_*j*_ after a generic experimental duration of *T* = 1. 0 using the R-package “deSolve” (Soetaert, Petzoldt & Setzer 2010) and computed 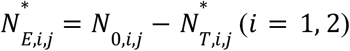.

**Third**, because feeding events happen randomly, we allowed for random variation around the predicted means 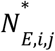 by drawing the final integer numbers of eaten prey items *N* _*E,i,j*_ from Binomial distributions with *N*_0,*i,j*_ trials and success probability 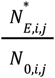. Here, *N*_*E,i,j*_ are bounded between 0 and *N*_0,*i,j*_ .

With this level of randomness we observed coefficients of variation (CV, standard deviation divided by mean) of 0.11 on average (95% interval [0.08,0.15]) in simulated single-prey experiments in highest available densities *N*_0,*i*_ = 200 (Fig. 3). However, variation in experimental datasets can be higher. A systematic investigation of 1,999 lab experiments from the FoRAGE database (Uiterwaal *et al*. 2022) revealed CVs with an average of 0.25 [0.05,0.92] (Fig. S13).

**Figure 3:**
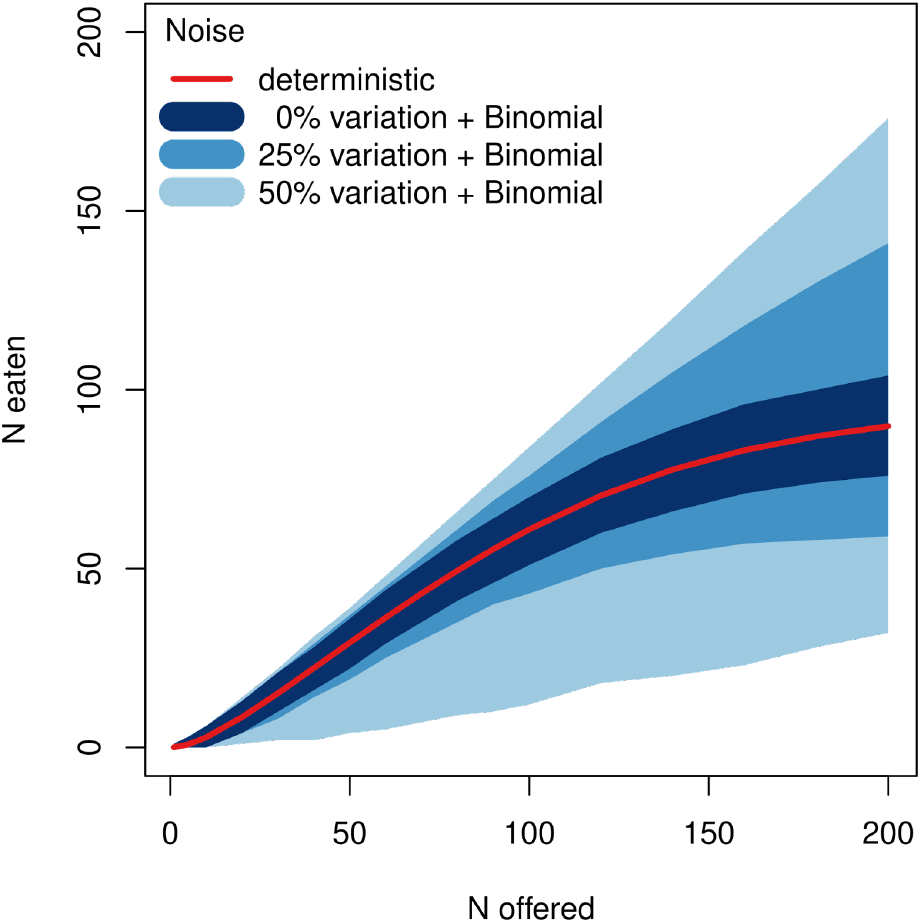
95% intervals of simulated data for a single-species Holling-3 FR (*b* = 0. 04, *h* = 0. 01, *q* = 1. 0) depending on the level of predator trait variation.

To increase the noise in simulated datasets to more realistic levels, we acknowledge that each study usually includes multiple predator individuals that may differ in their morphological and physiological traits, e.g. body size (DeLong 2021; DeLong, Uiterwaal & Dell 2021). We assumed that each trial (*j* = 1,…, *n*) is conducted with one predator individual, which is not reused in multiple trials. In the approach described above, all predators’ traits were identical to the population-level means as described by parameters such as attack rates *a*_*i*_ and handling times *h*_*i*_ (for prey species *i* = 1, 2). Then, we allowed intraspecific trait variation around the population-level means for attack rates *a*_*i,j*_ ∼*N*(*a*_*i*_, *σa*_*i*_) (Holling-2 and Yodzis-FR; same for *b*_*i,j*_, Holling-3 and Generalized-FR) and handling times *h*_*i,j*_ ∼*N*(*h*_*i*_, *σh*_*i*_), where *σ* describes the relative amount of variation. We used *σ* = 0. 05, *σ* = 0. 25 and *σ* = 0. 50, which produced average CVs of 0.12 [0.09,0.15], 0.24 [0.21,0.28] and 0.51 [0.44,0.61] (Fig. S13). Here, noise originates both from variability in traits between individuals (and therefore between trials) and from additional Binomial error due to the randomness of feeding events in each trial.

### Statistical approach

We combined numerical ODE simulations with MCMC to estimate model parameters θ (e.g., θ = (*a*_1_, *a*_2_, *h*_1_, *h*_2_) for Holling-2) from datasets of initial prey abundances (*N*_0,1_, *N*_0,2_)_*j*_ and observed numbers of eaten prey (*N* _*E*,1_, *N*_*E*,2_)_*j*_, *j* = 1,…, *n*. We used the R-package “rstan” (Stan Development Team 2023a) to sample from the posterior probability distribution *P*(θ|*Y*)∼*P*(*Y*|θ)*P*(θ) of model parameters θ given the data *Y*, where *P*(*Y*|θ) is the likelihood function connecting model predictions with data, and *P*(θ) is a prior distribution for the parameters. In each MCMC iteration, model predictions for eaten prey 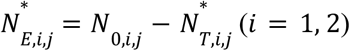 are computed by numerical integration of the coupled equations (1) in *t* ∈ [0, *T*] with initial densities (*N*_0,1_, *N*_0,2_) _*j*_ using Stan’s built-in ODE solver. We chose a Binomial distribution 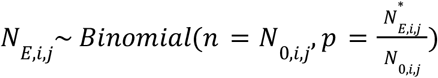 as likelihood function.

As prior distributions we chose exponential distributions with rates 0.5 for *a*_*i*_, 10.0 for *b*_*i*_, 20.0 for *h*, and 1.0 for *q* and *r*, with mean and standard deviation given by the inverse of the rate. Preference weights (*w*_1_, *w*_2_) were coded as two-dimensional simplex vectors with a Dirichlet distribution and its shape parameters (2, 2) as prior distribution to ensure *w*_1_ + *w*_2_ = 1. These vaguely informative priors merely represent the parameters’ order of magnitude, as not to confound results on estimation accuracy with prior information (Fig S14). We note that prior distributions are system-specific and uninformative priors for one experiment might be highly informative in other experiments. Furthermore, the importance of prior information depends on the sample size (Wolf, Novak & Gitelman 2017) and can be tested with prior predictive checks (Wesner & Pomeranz 2021; Stan Development Team 2023b).

Each model was fitted using 1,000 warmup steps and 2,000 samples in three chains, adding up to 6,000 samples of the parameters’ posterior distribution. We discarded datasets when the MCMC did not converge properly (more than 100 divergent iterations).

### Evaluation

To evaluate the accuracy of each model fit we compared metrics of parameter estimates 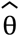, given by their posterior probability distribution 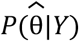, and the known true parameters θ which generated the dataset (Morris, White & Crowther 2019). Relative bias

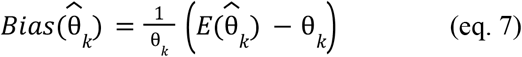

is a measure of how much a parameter’s point estimate, posterior mean 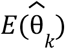, deviates from the true value (accuracy). Relative root mean squared error

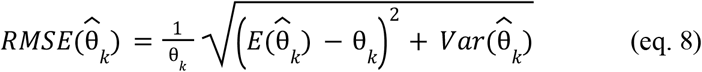

also accounts for the uncertainty of the estimate, expressed by posterior variance 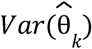 (precision). To summarize this information for each fit, single metrics were computed through geometric means of the individual parameters’ RMSEs

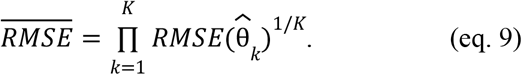

### Model comparison

We tested whether not only true model parameters could be accurately recovered by fitting the same MSFR model that generated the dataset, but also whether the correct model could be identified among other model candidates. For the data from designs that produced the most accurate estimates as described in the previous section (log-spaced grid), we additionally fitted all four model candidates to each dataset and performed model selection. We used the leave-one-out cross-validation (LOO-CV) method, which estimates a fitted model’s out-of-sample predictive accuracy. It can be used similarly to AIC, but it uses the parameters’ full uncertainty and does not rely on normality assumptions for the likelihood (Gelman, Hwang & Vehtari 2014). It is computed via the R-package “loo” (Vehtari *et al*. 2023). The model associated with the lowest information criterion was selected as the best performing model. However, if the difference in model performance was less than two standard deviations of the included uncertainty, the less complex model was selected (ranking Holling-2 < Holling-3 < Yodzis-FR < Generalized-FR in complexity).

### Three-prey scenarios

Additionally, we tested scenarios with a predator feeding on three prey species. All of the above methods can be easily extended to models, simulations and fittings with more than two prey species. We wanted to know if feeding trials involving all three prey species simultaneously are required to accurately estimate model parameters, or if pairwise combinations of two-prey feeding trials are sufficient. The log-spaced grid design, which performed best in the study described above, was extended to

1. a full factorial three-dimensional 7×7×7 grid
2. three 8×8 grids with species combinations (1,2), (1,3) and (2,3),

see Fig. S17. Subtracting single-prey trials that would occur twice, the pairwise trials sum up to 3×8^2^–3×8=168 unique combinations of prey abundances, which were replicated 1, 2, 4 and 6 times to generate sample sizes *n* of 168, 336, 672 and 1008. The full factorial trials total 7^3^=343 unique combinations, which were replicated 1, 1, 2 and 3 times (343, 343, 686, 1029) and randomly downsampled to the four sample sizes of the pairwise designs, respectively, for comparability. Data was generated for all four MSFR models and all three noise levels. All parameter ranges were as described above in the two-prey scenarios. Again, 500 replicates of each scenario were generated and fitted, summing up to 48,000 datasets.

### Empirical datasets

We analyzed 23 datasets from 10 publications, including 5 studies from the list of (Stouffer & Novak 2021) and 5 recent studies, see Table S1 for a full list of datasets. We required datasets to include both single- and two-prey experiments, and some variation in offered relative prey abundances *N*_0,1_/*N*_0,2_. For two studies that did not include raw data but summary statistics of eaten prey, we replicated full datasets. Numbers of individual feeding trials ranged from 48 to 290. Experiments had been conducted either without or with prey replacement, and two studies also included a mixture of both. In case of prey replacement, feeding rates were assumed to be constant during the whole experiment. We fitted direct predictions *N* _*E,i*_= *F*_*i*_ (*N*_0,1_, *N* _0,2_) · *T* (*i* = 1, 2) instead of ODE predictions, and used a Poisson likelihood instead of a Binomial likelihood function. Bayesian modeling allowed these two types of observations to be combined in one statistical model for the two studies with mixed data. We performed model selection as described above to determine the best MSFR for each dataset.

## Results

We observed good convergence of the MCMC algorithm when fitting simulated data (100% of all fits for Holling-2 and Holling-3, 99% Yodzis-FR, 97% Generalized-FR). Runtime per model fit ranged between a couple of minutes for the smallest and several hours for the largest datasets.

### Experimental design

We first assessed through which experimental design we could obtain the most accurate parameter estimates. Overall, experiments with the log-spaced grid resulted in smallest estimation errors in terms of RMSE for all four MSFRs and all levels of sample size and noise. Fig. 4 shows results for intermediate noise scenarios, but similar trends were observed in all three noise scenarios (Figs. S5-S7). In general, estimation performance increased with sample size as expected. But the choice of design was so influential that it could outperform other designs with twice the amount of feeding trials, e.g. log-spaced grid vs. a random or a space-filling design for Yodzis-FR and the Generalized-FR (Fig. 4c,d). For these two active switching MSFRs, the difference between the log-spaced grid and other designs was also the most pronounced.

**Figure 4:**
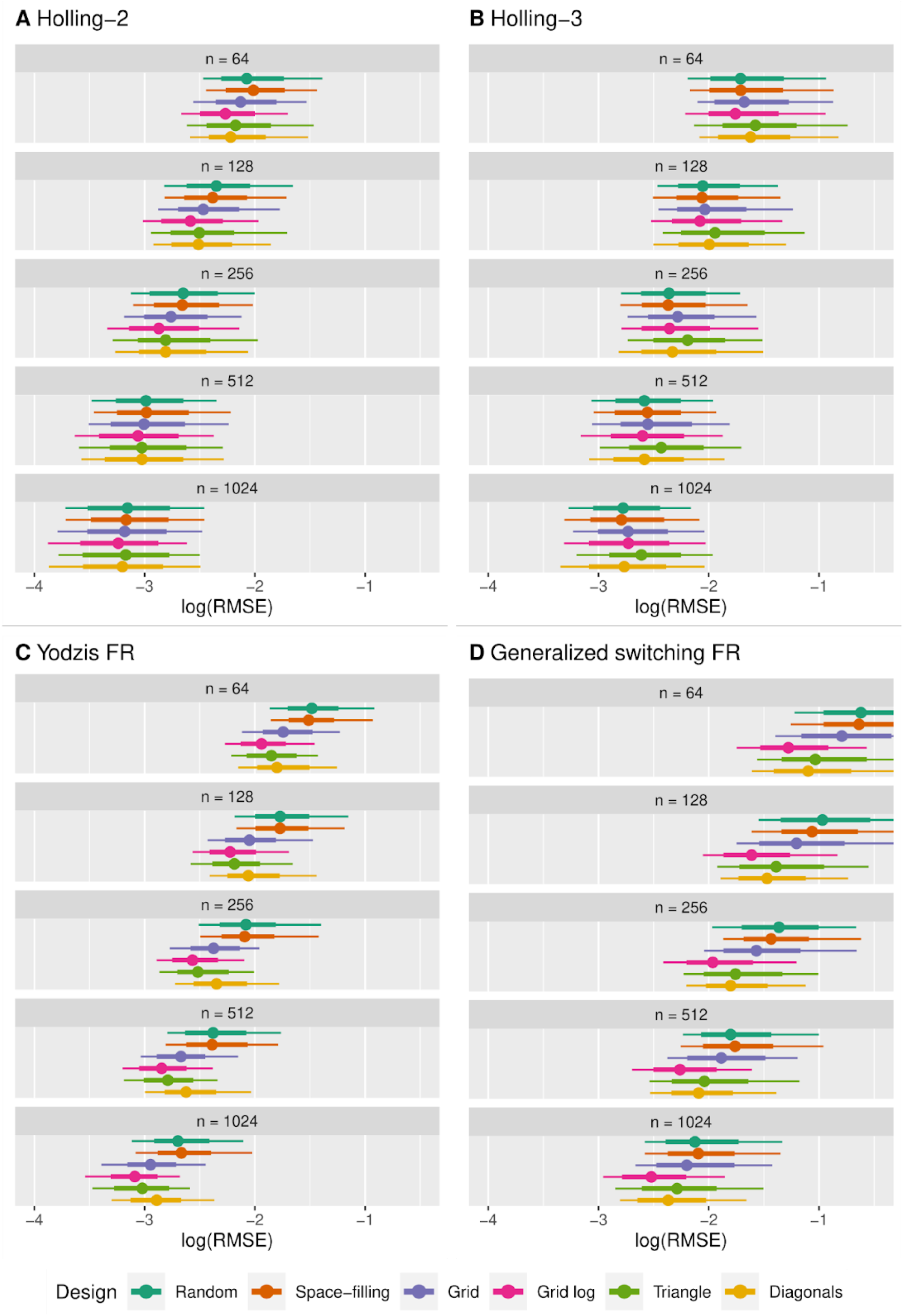
Distributions of parameters’ relative RMSE (eq. 9) vs. experimental design and sample size, noise level: medium. 500 datasets each. Dots represent means, bold lines are 66% intervals, thin lines are 95% intervals.

In many scenarios, the random and the space-filling designs performed worst. These were the only designs that do not explicitly include single-prey experiments (except by random chance), suggesting that experiments with mixed two-prey feeding trials only should be avoided.

The triangle design gave mixed results, e.g. performing second best for Yodzis-FR but worst for Holling-3. Such designs are common in the literature (Stouffer & Novak 2021). They are suitable for analyzing two-prey feeding data with generalized linear models (GLMs) by relating the proportion of species 1 consumed 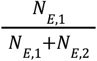 to the proportion of species 1 available 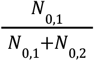, while the total amount of offered prey *N*_0,1_ + *N*_0,2_ is constant for the mixed-prey scenarios, e.g. (McCard *et al*. 2021). However, these fixed-sum feeding trials are not necessarily representative of all combinations of prey abundance (Fig. S15).

From here on, all results will be presented for the best performing design, log-spaced grid, only.

### Intraspecific variation effects

Second, we tested how noise (caused by the predator’s intraspecific trait variation) affected estimation performance (Fig. 5). Estimates generally worsened with increasing noise as expected. They improved with sample size in low- and medium-noise scenarios. In high-noise scenarios, however, estimates did not improve much on average with increasing sample size after a certain threshold was reached, e.g. comparing the performance of datasets with 512 and 1024 feeding trials.

**Figure 5:**
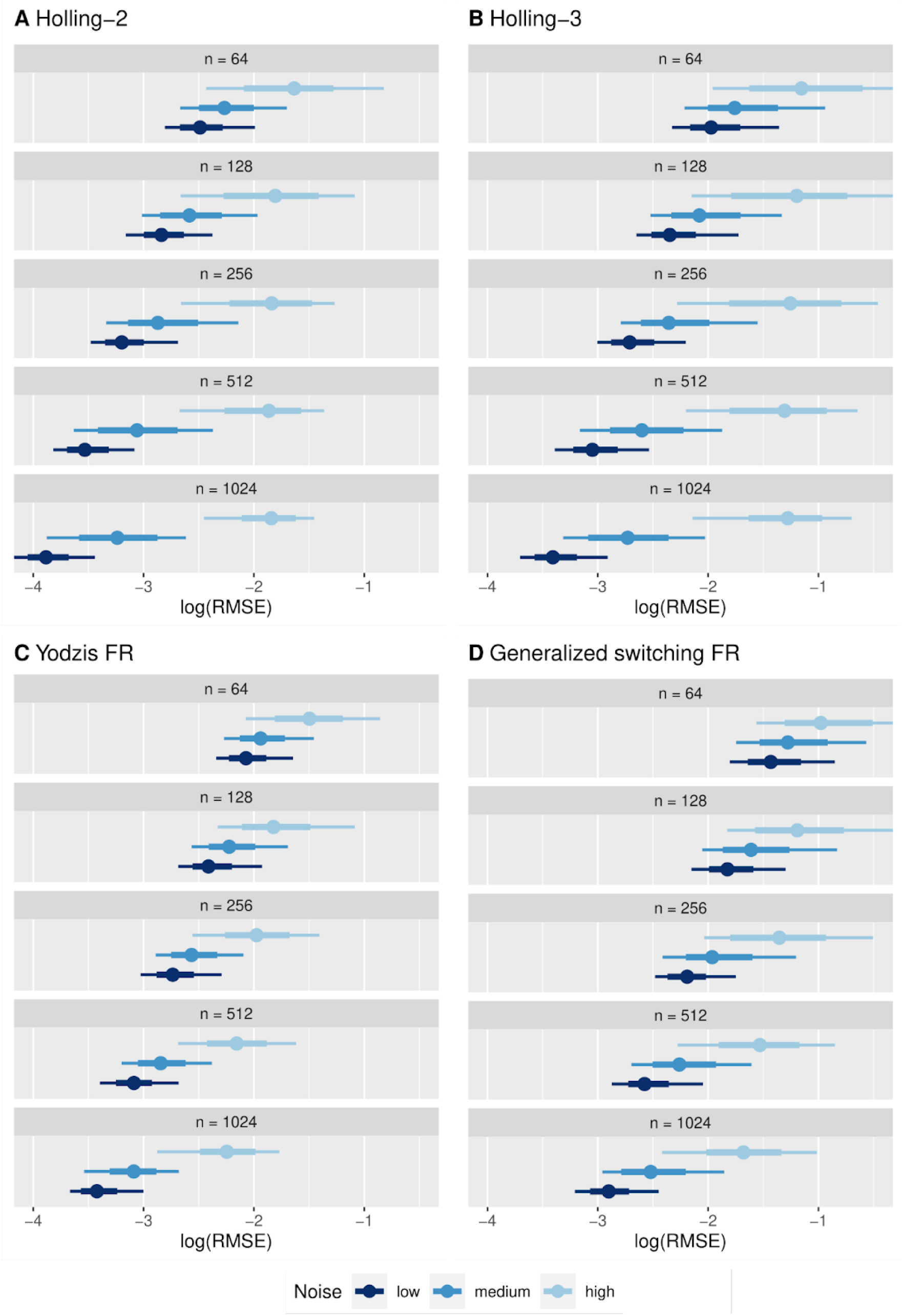
Distributions of parameters’ relative RMSE (eq. 9) vs. noise and sample size, design: log-spaced grid. 500 datasets each. Dots represent means, bold lines are 66% intervals, thin lines are 95% intervals.

This is because most parameters’ estimates are systematically biased in these high-noise datasets (Figs. S1-S4). Handling times were, on average, underestimated for all four MSFR models. Attack rates were also underestimated on average (Holling-2 and Yodzis-FR). For Holling-3 and the Generalized-FR, attack coefficients were overestimated, while their exponents were underestimated. Negative bias in exponents generally lead to underestimated attack rates *a* = *bN*^*q*^, too.

These systematic biases can be traced back to Jensen’s inequality for nonlinear functions: Variation in individual attack rates lowers mean feeding (mostly at low to intermediate abundances) and variation in handling times increases mean feeding (at high abundances), see (Bolnick *et al*. 2011) and Fig. S16. Fitting population-level mean parameters must necessarily compensate for this by underestimating feeding rates at low abundances (by underestimating attack rates or attack exponents) and overestimating feeding rates at high abundances (by underestimating handling times).

Additionally, we tested whether alternative likelihood functions that account for overdispersion in the data could mediate these biases. We used the Negative Binomial and the Beta Binomial distribution (Fenlon & Faddy 2006). Although these distributions account for higher variance in the data, their estimation performance was even worse than that of the Binomial distribution (Fig. S8). They all fit population-level mean feeding rates through population-level mean trait parameters, which are not identical. We hypothesize that overdispersion parameters introduce additional degrees of freedom, which may worsen estimation performance (see also (Novak & Stouffer 2021b)) while not solving the problem of Jensen’s inequality.

### Model comparison

Model selection was almost always able to identify the model used to simulate the data from the four candidate models in low to medium noise scenarios (Fig. S12). Importantly, small datasets (n=64) were sufficient to select the correct model in more than 99% of the cases in total. Although true parameter values could mostly not be identified without bias in high-noise scenarios, model selection showed an equally good performance for Holling-2 and Holling-3. For Yodzis-FR and the Generalized-FR sample sizes of n=128 and n=256, respectively, were required to reach an equivalent level of certainty.

### Three-prey scenarios

Estimation errors (RMSE) generally decreased with sample size for all four MSFRs in both full factorial and pairwise experimental designs (see Fig. 6 for medium noise, Figs. S9-S11 for all levels). As in the two-prey scenarios, noise in the data increased bias, and in the high noise scenarios improvement with sample size was only small (Fig. S11). Parameter estimates were generally better when using the pairwise experimental design for Holling-2, Yodzis-FR and the Generalized-FR, while they were slightly worse than the full factorial design for Holling-3. At least for our four MSFRs, we here support the use of pairwise experiments, which may be logistically more feasible than feeding trials involving three prey species.

**Figure 6:**
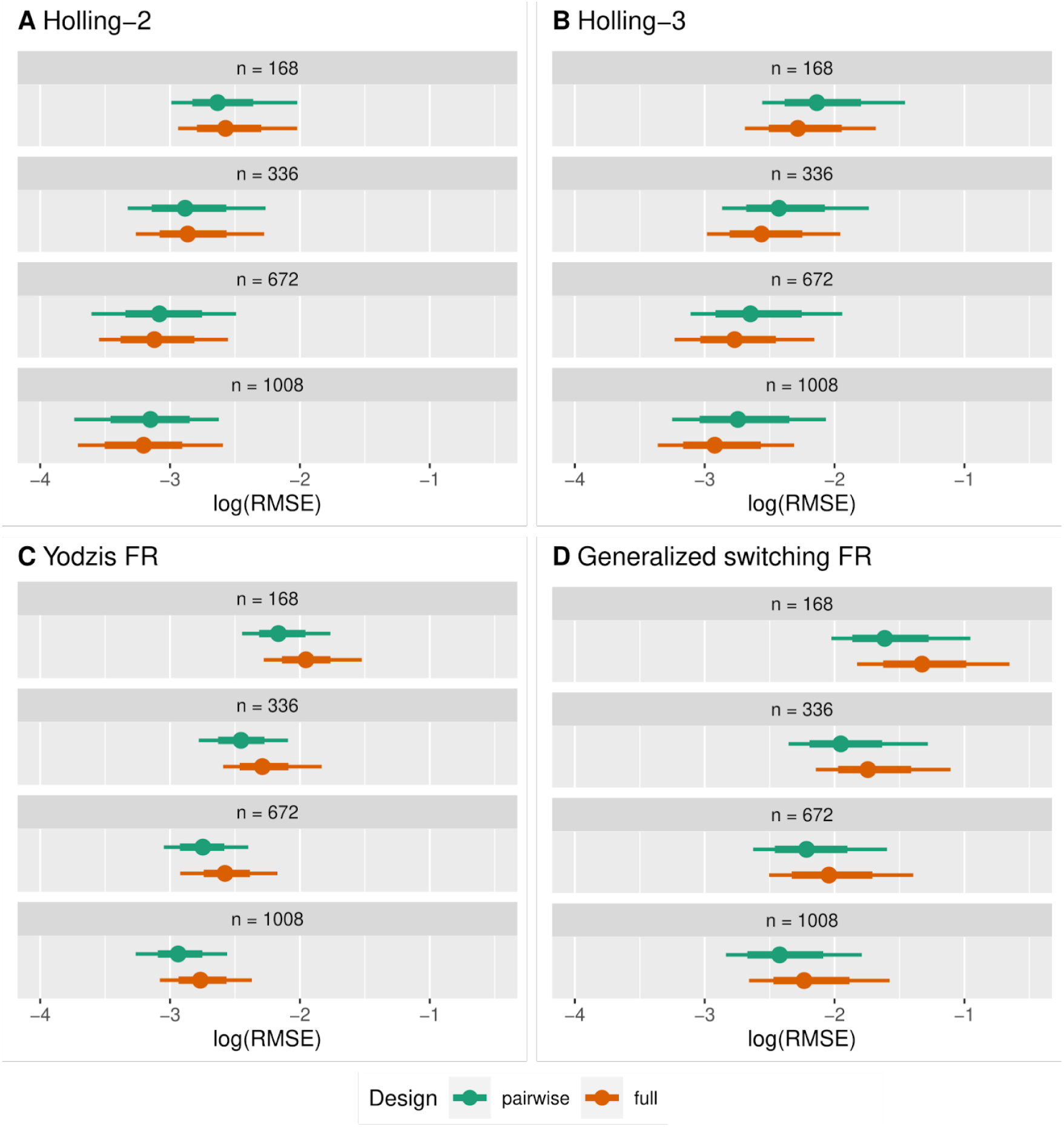
Three-prey scenarios. Distributions of parameters’ relative RMSE (eq. 9) vs. experimental design and sample size, noise level: medium. 500 datasets each. Dots represent means, bold lines are 66% intervals, thin lines are 95% intervals.

### Empirical datasets

Of the 23 datasets the Holling-2 MSFR was the best-performing model 15 times, followed by Holling-3 (5 times) and Yodzis-FR (3 times), see Tables S1, S2 and [Zenodo]. Switching was observed in 5 datasets, i.e. the relative proportion of eaten prey switched from avoidance to preference (see e.g. Fig. 7c). Interestingly, all of these 5 datasets were best described by the Holling-3. All 3 of the best-performing Yodzis-FRs did not include switching, but multi-prey consumption differed from

**Figure 7:**
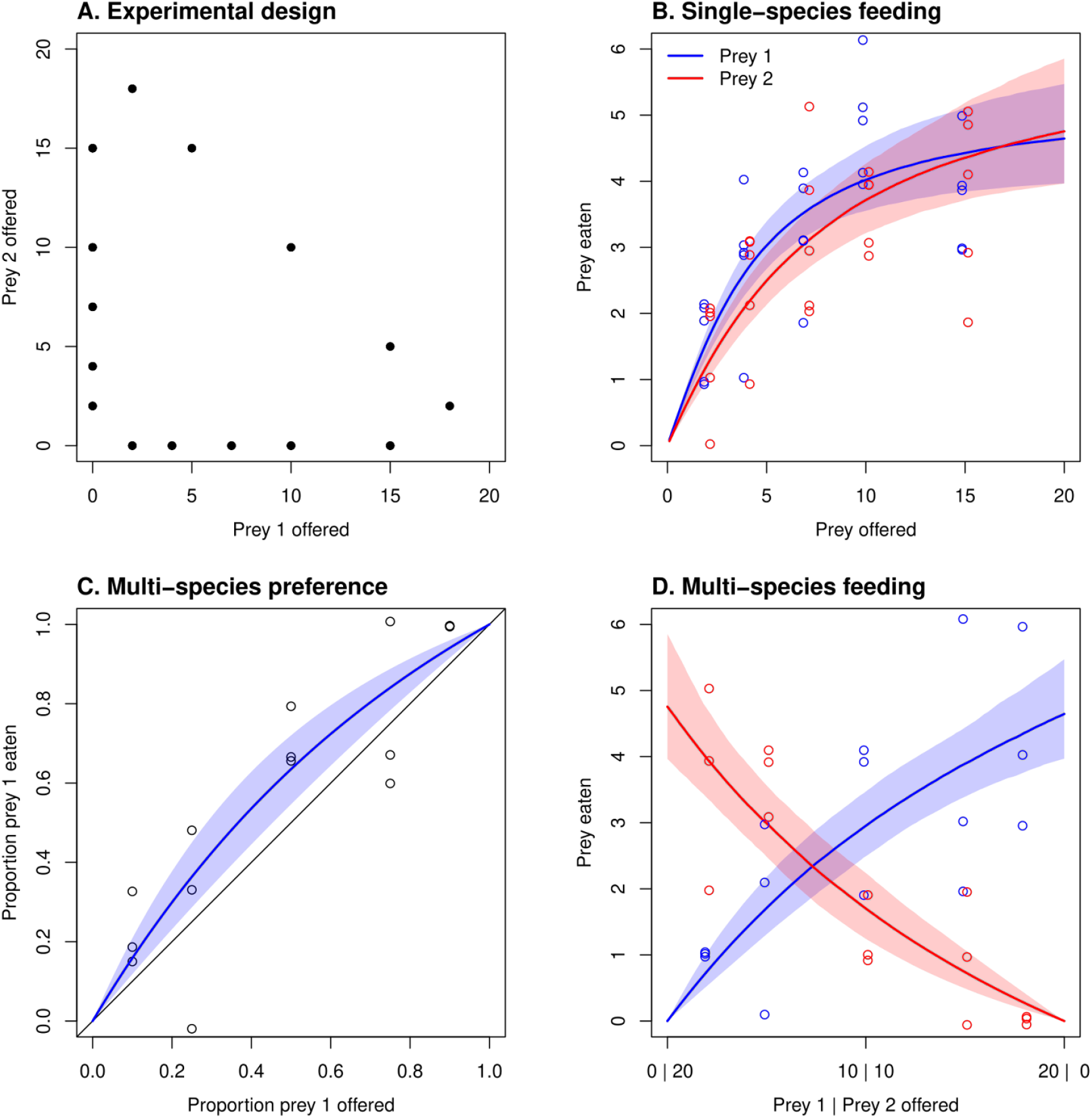
Best-fitting model Holling-2 for empirical data from (Cuthbert *et al*. 2019). (A) Experimental design (“Triangle”) with 5 replicates per single-species and 3 replicates per multi-species combination, each. (B) Single-species observed and predicted eaten prey (line is median, shaded areas 90% credible intervals). Multi-species observed and predicted for (C) proportion of prey 1 eaten 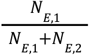 vs. proportion of prey 1 offered 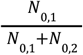, and (D) eaten prey along the diagonal (panel A top-left to bottom-right, *N* _0,1_+ *N*_0,2_ = 20).

single-prey consumption through preferences *w*_1_ ≠ *w*_2_ of one species over the other. None of the datasets were best described by our new Generalized-FR, meaning it did not show a difference in model fit over another MSFR of more than 2 standard deviations (associated uncertainty). However, in 9 cases it performed as well as the best model under uncertainty, but a less complex model was chosen. This was also the case for Yodzis-FR (8 times) and Holling-3 (13 times).

In 5 datasets, where experimental duration differed substantially between single- and multi-prey feeding trials (e.g., 24h vs. 3h), none of the models could fit all observed data sufficiently. Observed feeding rates (averaged by time) were systematically lower in long than in short experiments, as reported by (Li, Rall & Kalinkat 2018). Thus, modeled feeding rates tended to systematically overpredict observed feeding in the longer single-prey trials and underpredict in shorter multi-prey trials, even though our fitting method explicitly accounted for the individual duration of each trial. We assume that experimental conditions were not constant (e.g. predator hunger levels, or additional resting times) and fitting models that do not account for this to data with substantial variation in experimental duration could lead to biased parameter estimates.

We here showcase the analysis of one dataset from (Cuthbert, Callaghan & Dick 2019), with a native cyclopoid copepod feeding on the invasive Asian tiger mosquito (prey 1) and a representative native common house mosquito (prey 2). They used a “triangle” design and conducted 65 feeding trials (6 hours without prey replacement, each). For the single-species experiments, they reported type 2 functional responses. We also identified Holling-2 as the best-performing MSFR (Fig. 7). The predator shows a preference for prey 1 due to differences in attack rates (posterior mean *a*_1_ − *a*_2_ = 1. 11 with a 95% credible interval [0.23,2.31]), but not in handling times (*h*_1_ − *h*_2_ = 0. 03 [-0.05,0.11]). It was pointed out in the original study that the predator’s preference might mitigate the invasive species’ superior competitive ability and facilitate coexistence with native prey (as has been observed in nature, but see (Landi, McCoy & Vonesh 2022) for a discussion). With our fully parameterized MSFR, these mechanisms could be explored more thoroughly by modeling community dynamics.

## Discussion

We explored previously neglected topics in empirical MSFR research: experimental design and prey depletion. We integrated previous approaches into a flexible framework for parameterizing MSFR models from both single-prey and multi-prey feeding trials simultaneously and demonstrated its application thoroughly. Using ODE simulations to compute model predictions allows correcting for prey depletion during the course of the experiments, which is necessary if eaten prey are not constantly replaced (and would involve an extensive logistical effort). Although not tested here, ODE simulations could be easily extended to include processes such as prey mortality or growth if required (Rosenbaum & Rall 2018; Daugaard *et al*. 2019). If, however, experiments are conducted with prey replacement, they can be fitted through direct predictions from MSFR models (without ODE simulation). A Bayesian approach even allows using mixed datasets from experiments with and without replacement, as demonstrated here in empirical datasets and the tutorial. While fitting these models could theoretically be achieved with, e.g., maximum likelihood approaches, too, Bayesian inference quantifies uncertainty in parameters estimates more accurately and also uncertainty in model comparison (LOO-CV) is accounted for through posterior distribution samples (Gelman *et al*. 2014).

We comprehensively tested the identifiability of four MSFR models with realistically simulated feeding trials for different noise levels, numbers of feeding trials, and experimental designs. As practical advice, we recommend a log-spaced grid design for experimental planning, which distributes more feeding trials at low abundances of individual prey species and fewer at high total prey abundances, and also multi-prey trials at different levels of total prey abundances. It produced the most accurate parameter estimates across all models, sample sizes, and noise levels. This is an extension of results from (Uszko *et al*. 2020; Novak & Stouffer 2021a), who found that log-spaced distances of abundances were more effective than equidistant designs in single-prey experiments. Additionally, for three-prey systems, we found that pairwise trials with two prey species each (which might be logistically less demanding) produce more accurate estimates than trials with all species present, at least for our four MSFRs without higher-order interactions. Recently, sequential designs have been proposed to reduce the total number of feeding trials in single-prey experiments (Moffat *et al*. 2020), and their extension to the multi-prey context has potential to decrease experimental effort even more.

Classically, two-prey experiments have often been conducted using a ‘triangle’ design with single-prey trials for each species and multi-prey trials along a fixed number of total prey offered but varying prey composition (Table S1). Prey switching was then investigated via GLMs, but these studies were not able to parameterize a full MSFR model. With parameterized MSFRs, switching can be detected through model comparison and visualization of model predictions, and the best-performing model can be used for dynamical simulation of species communities and hypothesis testing, e.g. effects of allometry or temperature (Kalinkat *et al*. 2011; Gauzens *et al*. 2024).

Although we have only discussed four MSFR models here, these cover important properties such as no, passive and active switching in multi-prey scenarios, as well as type 2 and type 3 behavior in single-prey scenarios. Furthermore, our proposed Generalized-FR represents a flexible but tractable model that generalizes the three other models by using focal species density-dependent attack rates *a*_*i*_ (*N*_*i*_) and community density-dependent selection factors *p*_*i*_ (*N*_1_ … *N*_*m*_) (Koen-Alonso & Yodzis 2005). It also includes the three MSFRs when respective exponents *r* or *q*, or both, are set to zero.

While a plethora of models have been proposed (Gentleman *et al*. 2003; Morozov & Petrovskii 2013; Baudrot *et al*. 2016; Lehtinen *et al*. 2024), to our knowledge this is the first MSFR that combines sigmoidal behavior in single-prey scenarios, active switching in multi-prey scenarios, and species-specific handling times. An MSFR with the first two properties was proposed by (Baudrot *et al*. 2016), but an extension of their model H3.2 to include individual handling times in their supplement contained an error (see SI for a correction). As the strength of switching (expressed through MSFR exponents) can affect stability and coexistence (Archibald *et al*. 2023), active switching models such as Yodzis-FR or Generalized-FR should be considered and tested against data. This often requires allowing prey species-specific maximum feeding rates, and therefore handling times (as demonstrated in empirical datasets here).

All four models can produce feeding that is suboptimal (an increase in one prey species can lead to a decline of total feeding) or antagonistic (total feeding is lower in a mixed-prey scenario than in a single-prey scenario with identical total abundance), either through a difference in handling times or as a result of prey switching (Gentleman *et al*. 2003). (Vallina *et al*. 2014) proposed a phenomenological Kill-the-winner model as a remedy to achieve optimal feeding including prey switching. However, this comes at the cost of identical handling times and synergistic feeding (a counterintuitive pattern, where an increase in alternative prey abundance can increase focal prey feeding) (Archibald *et al*. 2023). In general, it is not clear whether experimentally measured feeding is necessarily optimal (Griffen 2021), and processes that might lead to optimal or suboptimal feeding remain to be empirically tested in the MSFR framework.

Other studies have found evidence against the Holling-2 MSFR in empirical data. In (Stouffer & Novak 2021), a generalized Holling-2 model included parameters describing the effect of alternative prey on the time lost to handling and therefore the feeding rate of a focal species. It outperformed the classical Holling-2 model in terms of AIC in their database of 30 experiments. Also, (Smith & Smith 2020) found a predominance of Holling-3 over Holling-2 in field data of 48 predator size categories of fish. Here, we identified Holling-2 as the best fitting model in 15 out of 23 datasets. However, this choice was mostly conservative, as in 14 datasets other models were ranked equally under model uncertainty and Holling-2 was chosen as the least complex model. We therefore advocate for a hypothesis-driven approach when selecting model candidates for model comparison (e.g., switching vs. non-switching, or optimal vs. suboptimal). While testing dozens of previously published MSFRs is beyond the scope of this paper, this framework and the tutorial should allow researchers to assess the performance of their own models and experimental designs.

There are some general limitations to fitting MSFR data from experiments, which however are not specific to multi-prey data, but also apply to single-prey experiments. Low sample size studies suffer from biased parameter estimates through the nonlinear nature of the likelihood function (Stouffer & Novak 2021). If individual predators’ traits vary strongly, we have shown that parameter estimates are biased even for large sample sizes. Jensen’s inequality, which states in this context that the population-level mean feeding rate 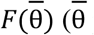 population-level mean trait) is different from the mean of individual feeding rates 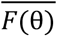 (θ collection of individual trait values), is still an unresolved problem (Bolnick *et al*. 2011). If predator individuals are reused in multiple feeding trials, mixed effects models with predator ID as a random factor could account for intraspecific variation to reduce bias. However, if individual predators are used for each trial (or identity was not recorded in the experiment), this is not feasible, and our study at least quantifies the expected direction and strength of these biases.

Another problem was that some empirical datasets used a wide range of experimental duration up to a factor of 8 (mostly between single-prey and multi-prey trials). Here, a joint model that fits all feeding trials simultaneously may be inappropriate, even if model predictions explicitly include duration. In longer experiments, non-foraging behavior becomes more likely (e.g. reduced hunger levels, resting times) (Griffen 2021), and longer trials produce smaller estimates of attack rates and longer estimates of handling times (Li *et al*. 2018). We therefore suggest the use of identical trial durations when aiming to parameterise a functional response model. (Griffen 2021) argued that it should be long enough to include non-foraging behavior, which could facilitate scaling up functional responses to realistic (field) settings (Novak *et al*. 2017; Uiterwaal & DeLong 2024). If experimentally feasible, entire time series of observed abundances from longer trials could be analyzed with dynamical models (Coblentz & DeLong 2021; Rosenbaum & Fronhofer 2023) and should contain more information than the records of only initial and final abundances.

Filling the gap between experimental evidence for single- and for multi-prey functional responses will be a challenging and lengthy endeavor. We wanted to facilitate this task by equipping researchers with appropriate statistical tools and also practical considerations for experimental planning. We hope that our proposed integration of methods will pave the way for more empirical research on predators feeding on multiple resources, thus improving our understanding of trophic interactions in complex ecosystems.

## Supporting information

Tutorial

## Acknowledgements

We gratefully acknowledge the support of iDiv funded by the German Research Foundation (DFG-FZT 118, 202548816). The scientific results have in part been computed at the High-Performance Computing Cluster EVE of the Helmholtz Centre for Environmental Research (UFZ) and iDiv, and we thank Christian Krause for technical support. We thank Ross Cuthbert for making additional data openly available.

