## Supplement for "Towards understanding interactions in a complex world: Design and analysis of multi-species functional response experiments"

Journal title

### Supporting Information S1

Benjamin Rosenbaum[1,2,*], Jingyi Li[1,2], Myriam R. Hirt[1,2], Remo Ryser[1,2], Ulrich Brose[1,2]

1. EcoNetLab, German Centre for Integrative Biodiversity Research (iDiv), Halle-Jena-Leipzig, Germany
2. Institute of Biodiversity, Friedrich Schiller University Jena, Jena, Germany

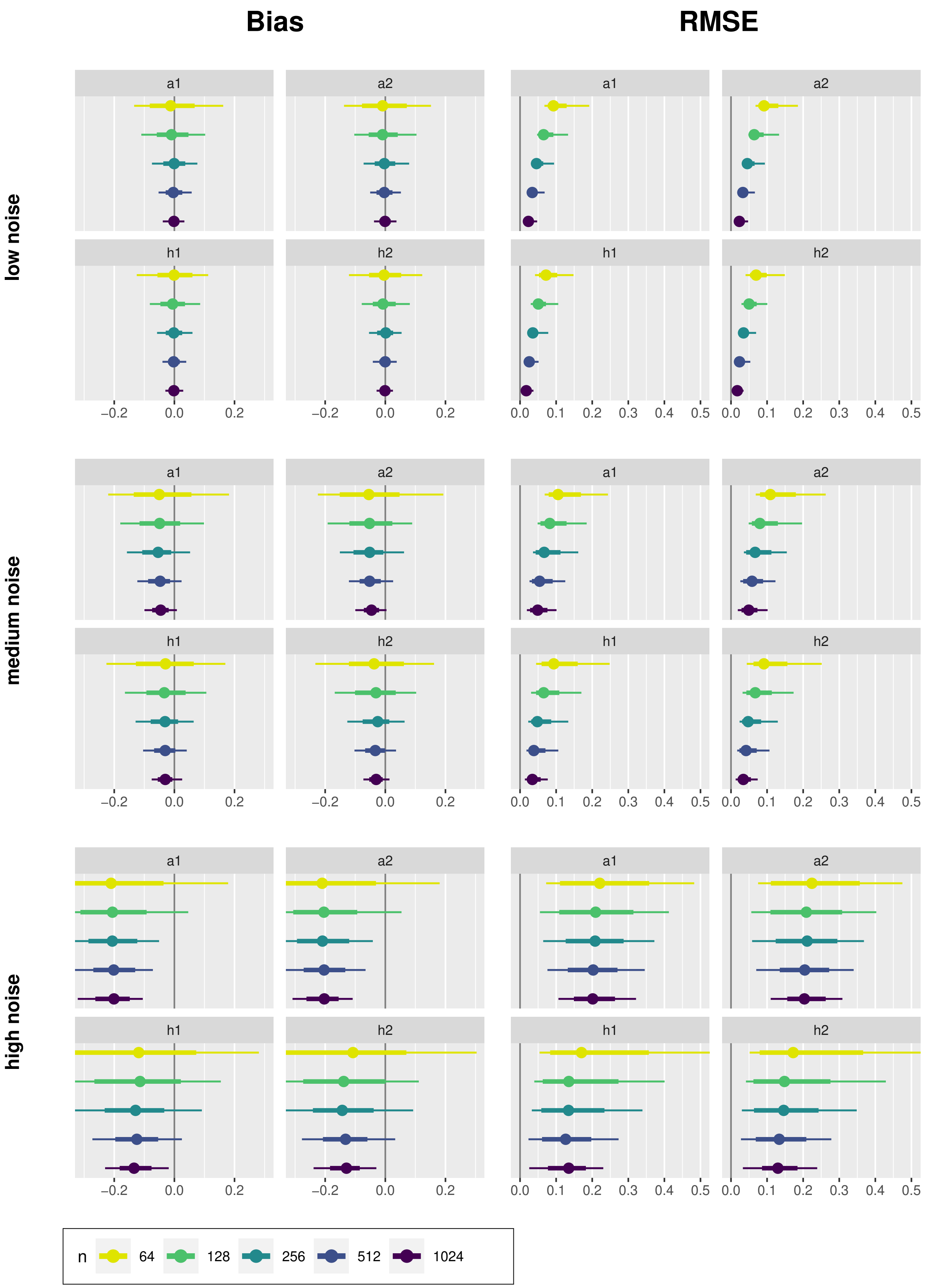


**Figure S1: Holling-2** model. Distribution of each parameter's relative Bias and RMSE vs. noise level and sample size, 500 replicates each. Experimental design: log-spaced grid.

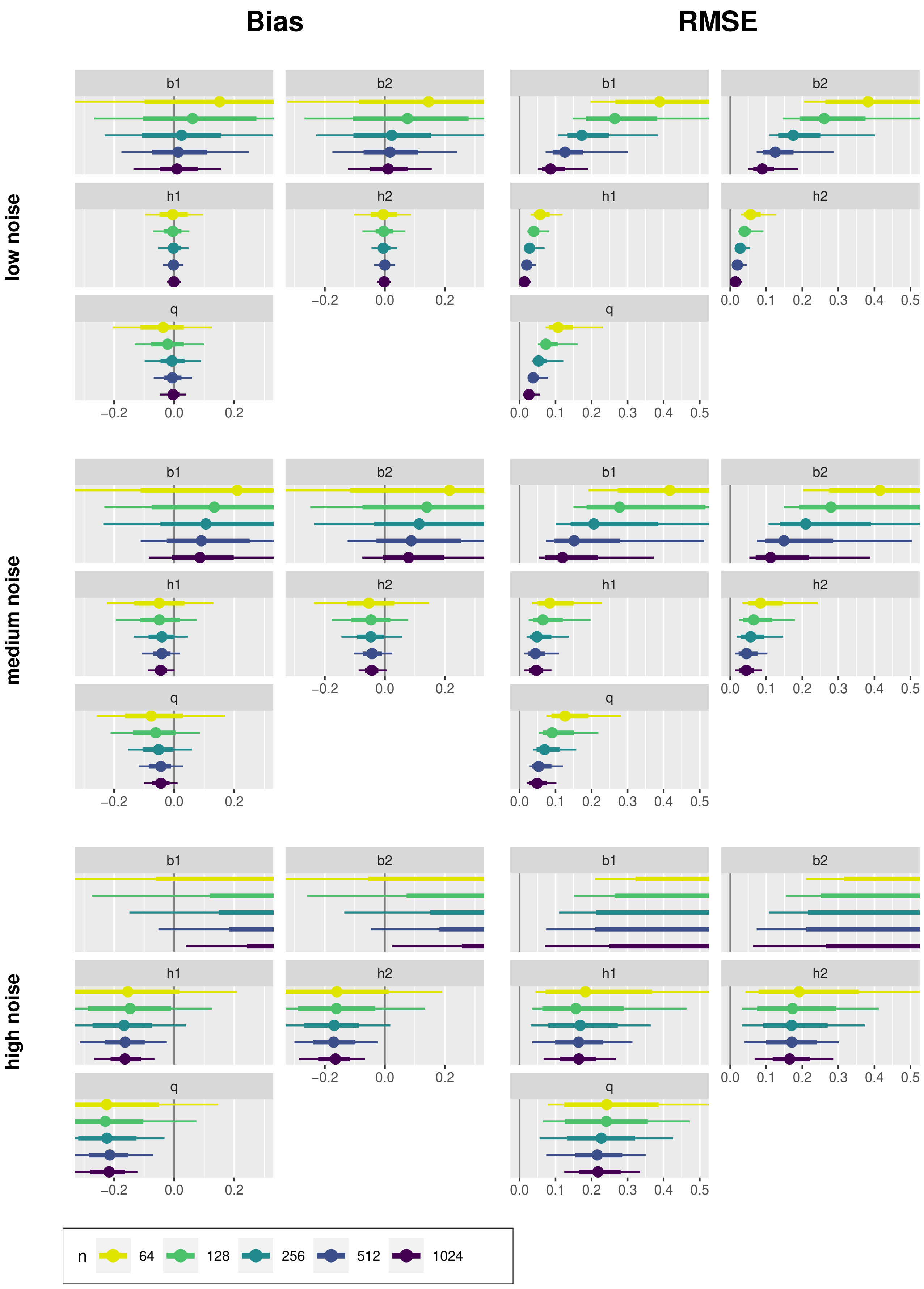


**Figure S2: Holling-3** model. Distribution of each parameter's relative Bias and RMSE vs. noise level and sample size, 500 replicates each. Experimental design: log-spaced grid.

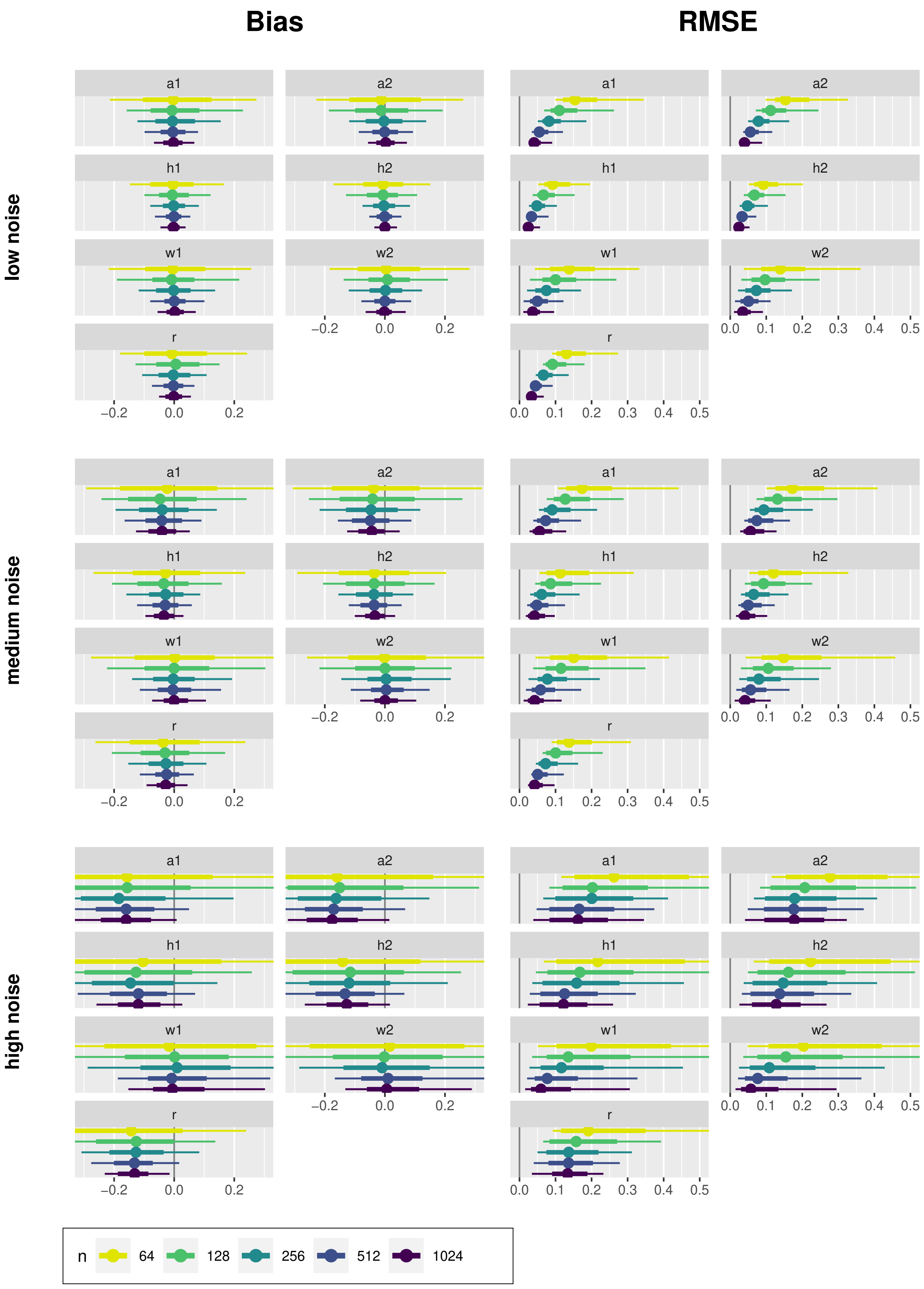


**Figure S3: Yodzis FR** model. Distribution of each parameter's relative Bias and RMSE vs. noise level and sample size, 500 replicates each. Experimental design: log-spaced grid.

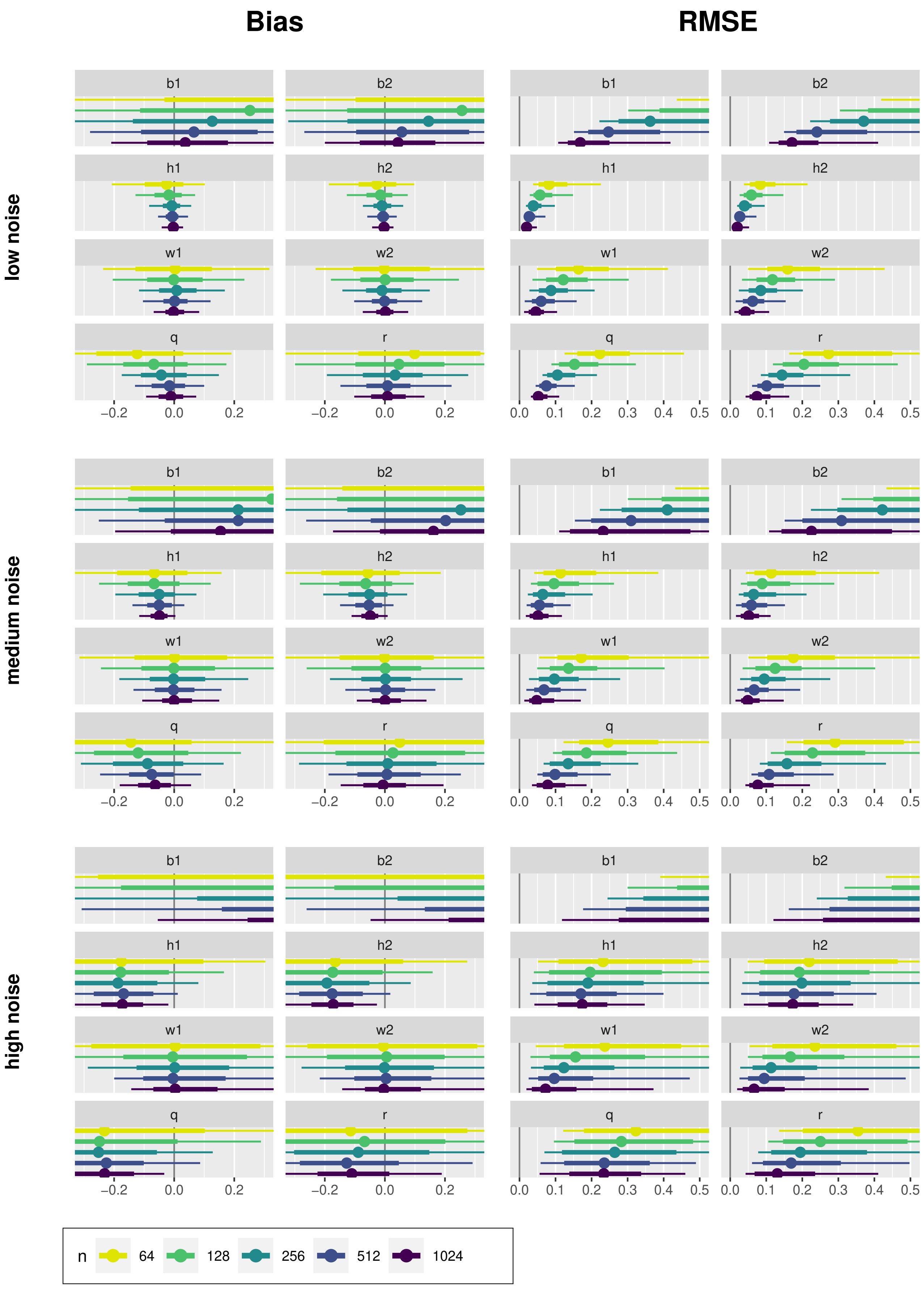


**Figure S4: Generalized switching FR** model. Distribution of each parameter's relative Bias and RMSE vs. noise level and sample size, 500 replicates each. Experimental design: log-spaced grid.

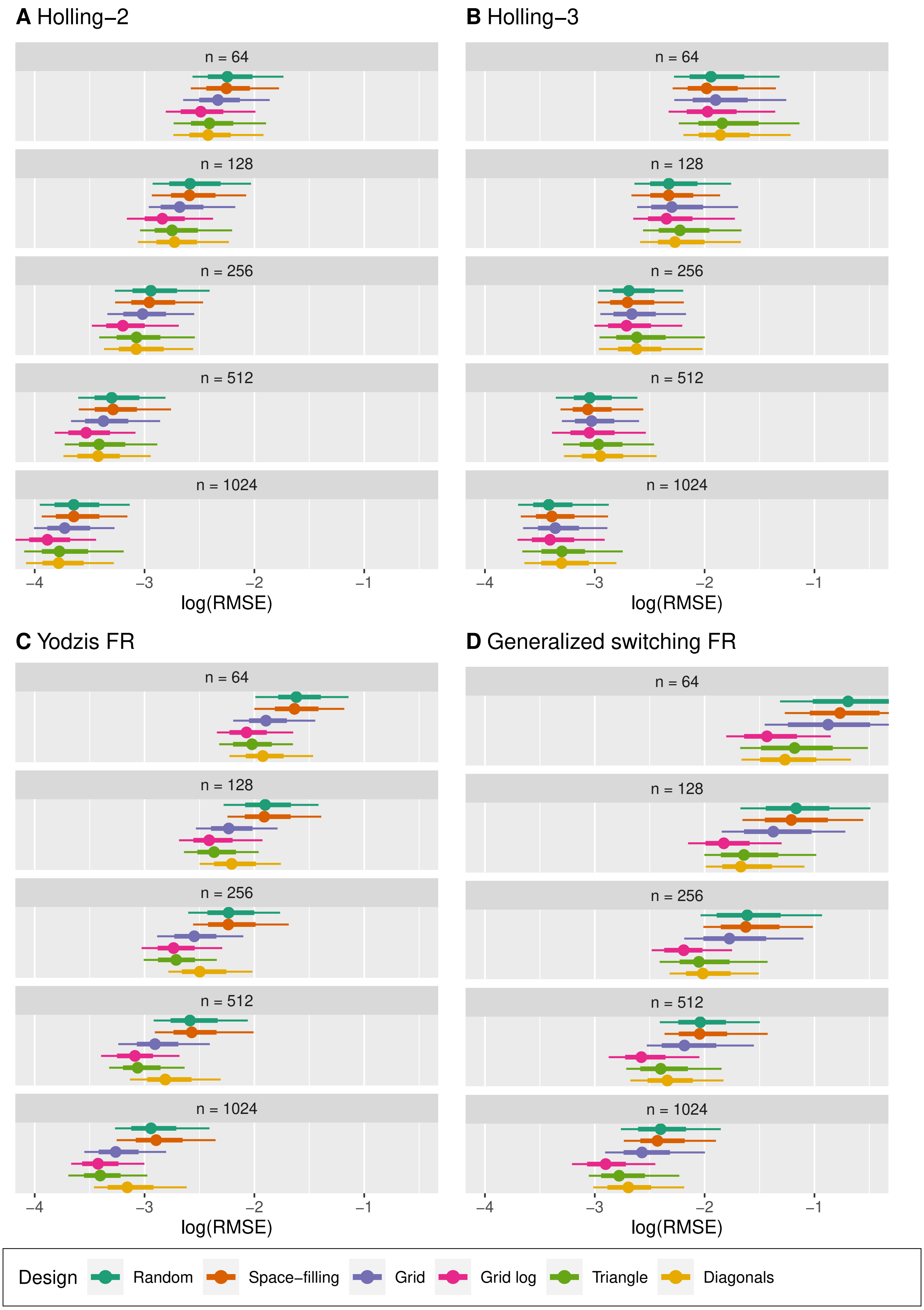


**Figure S5:** Distributions of relative RMSE vs. experimental design and sample size, **noise level: low**. 500 datasets each. Dots represent means, bold lines are 66% intervals, thin lines are 95% intervals.

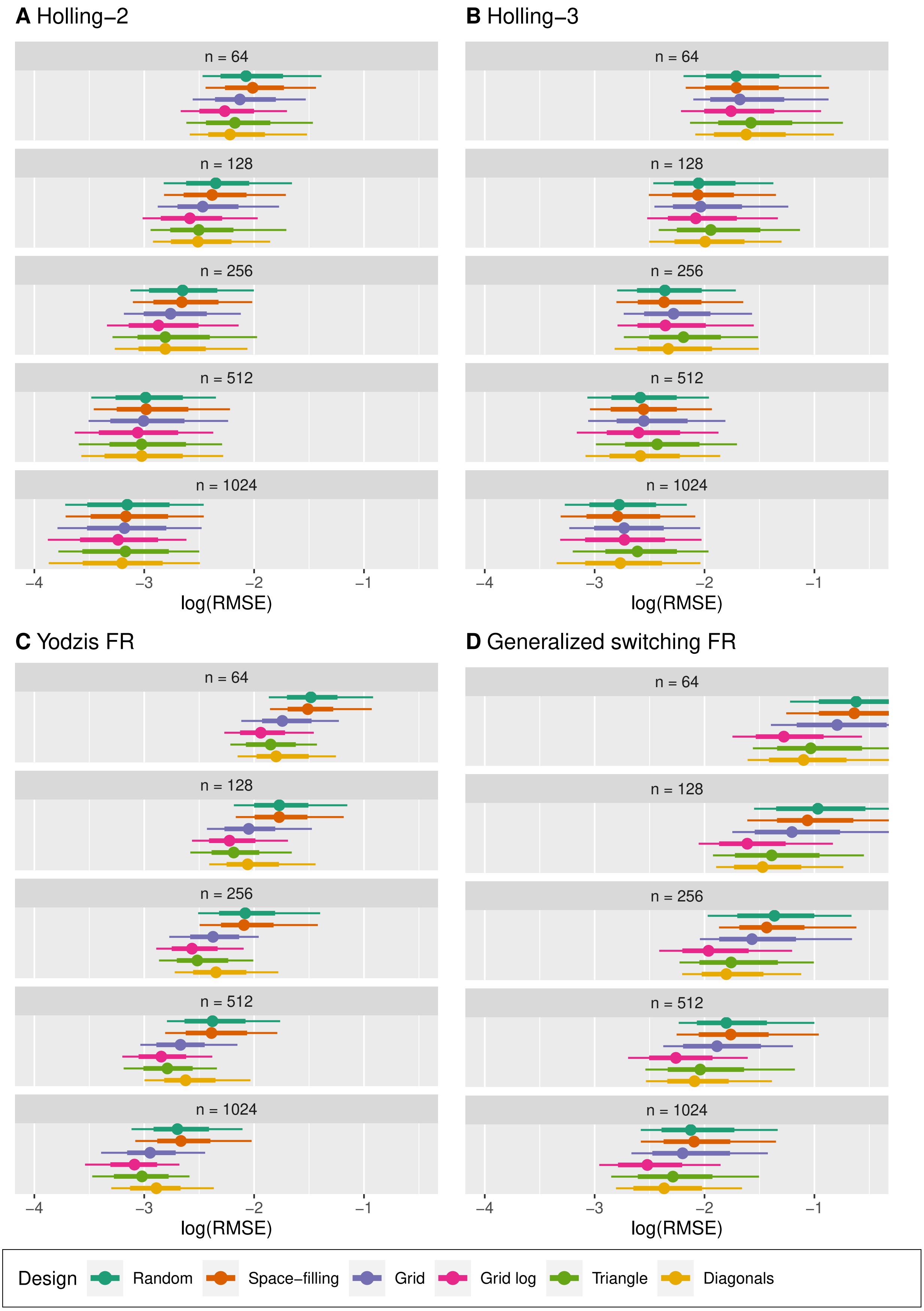


**Figure S6:** Distributions of relative RMSE vs. experimental design and sample size, **noise level: medium**. 500 datasets each. Dots represent means, bold lines are 66% intervals, thin lines are 95% intervals.

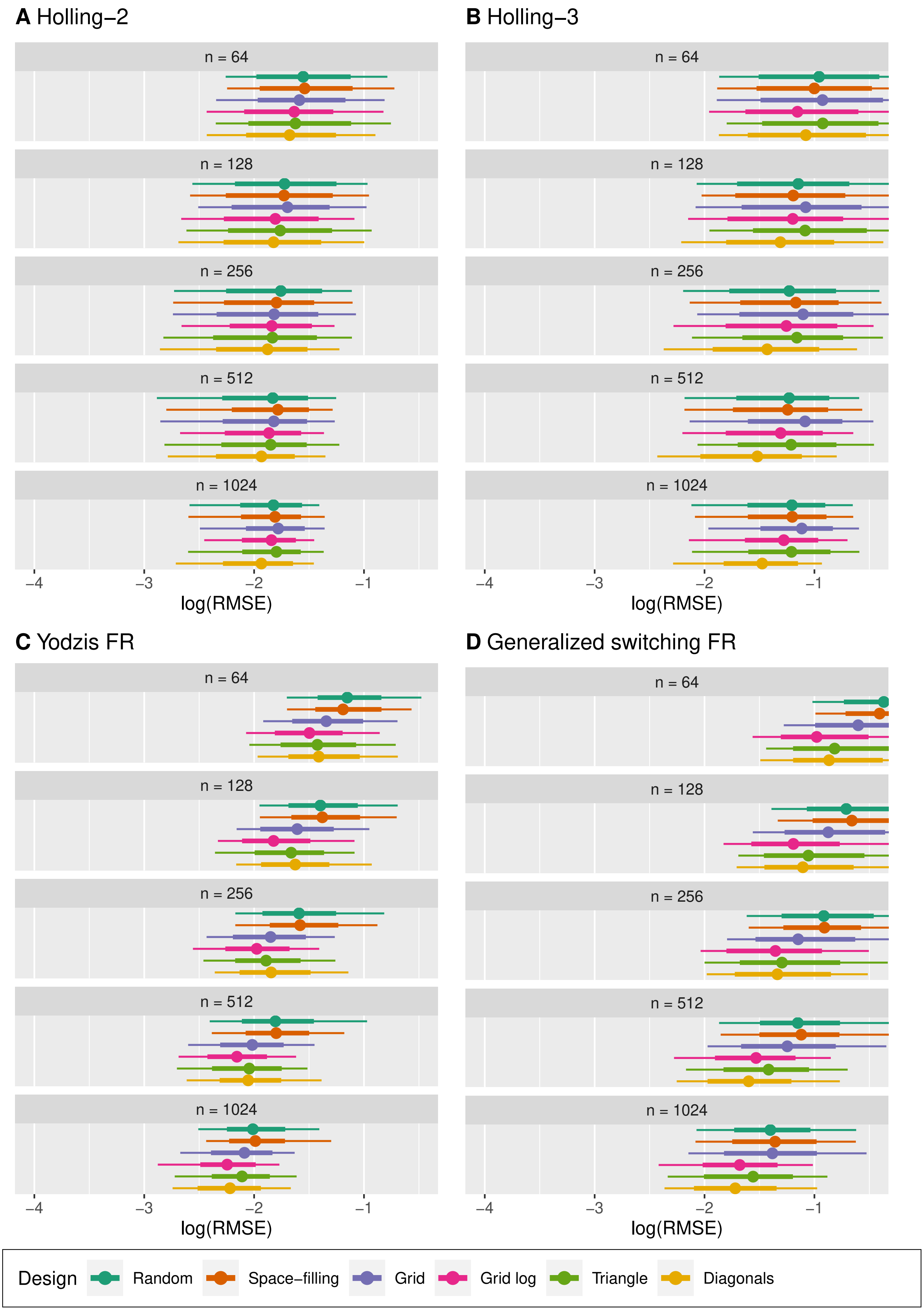


**Figure S7:** Distributions of relative RMSE vs. experimental design and sample size, **noise level: high**. 500 datasets each. Dots represent means, bold lines are 66% intervals, thin lines are 95% intervals.

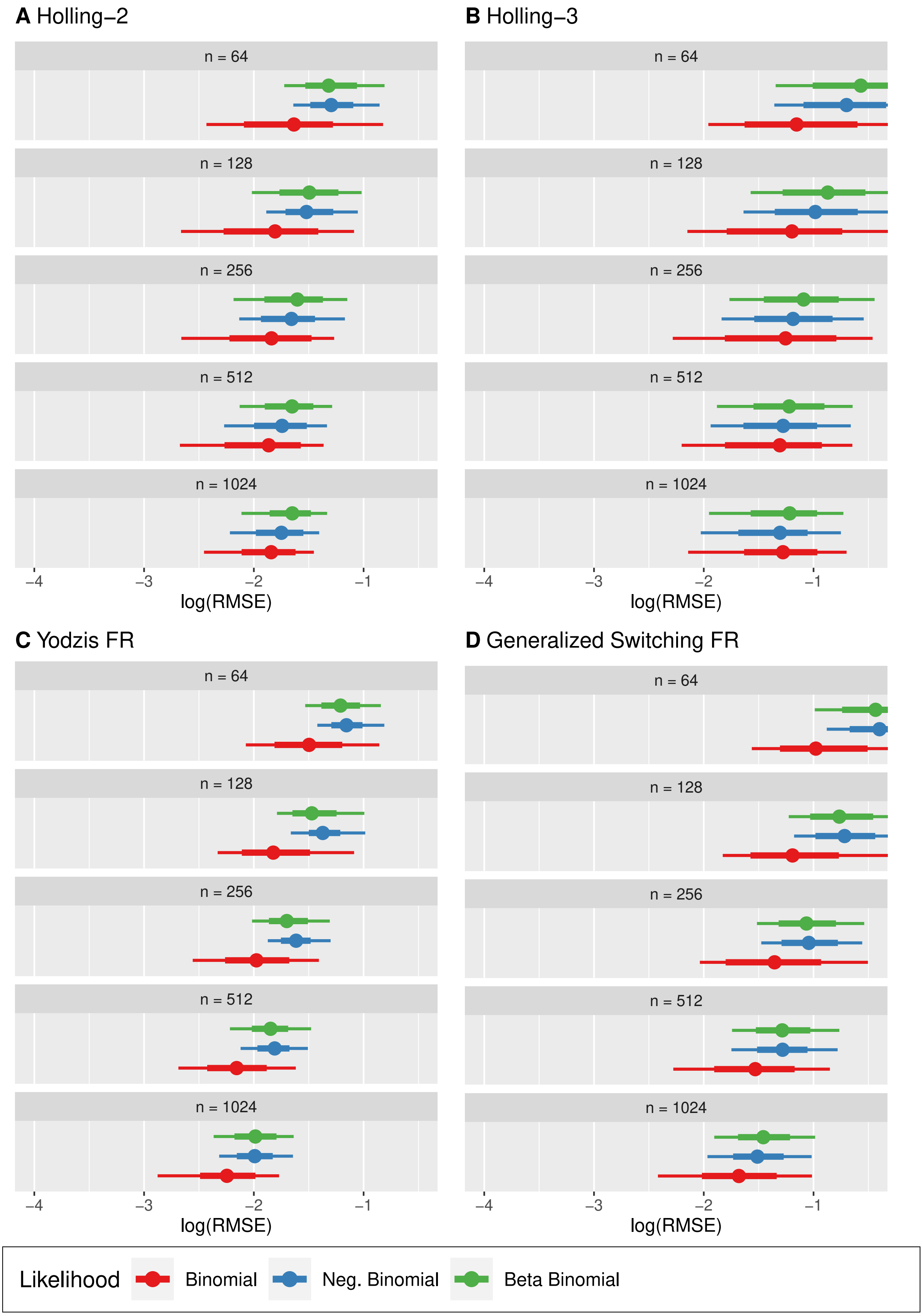


**Figure S8:** Distributions of relative RMSE vs. likelihood function and sample size, **noise level: high, design: log-spaced grid**. 500 datasets each. Dots represent means, bold lines are 66% intervals, thin lines are 95% intervals.

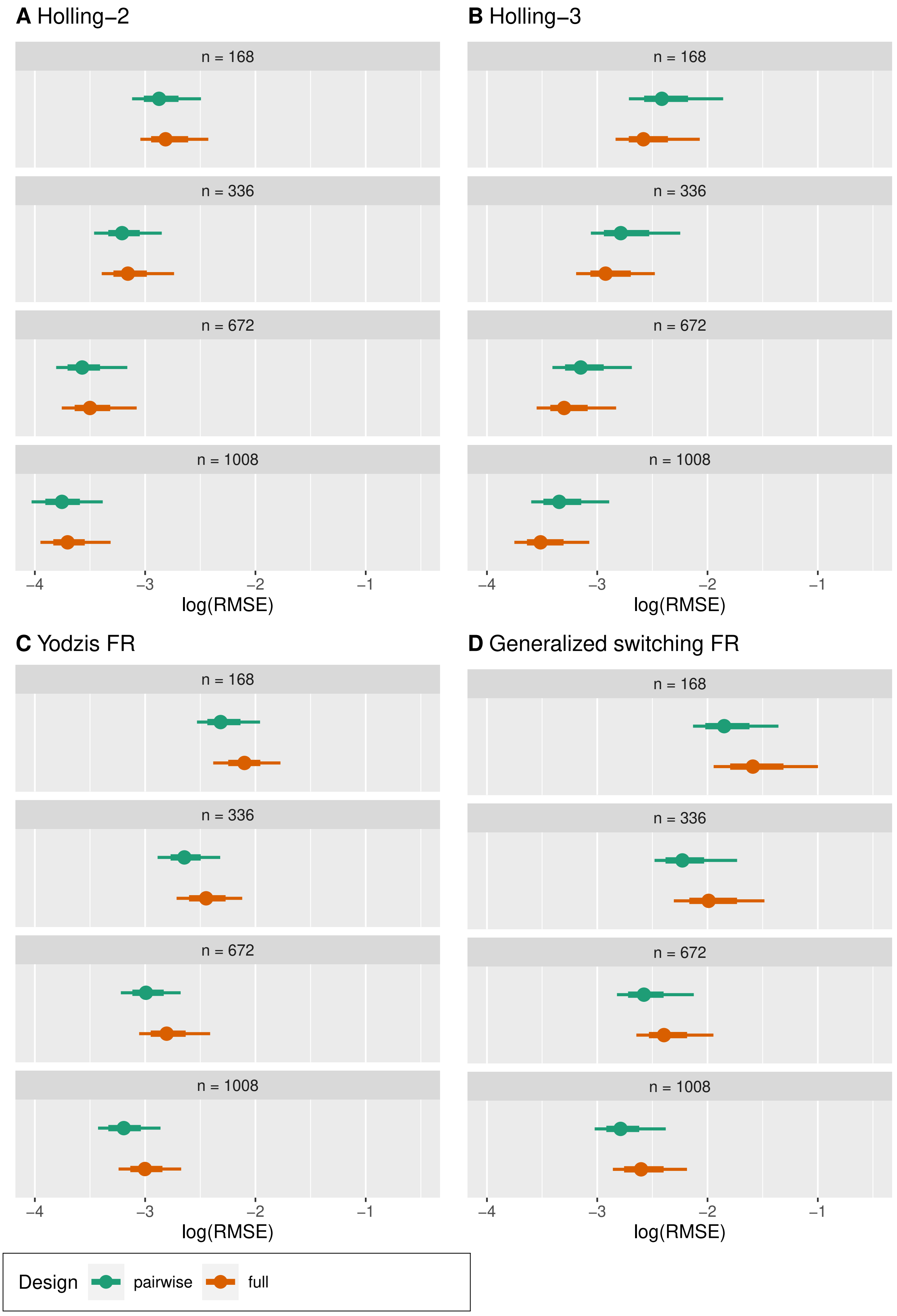


**Figure S9:** Three-prey scenarios. Distributions of relative RMSE vs. experimental design and sample size, **noise level: low**. 500 datasets each. Dots represent means, bold lines are 66% intervals, thin lines are 95% intervals.

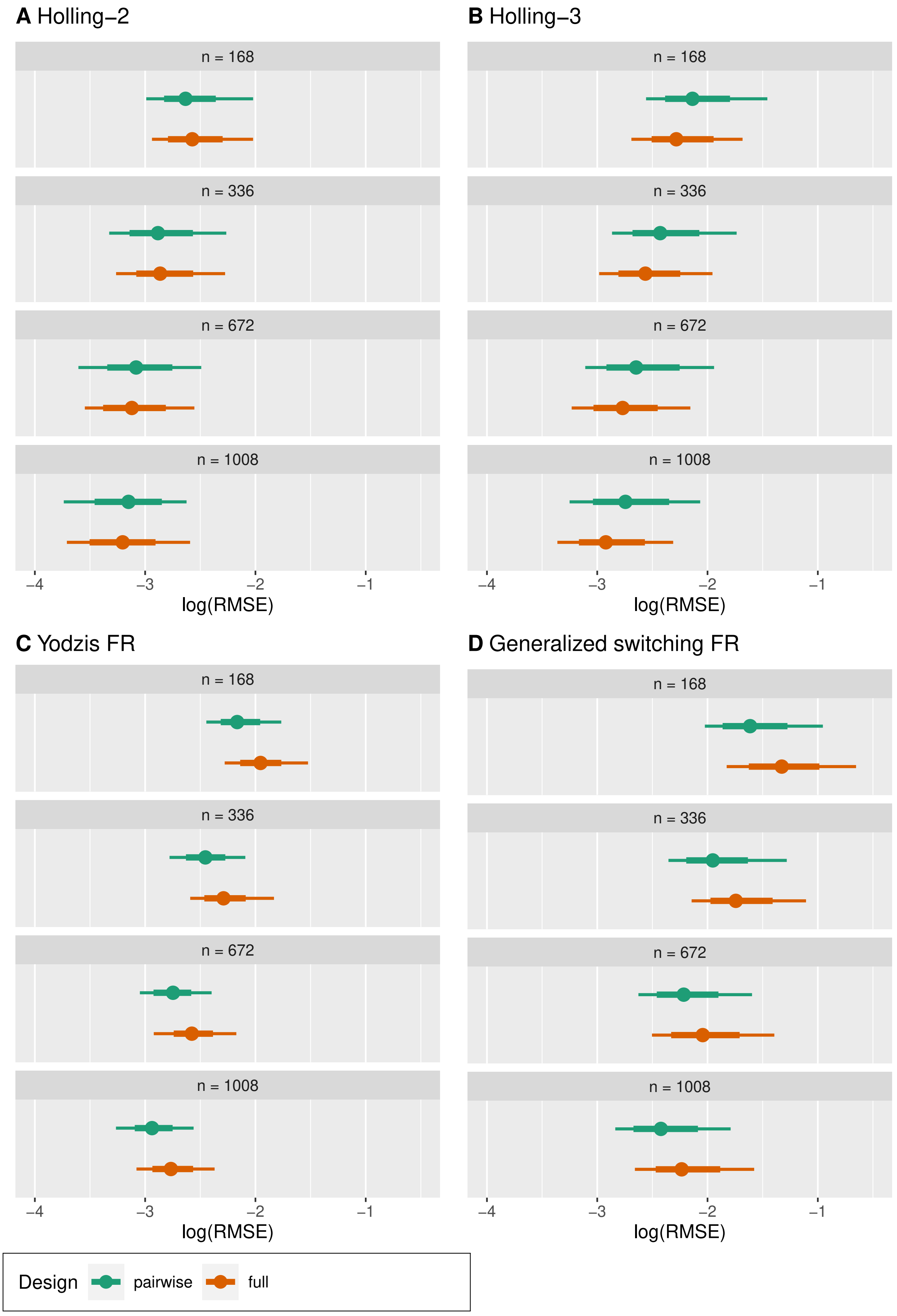


**Figure S10:** Three-prey scenarios. Distributions of relative RMSE vs. experimental design and sample size, **noise level: medium**. 500 datasets each. Dots represent means, bold lines are 66% intervals, thin lines are 95% intervals.

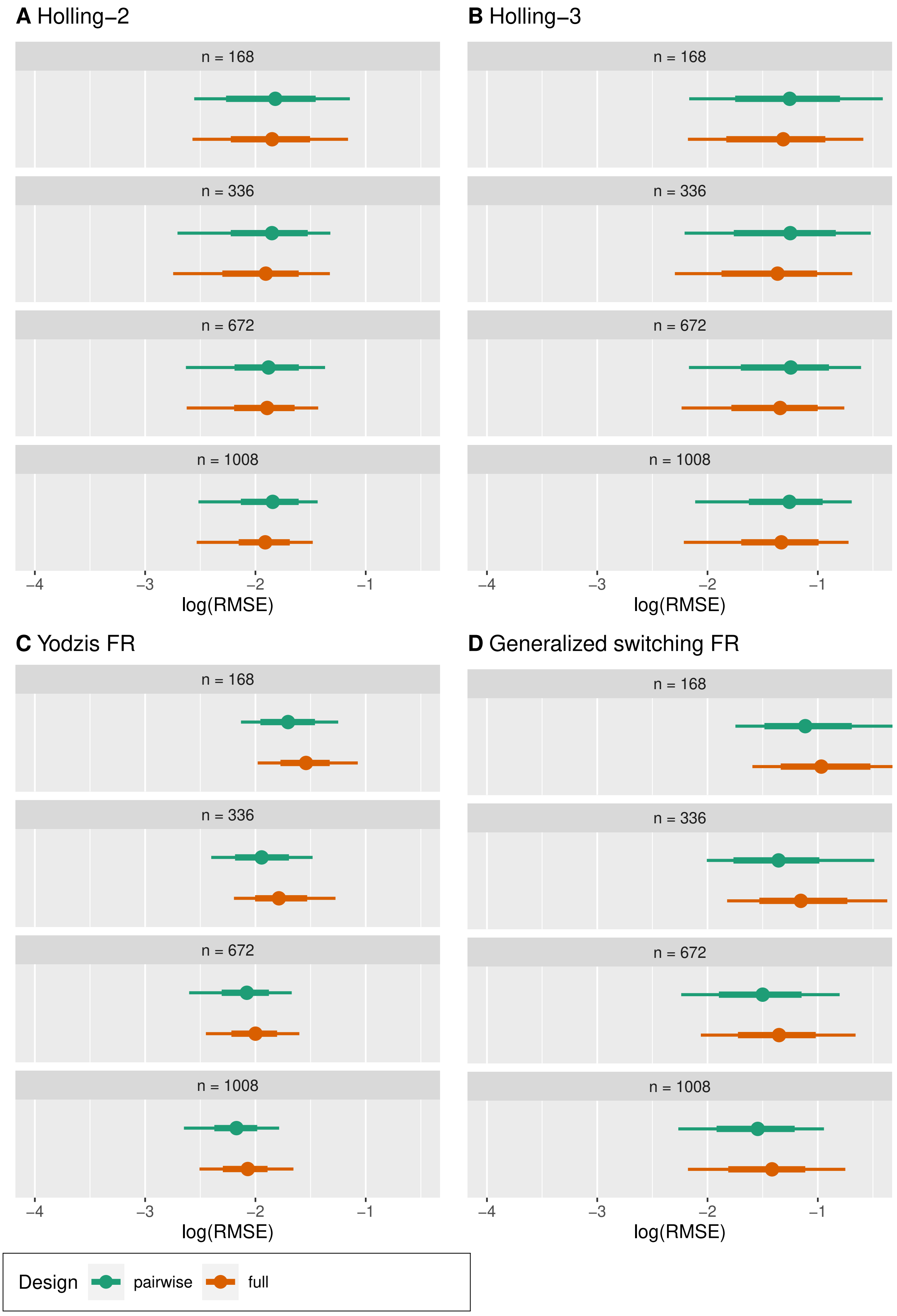


**Figure S11:** Three-prey scenarios. Distributions of relative RMSE vs. experimental design and sample size, **noise level: high**. 500 datasets each. Dots represent means, bold lines are 66% intervals, thin lines are 95% intervals.

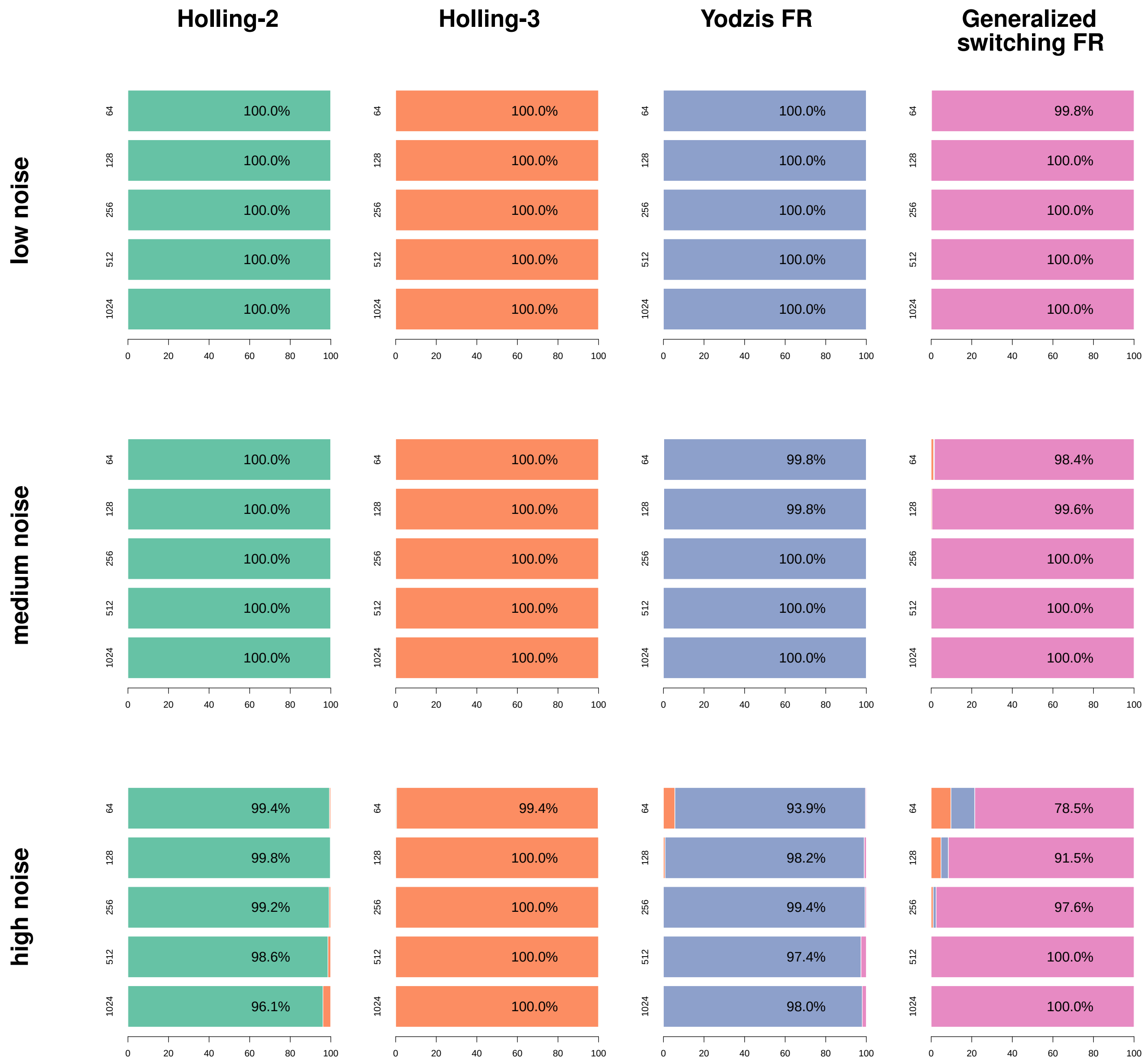


**Figure S12:** Model comparison: percentage of cases in which the correct model was identified. Columns indicate the model used for data simulation, outer rows indicate level of noise in the data, inner rows indicate sample size, colors identify best fitting model. 500 datasets were used for each combination of generating model, noise and sample size. All used the log-spaced grid as experimental design.

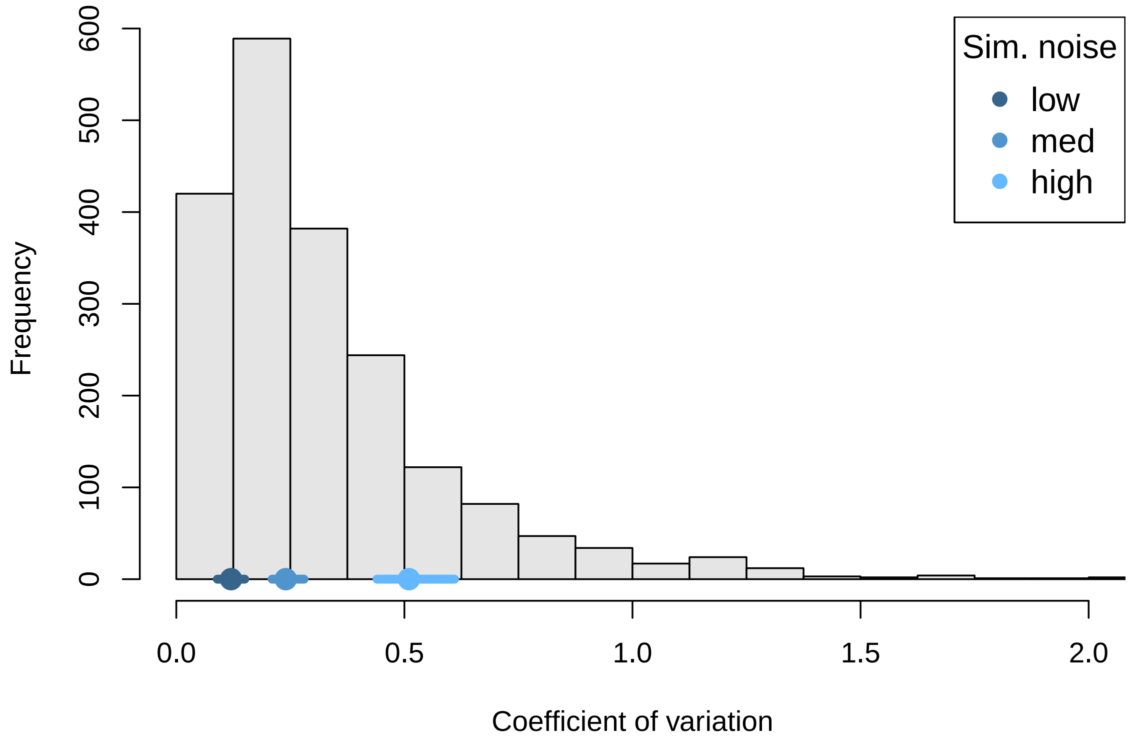


**Figure S13:** Comparison of noise in empirical datasets from the FoRAGE database (grey, histogram) and our synthetic datasets (blue, dots represent means and lines represent 95% intervals). We extracted 1,999 lab experiments (excluding field data) and computed coefficients of variation of eaten prey in the highest levels of offered prey. For raw data, we calculated empirical CVs. When only summary statistics were available (mean $\mu$ and standard deviation $SD$ or standard error $SE$), CVs were calculated as $\frac{\mu}{SD}$ or $\frac{\mu}{\sqrt{n}SE}$. Synthetic low, medium and high noise scenarios on average correspond to empirical 20%, 50% and 83% quantiles, respectively.

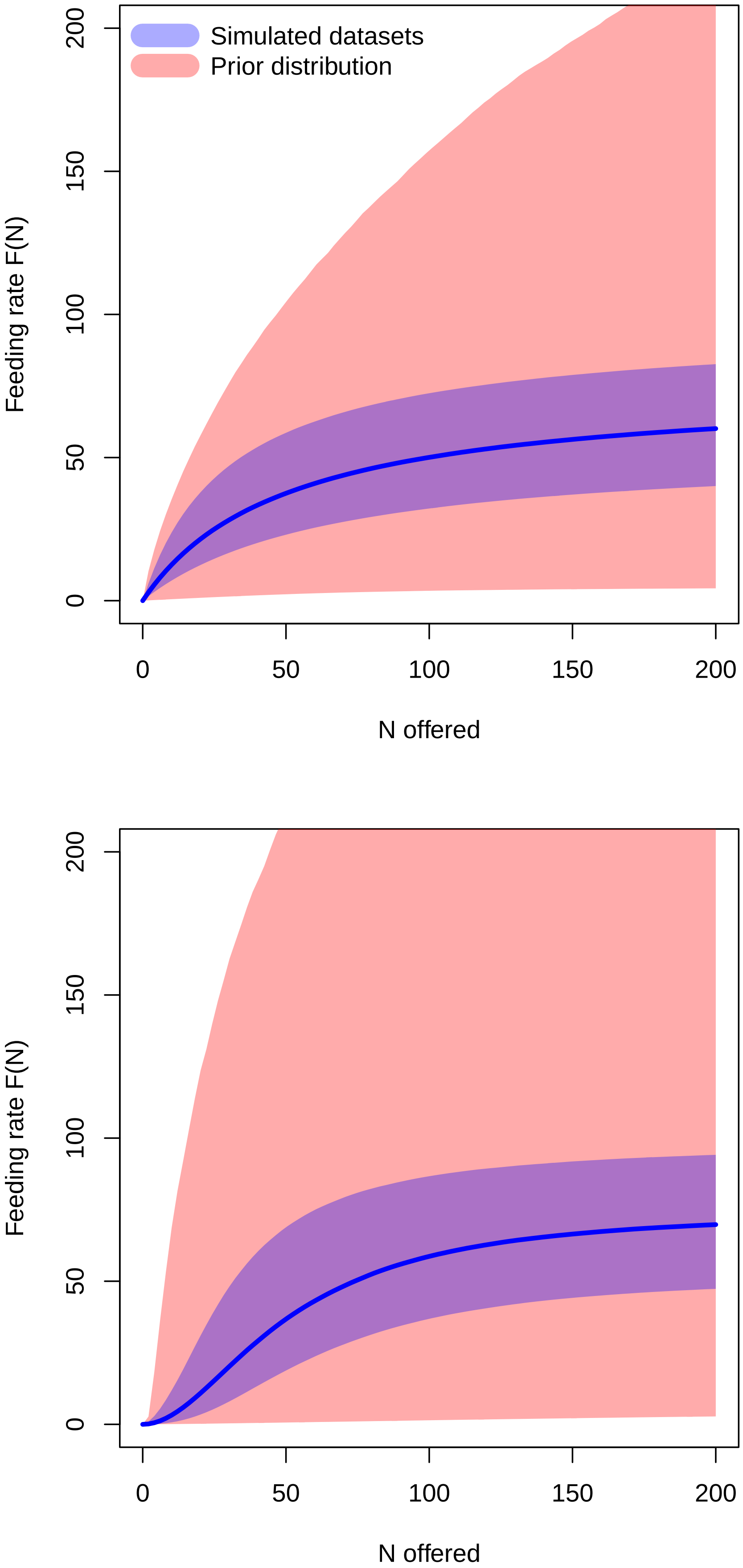


**Figure S14:** Prior predictive checks for Holling-2 (top) and Holling-3 (bottom). Because we could not perform individual checks for each of the thousands of synthetic datasets, we compared 90% intervals of single-species feeding rates used for data simulation (blue, $F_{max} \sim U(50, 100)$, $N_{half} \sim U(25, 75)$, $q \sim U(0.5, 1.5)$) with feeding rates sampled from prior distributions (blue, $a \sim Exp(0.5)$, $h \sim Exp(20.0)$, $b \sim Exp(10.0)$, $q \sim Exp(1.0)$). Prior distributions widely covering simulated feeding rates indicate vaguely informative priors.

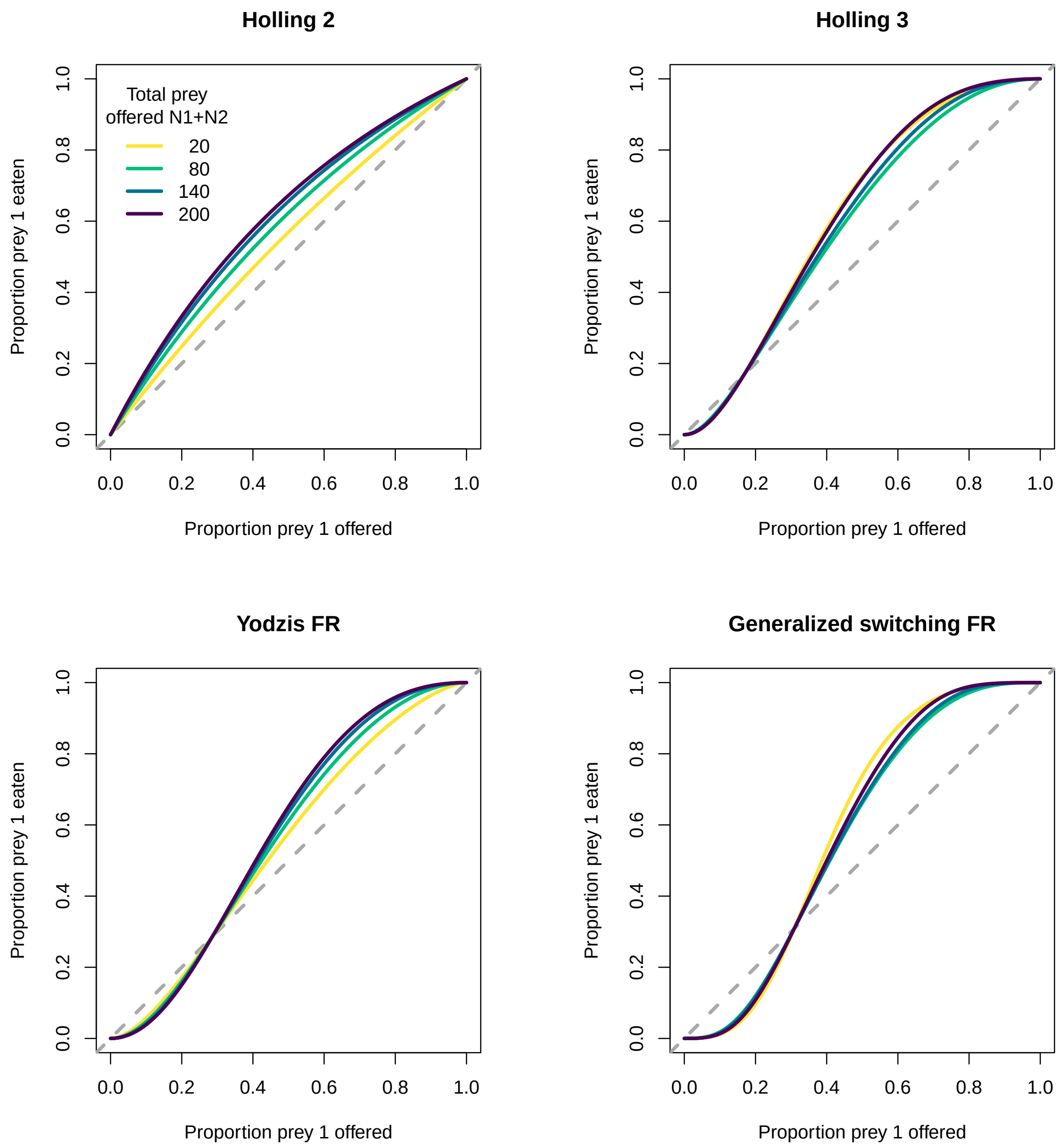


**Figure S15:** Proportion of eaten prey $\frac{N_{E,1}}{N_{E,1}+N_{E,2}}$ vs proportion of (initially) available prey $\frac{N_{0,1}}{N_{0,1}+N_{0,2}}$, colors indicating different numbers of total initial available prey $N_{0,1} + N_{0,2}$. Parameters: $F_{max} = (100, 80)$, $N_{half} = (25, 50)$, $q = 1$, $r = 1$, $w = (0.5, 0.5)$. In general, for all four MSFR models the proportion of feeding rates $\frac{F_1}{F_1+F_2}$ vs. proportion of prey availability $\frac{N_{0,1}}{N_{0,1}+N_{0,2}}$ is independent of total prey number $N_{0,1} + N_{0,2}$ (increasing both $N_{0,1}$ and $N_{0,2}$ by a factor $c$ increases $N_{0,1} + N_{0,2}$ by factor $c$, which cancels out in $\frac{N_{0,1}}{N_{0,1}+N_{0,2}}$ and in $\frac{F_1}{F_1+F_2}$). However, this only holds when prey abundance is constant throughout the whole feeding trial, whereas for ODE predictions (and experiments without prey replacement) $\frac{N_{E,1}}{N_{E,1}+N_{E,2}}$ depends on total prey abundance $N_{0,1} + N_{0,2}$. We conclude that two-prey experiments with a fixed sum of available prey abundances alone (triangle design) are not necessarily representative of all combinations of prey abundances.

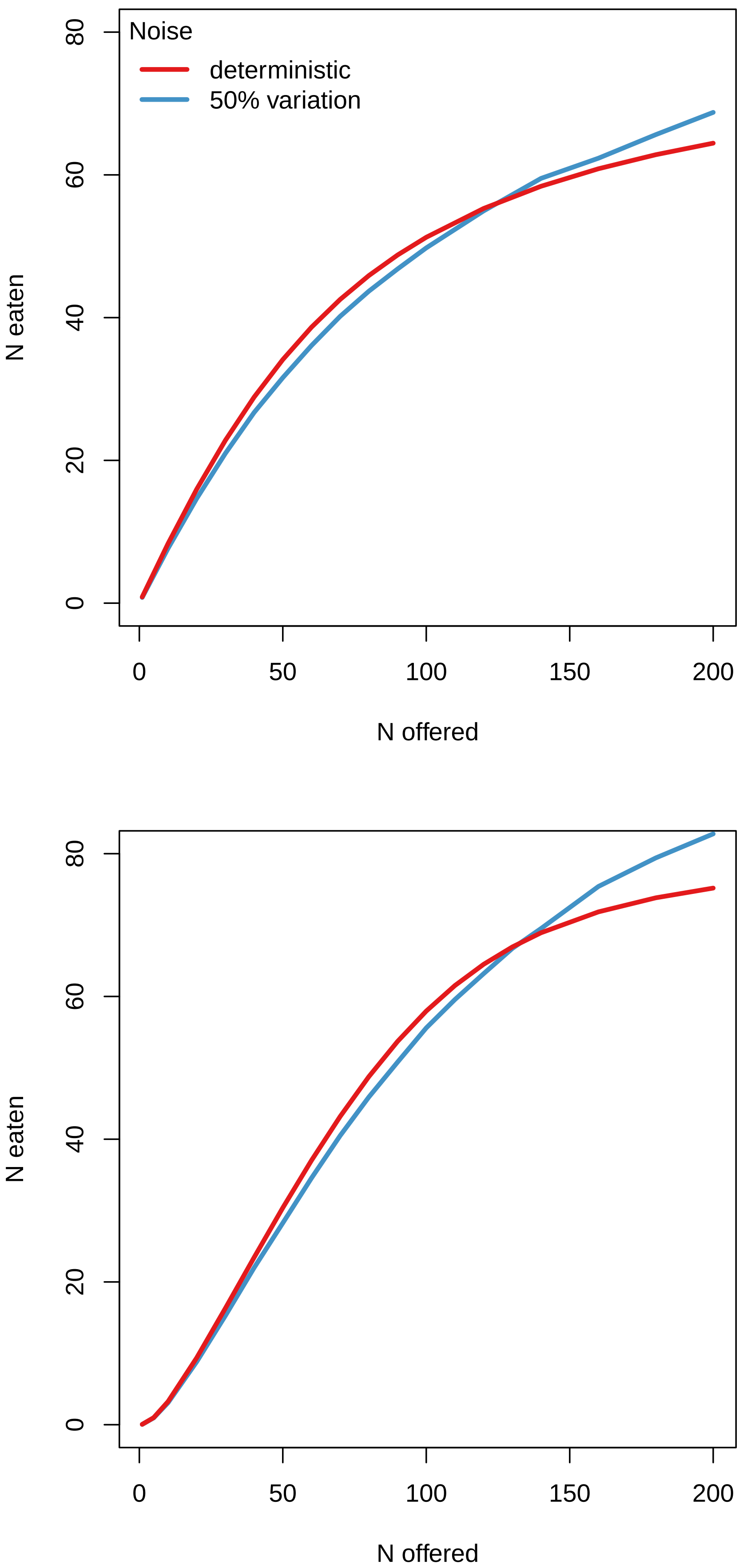


**Figure S16:** Illustration of Jensen's inequality for single-prey feeding trial data, top: Holling-2, bottom: Holling-3. Number of eaten prey for a predator with mean trait values (red) differs from mean of a distribution of data featuring variation in trait values (blue). Parameters: $F_{max} = 80$, $N_{half} = 40$, $q = 1$.

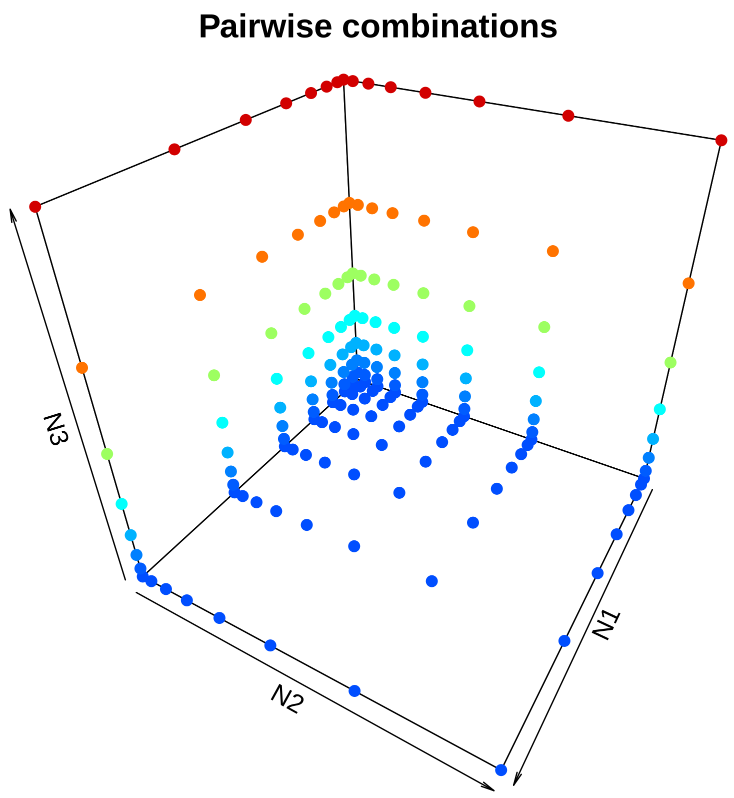


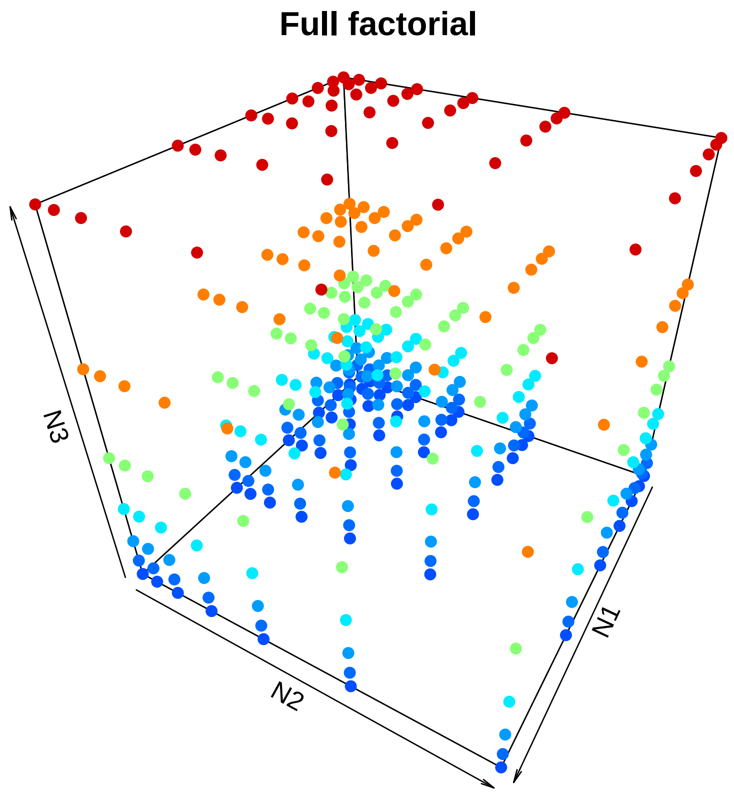


**Figure S17:** Experimental designs for three prey species. Top: Pairwise combinations of two prey species on three $8 \times 8$ grids with log-spaced distances, $n = 168$. Bottom: full factorial design of a $7 \times 7 \times 7$ grid with log-spaced distances, $n = 342$. Color indicates $N_3$ abundance for illustration.

**Table S1:** List of empirical datasets used. "Dataset" specifies the experiment when multiple were performed. "Obs" is the number of feeding trials in this experiment. "Replaced" denotes if eaten prey were constantly replaced. "Raw" specifies if data contains all individual observations, otherwise summary statistics of replicates were available. "Duration" indicates if all feeding trials were performed with identical or different experimental duration. "Best model" is the model with highest expected predictive accuracy according to the 'loo' package, "Alternative models" perform equally with respect to associated uncertainty. "Switch" indicates if the best model predicts prey switching. "RMSE" is the relative root mean squared error $\frac{1}{\bar{N}_E}\frac{1}{2n}\sqrt{\sum_{i=1}^{n}\sum_{j=1}^{2}\left(N_{Eji}-\widehat{N}_{Eji}\right)^2}$ of predicted eaten prey $\widehat{N}_{Eji}$, which is normalized by the observed mean of eaten prey $\bar{N}_E = \frac{1}{2n}\sum_{i=1}^{n}\sum_{j=1}^{2}N_{Eji}$. Individual results and code for each dataset are available on [Zenodo REF].

| Study | Data published | Dataset | Obs. | Replaced | Raw | Duration | Best model | Alternative models | Switch | RMSE |
|---|---|---|---|---|---|---|---|---|---|---|
| Colton (1987) | Novak and Stouffer (2020) | 1 | 108 | No | Yes | same | Holling 2 | Holling 3 | No | 0.463 |
| Colton (1987) | Novak and Stouffer (2020) | 2 | 108 | No | Yes | same | Holling 2 | Holling 3 | No | 0.394 |
| Cuthbert et al. (2018a) | Cuthbert et al. (2018b) | - | 72 | Yes & No | Yes | diff | Holling 2 | Holling 3 | No | 0.963 |
| Cuthbert et al. (2019) | Cuthbert et al. (2019) | ma | 65 | No | Yes | same | Holling 2 | Holling 3 | No | 0.346 |
| Cuthbert et al. (2019) | Cuthbert et al. (2019) | mf | 65 | No | Yes | same | Holling 2 | Holling 3, Yodzis, Gen. | No | 0.394 |
| Cuthbert et al. (2019) | Cuthbert et al. (2019) | mv | 65 | No | Yes | same | Holling 2 | Holling 3, Yodzis, Gen. | No | 0.370 |
| Cuthbert et al. (2020) | Cuthbert et al. (2023) | - | 115 | No | Yes | diff | Holling 2 | Holling 3, Yodzis, Gen. | No | 0.610 |
| Elliot (2006) | Elliot (2020) | i2 | 290 | Yes | Yes & No | same | Holling 2 | Holling 3 | No | 0.365 |
| Elliot (2006) | Elliot (2020) | i3 | 290 | Yes | Yes & No | same | Holling 2 | - | No | 0.224 |
| Elliot (2006) | Elliot (2020) | i4 | 290 | Yes | Yes & No | same | Holling 3 | - | Yes | 0.153 |
| Elliot (2006) | Elliot (2020) | i4B | 290 | Yes | Yes & No | same | Holling 3 | - | Yes | 0.165 |
| Elliot (2006) | Elliot (2020) | i5 | 290 | Yes | Yes & No | same | Holling 3 | - | Yes | 0.135 |
| Elliot (2006) | Elliot (2020) | i5B | 290 | Yes | Yes & No | same | Holling 3 | - | Yes | 0.135 |
| Joyce et al. (2019) | Joyce et al. (2019) | Ar | 89 | Yes & No | Yes | diff | Holling 2 | Holling 3 | No | 1.152 |
| Joyce et al. (2019) | Joyce et al. (2019) | Cm | 67 | Yes & No | Yes | diff | Holling 2 | Holling 3, Gen. | No | 1.078 |
| Long et al. (2012) | Long (2020) | - | 94 | No | Yes | same | Holling 2 | Holling 3, Yodzis, Gen. | No | 0.832 |
| McCard et al. (2021) | McCard et al. (2021) | - | 287 | Yes & No | Yes | diff | Holling 3 | Holling 2, Yodzis, Gen. | Yes | 0.464 |
| Ranta and Nuutinen (1985) | Novak and Stouffer (2020) | 10 | 123 | Yes | No | same | Holling 2 | Holling 3, Yodzis, Gen. | No | 0.362 |
| Ranta and Nuutinen (1985) | Novak and Stouffer (2020) | 13 | 123 | Yes | No | same | Yodzis | Gen. | No | 0.270 |
| Ranta and Nuutinen (1985) | Novak and Stouffer (2020) | 18 | 123 | Yes | No | same | Yodzis | Gen. | No | 0.297 |
| Ranta and Nuutinen (1985) | Novak and Stouffer (2020) | Ad | 123 | Yes | No | same | Yodzis | - | No | 0.307 |
| Wong and Barbeau (2005) | Wong and Barbeau (2020) | rc | 48 | Yes | Yes | same | Holling 2 | Yodzis | No | 0.463 |
| Wong and Barbeau (2005) | Wong and Barbeau (2020) | ss | 48 | Yes | Yes | same | Holling 2 | Holling 3, Yodzis, Gen. | No | 0.589 |

**Table S2:** Parameter estimates for empirical datasets. Posterior means and standard deviations for best fitting models. Parameter estimates for alternative models can be found in the online material [Zenodo REF].

| Dataset | Best model | $a_1$ | $a_2$ | $h_1$ | $h_2$ | $b_1$ | $b_2$ | $q$ | $w_1$ | $w_2$ | $r$ |
|---|---|---|---|---|---|---|---|---|---|---|---|
| Colton_1987_1 | H2 | 0.623 | 0.908 | 0.014 | 0.026 | | | | | | |
| | | (0.057) | (0.083) | (0.001) | (0.001) | | | | | | |
| Colton_1987_2 | H2 | 0.961 | 1.106 | 0.022 | 0.033 | | | | | | |
| | | (0.113) | (0.130) | (0.001) | (0.001) | | | | | | |
| Cuthbert_2018 | H2 | 1.670 | 0.652 | 0.001 | 0.001 | | | | | | |
| | | (0.097) | (0.056) | (0.001) | (0.001) | | | | | | |
| Cuthbert_2019_ma | H2 | 2.335 | 1.226 | 0.190 | 0.162 | | | | | | |
| | | (0.626) | (0.696) | (0.025) | (0.035) | | | | | | |
| Cuthbert_2019_mf | H2 | 4.033 | 2.546 | 0.116 | 0.132 | | | | | | |
| | | (0.843) | (0.572) | (0.012) | (0.017) | | | | | | |
| Cuthbert_2019_mv | H2 | 2.699 | 0.981 | 0.131 | 0.195 | | | | | | |
| | | (0.628) | (0.254) | (0.017) | (0.039) | | | | | | |
| Cuthbert_2020 | H2 | 0.346 | 3.182 | 0.129 | 0.063 | | | | | | |
| | | (0.040) | (0.353) | (0.019) | (0.003) | | | | | | |
| Elliot_2006_i2 | H2 | 0.018 | 0.012 | 0.059 | 0.089 | | | | | | |
| | | (0.002) | (0.002) | (0.042) | (0.061) | | | | | | |
| Elliot_2006_i3 | H2 | 0.036 | 0.027 | 0.049 | 0.059 | | | | | | |
| | | (0.005) | (0.003) | (0.023) | (0.029) | | | | | | |
| Elliot_2006_i4 | H3 | | | 0.060 | 0.060 | 0.020 | 0.026 | 0.301 | | | |
| | | | | (0.012) | (0.010) | (0.007) | (0.009) | (0.087) | | | |
| Elliot_2006_i4B | H3 | | | 0.087 | 0.078 | 0.011 | 0.008 | 0.629 | | | |
| | | | | (0.007) | (0.008) | (0.004) | (0.003) | (0.103) | | | |
| Elliot_2006_i5 | H3 | | | 0.046 | 0.050 | 0.022 | 0.031 | 0.474 | | | |
| | | | | (0.004) | (0.003) | (0.006) | (0.008) | (0.074) | | | |
| Elliot_2006_i5B | H3 | | | 0.055 | 0.052 | 0.028 | 0.018 | 0.598 | | | |
| | | | | (0.003) | (0.003) | (0.008) | (0.005) | (0.078) | | | |
| Joyce_2019_Ar | H2 | 0.884 | 0.606 | 0.001 | 0.003 | | | | | | |
| | | (0.039) | (0.030) | (0.001) | (0.002) | | | | | | |
| Joyce_2019_Cm | H2 | 8.393 | 4.972 | 0.003 | 0.004 | | | | | | |
| | | (0.717) | (0.386) | (0.001) | (0.001) | | | | | | |
| Long_2012 | H2 | 0.664 | 1.357 | 0.157 | 0.025 | | | | | | |
| | | (0.099) | (0.159) | (0.027) | (0.006) | | | | | | |
| McCard_2021 | H3 | | | 0.070 | 0.080 | 1.122 | 0.904 | 0.510 | | | |
| | | | | (0.003) | (0.003) | (0.118) | (0.095) | (0.096) | | | |
| Ranta_1985_10 | H2 | 3.103 | 4.142 | 0.063 | 0.237 | | | | | | |
| | | (0.531) | (0.835) | (0.004) | (0.015) | | | | | | |
| Ranta_1985_13 | Yodzis | 5.949 | 1.757 | 0.037 | 0.145 | | | | 0.126 | 0.874 | 0.020 |
| | | (0.913) | (0.407) | (0.002) | (0.010) | | | | (0.029) | (0.029) | (0.020) |
| Ranta_1985_18 | Yodzis | 5.870 | 1.829 | 0.033 | 0.106 | | | | 0.124 | 0.876 | 0.021 |
| | | (0.795) | (0.342) | (0.002) | (0.007) | | | | (0.025) | (0.025) | (0.020) |
| Ranta_1985_Ad | Yodzis | 1.975 | 2.262 | 0.050 | 0.039 | | | | 0.105 | 0.895 | 0.011 |
| | | (0.331) | (0.218) | (0.004) | (0.002) | | | | (0.022) | (0.022) | (0.011) |
| Wong_2005_rc | H2 | 5.939 | 14.602 | 0.025 | 0.026 | | | | | | |
| | | (0.878) | (1.867) | (0.001) | (0.001) | | | | | | |
| Wong_2005_ss | H2 | 7.570 | 0.468 | 0.019 | 0.002 | | | | | | |
| | | (1.565) | (0.039) | (0.001) | (0.002) | | | | | | |

Baudrot et al. (2016) derived a functional response $\Phi_i(x)$ mechanistically accounting for individually different handling times $h_i$:

$$\Phi_i(x) = \frac{\alpha(x)p_i(x)}{1 + \alpha(x)\sum_j h_j p_j(x)}. \tag{1}$$

We assume $\alpha(x)$ and $p_i(x)$ follow their formulation H3.2, where $\alpha(x)$ is associated with exponents $n_j$ and $p_i(x)$ are associated with exponents $m_j$

$$\alpha(x) = \sum_j a_j x^{n_j}$$
$$p_i(x) = \frac{a_i x^{m_i}}{\sum_j a_j x^{m_j}}$$

without loss of generality, i.e. they could take any of the proposed formulations. For identical handling times ($h_i = h$), exploiting $\sum_j p_j(x) = 1$ yields

$$\begin{aligned}\Phi_i(x) &= p_i(x)\frac{\alpha(x)}{1 + \alpha(x)h\sum_j p_j(x)} \\ &= p_i(x)\frac{\alpha(x)}{1 + \alpha(x)h} \\ &= p_i(x)\frac{\sum_j a_j x_j^{n_j}}{1 + h\sum_j a_j x_j^{n_j}}.\end{aligned} \tag{2}$$

In their supplement, the following formulation for individually different handling times $h_i$ was proposed (again, formulation H3.2 as an example)

$$\Phi_i(x) = p_i(x)\frac{\sum_j a_j x_j^{n_j}}{1 + \sum_j h_j a_j x_j^{n_j}}. \tag{3}$$

However, following eq. (1), the corresponding H3.2 formulation reads

$$\begin{aligned}\Phi_i(x) &= p_i(x)\frac{\sum_j a_j x^{n_j}}{1 + \left(\sum_j a_j x^{n_j}\right)\left(\sum_j h_j p_j(x)\right)} \\ &= p_i(x)\frac{\sum_j a_j x^{n_j}}{1 + \left(\sum_j a_j x^{n_j}\right)\left(\sum_j h_j \frac{a_j x_j^{m_j}}{\sum_k a_k x_k^{m_k}}\right)} \\ &= p_i(x)\frac{\sum_j a_j x^{n_j}}{1 + \left(\frac{\sum_j a_j x^{n_j}}{\sum_j a_j x^{m_j}}\right)\left(\sum_j h_j a_j x_j^{m_j}\right)}\end{aligned} \tag{4}$$

Here, (4) features an additional term $\left(\frac{\sum\cdots}{\sum\cdots}\right)$ and the sum involving $h_j$ features exponents $m_j$, while in (3) it features exponents $n_j$. Unless $\alpha(x)$ and $p_i(x)$ follow the same relationship (here: identical exponents $n_j = m_j$), (3) and (4) are different functional responses.
