## Supplementary material for "Towards understanding interactions in a complex world: Design and analysis of multi-species functional response experiments": Tutorial

### Tutorial: Fitting multi-species functional responses

Benjamin Rosenbaum

##### Introduction

This tutorial describes how to fit MSFR models to data in R using Bayesian inference with Stan. Ready-to-use Stan code for 4 different models (Holling-2, Holling-3, Yodzis FR, and the Generalized switching FR) is provided. A detailed description of the Stan code can be found at the end of this tutorial, but is not needed for the application.

The models can be fitted to data including single- and multi-species feeding trials simultaneously. Data with and without prey replacement can be fitted, and also a mixture of both is possible. For feeding trials without prey replacement, predictions are generated by numerical simulation of ODEs, and a Binomial likelihood is used. For feeding trials in which prey were constantly replaced, predictions are directly computed by the functional response formula, and a Poisson likelihood is used. Also, experimental durations can be provided for each feeding trial individually. However, we observed that strong differences in experimental duration lead to poor correspondance of observed and fitted values in some empirical testcases, potentially due to non-constant conditions over the whole duration of a feeding trial.

Fitted models are compared via an information criterion from the loo-package, that accounts for predictive accuracy as well as model complexity. Code for plotting observed against predicted feeding rates is provided as well.

#### Setup

```
rm(list=ls())
library("rstan")         # fitting Stan models
library("loo")           # model comparison
library("coda")          # plotting model output
library("BayesianTools") # plotting model output
library("deSolve")       # computing ODE predictions
```

Four Stan models are coded in `.stan` files. Compiling them takes a couple of minutes, but can be skipped if already executed before, since models are saved in `.Rdata` files.

```
model.H2 = stan_model("MSFR_H2.stan")
save(model.H2, file="stan_model_H2.RData")
model.H3 = stan_model("MSFR_H3.stan")
save(model.H3, file="stan_model_H3.RData")
model.Yo = stan_model("MSFR_Yo.stan")
save(model.Yo, file="stan_model_Yo.RData")
model.Gen = stan_model("MSFR_Gen.stan")
save(model.Gen, file="stan_model_Gen.RData")
```

Alternatively, previously compiled models are loaded.

```
load("stan_model_H2.RData")
load("stan_model_H3.RData")
load("stan_model_Yo.RData")
load("stan_model_Gen.RData")
```

#### Data simulation

**This whole section can be skipped if empirical data is used.** One of the four provided MSFR models can be used for simulating feeding experiments, for a given experimental design. Feeding trials are conducted with or without prey replacement. This also determines if a Binomial or a Poisson distribution is used to introduce variation around deterministic predicted feeding rates. Additionally, user-specified relative amount of between-individual predator trait variation increases variation in feeding rates.

First, functions for generating experimental design and functional response model are included.

```
source("experimental_designs.R")
source("MSFR_models.R")
```

Number of prey species, number of feeding trials, and maximum number of offered prey (per species) are chosen. Note that provided functions for generating an experimental design here work for 2 prey species only, but code for simulation and model fitting work for more than 2 species, too. Here, we set up a log-spaced grid design.

```
m = 2 # number of species
n = 128 # number of feeding trials: 64, 128, 256, 512, 1024
Nmax = 200 # maximum number of offered prey per species

DOE = doe.gridlog # choices:
df.all = DOE(n, Nmax) |> as.data.frame()

colnames(df.all) = c("Prey1", "Prey2", "Prey1.Eaten", "Prey2.Eaten" )
df.all$Replaced = FALSE
df.all$Time = 1.0
str(df.all)
```

```
'data.frame':   128 obs. of  6 variables:
 $ Prey1      : num  6 6 16 30 51 82 129 200 0 6 ...
 $ Prey2      : num  6 0 0 0 0 0 0 0 6 6 ...
 $ Prey1.Eaten: num  NA NA NA NA NA NA NA NA NA NA ...
 $ Prey2.Eaten: num  NA NA NA NA NA NA NA NA NA NA ...
 $ Replaced   : logi  FALSE FALSE FALSE FALSE FALSE FALSE ...
 $ Time       : num  1 1 1 1 1 1 1 1 1 1 ...
```

We choose a system following a Yodzis FR, with some amount of intraspecific predator trait variation. Population-level means for maximum feeding rate `Fmax` and half-saturation density `Nhalf` (prey density, for which feeding rate is half of maximum) are chosen for each prey species. They are converted into MSFR parameters attack rates / attack coefficients `a` and handling times `h`. This conversion is different for each MSFR model. Additionally, parameters `w`, `q` or `r` have to be specified, depending on the used model.

```
MSFR = Yodzis # Holling2, Holling3, Yodzis, New
strait = 0.1 # between [0.0, 0.4] in the manuscript
```

```r
set.seed(100)

# mean traits
Fmax = runif(m, 50, 100)
Nhalf = runif(m, 25, 75)
w = c(0.5,0.5)
# q = 1.0
r = 1.0

# pars.true = list(a=Fmax/Nhalf, h=1/Fmax) # Holling2
# pars.true = list(a=Fmax/Nhalf^(1+q), h=1/Fmax, q=q) # Holling3
pars.true = list(a=Fmax/Nhalf, h=1/Fmax, w=w, r=r) # Yodzis
# pars.true = list(a=Fmax/Nhalf^(1+q), h=1/Fmax, w=w, q=q, r=r) # Generalized

pars.true = append(pars.true, list(m=m, n=n, strait=strait, DOE=DOE))
pars.individual = pars.true
ai = rep(0,m)
hi = rep(0,m)
```

This for-loop simulates the `n` feeding trials from the experimental design. Number of offered prey is given in `df.all[i, 1:2]`. Each feeding trial is assigned individual attack rates and handling times randomly (predator traits with population-level means `a` and `h`). If prey is not replaced (`Replaced[i]==FALSE`), number of eaten prey is computed with an ODE simulation and a draw from a Binomial distribution. If prey are constantly replaced (`Replaced[i]==TRUE`), number of eaten prey is directly computed from the MSFR and a draw from a Poisson distribution. Each simulation accounts for individual experimental duration `Time[i]`. In this example, however, we chose identical duration (see above).

```r
for(i in 1:n){ # n feeding trials
  # individual predator traits
  for(j in 1:m){ # each species
    ai[j] = max(rnorm(1, pars.true$a[j], strait*pars.true$a[j]),
                pars.true$a[j]*(1-2*strait))
    hi[j] = max(rnorm(1, pars.true$h[j], strait*pars.true$h[j]),
                pars.true$h[j]*(1-2*strait))
  }
  pars.individual$a = ai
  pars.individual$h = hi
  if(df.all$Replaced[i]==0){
    # simulation with ODE
    out = ode(y = as.numeric(df.all[i, 1:m]),
```

```
                times = c(0,df.all$Time[i]),
                func = MSFR,
                parms = pars.individual)
      # integer eaten prey
      for(j in 1:m){
        df.all[i,m+j] = df.all[i,j]-out[2,j+1] # eaten prey
        if(df.all[i,m+j]>0){ # draw from binomial
          df.all[i,m+j] = rbinom(1, size=df.all[i,j], prob=df.all[i,m+j]/df.all[i,j])
        }
      }
    } else {
      # simulation, directly with MSFR function
      out = (-1)*unlist(MSFR(N=as.numeric(df.all[i, 1:m]),
                             parms = pars.individual))
      # integer eaten prey
      for(j in 1:m){
        df.all[i,m+j] = df.all$Time[i] * out[j] # eaten prey
        if(df.all[i,m+j]>0){ # draw from poisson
          df.all[i,m+j] = rpois(1, df.all[i,m+j])
        }
      }
    }
  }
  head(df.all)
```

```
  Prey1 Prey2 Prey1.Eaten Prey2.Eaten Replaced Time
1     6     6           2           4    FALSE    1
2     6     0           4           0    FALSE    1
3    16     0           8           0    FALSE    1
4    30     0          19           0    FALSE    1
5    51     0          29           0    FALSE    1
6    82     0          30           0    FALSE    1
```

Subsets of the complete dataset are used for plotting only.

```
df.single.1 = subset(df.all, Prey2==0)
df.single.2 = subset(df.all, Prey1==0)
df.multi = subset(df.all, Prey1>0 & Prey2>0)

par(mfrow=c(2,2))
plot(df.all[, 1:2], pch=16, xlab="Prey 1", ylab="Prey 2", main="Experimental design")
```

```
plot(df.single.1$Prey1, df.single.1$Prey1.Eaten,
     xlim=c(0,Nmax), ylim=c(0,max(df.all[, 3:4])),
     xlab="Prey 1 offered", ylab="Prey 1 eaten", main="Single-species feeding: Prey 1")
plot(df.single.2$Prey2, df.single.2$Prey2.Eaten,
     xlim=c(0,Nmax), ylim=c(0,max(df.all[, 3:4])),
     xlab="Prey 2 offered", ylab="Prey 2 eaten", main="Single-species feeding: Prey 2")
plot(df.multi$Prey1/(df.multi$Prey1+df.multi$Prey2),
     df.multi$Prey1.Eaten/(df.multi$Prey1.Eaten+df.multi$Prey2.Eaten),
     xlim=c(0,1), ylim=c(0,1),
     xlab="Proportion prey 1 offered", ylab="Proportion prey 1 eaten", main="Multi-species
abline(0,1)
```

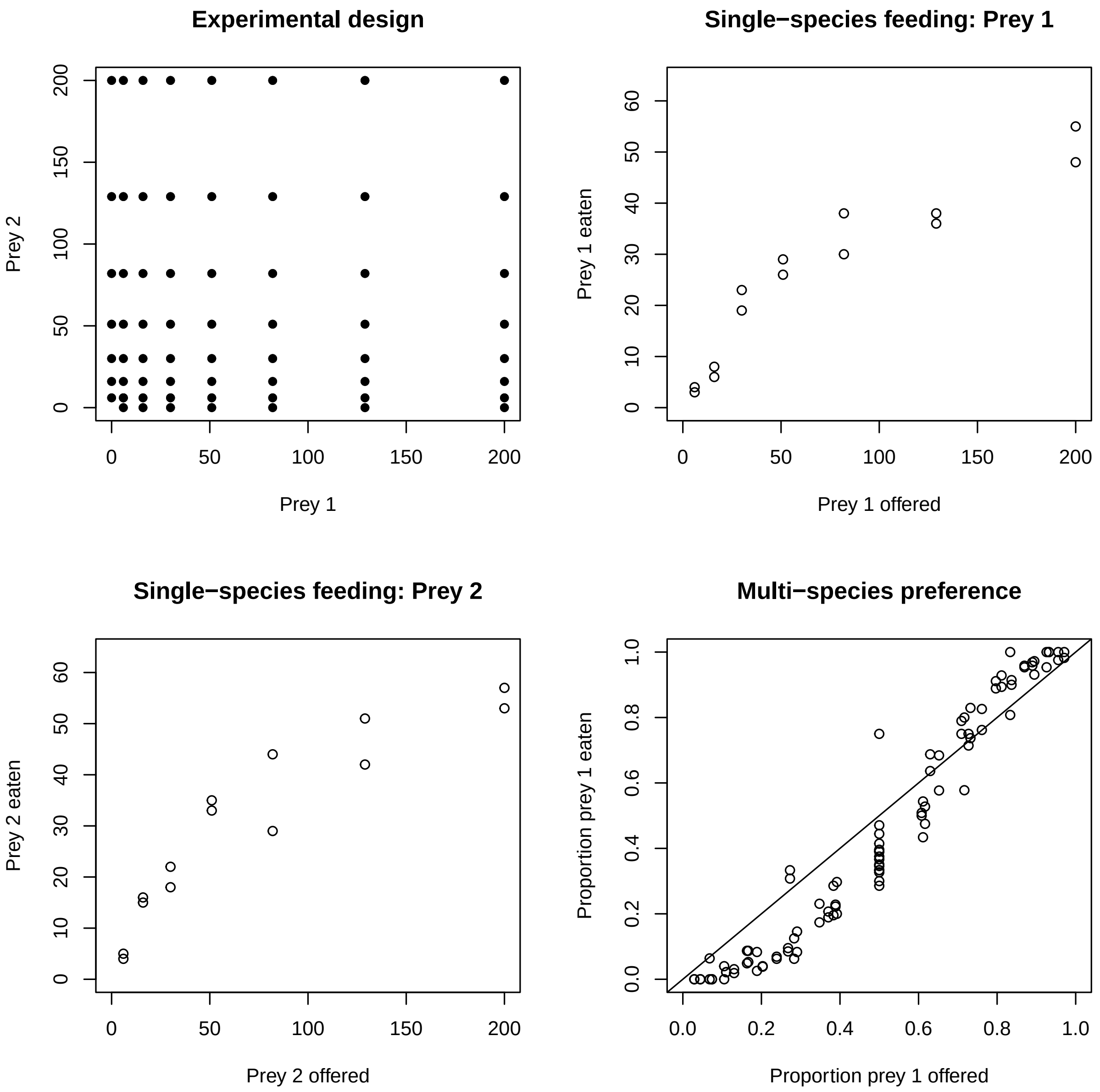

#### Model fitting

Data has to be coded as a named list for Stan, containing number of feeding trials `n`, number of prey species `m`, a matrix or data frame of offered prey `N_O` (colums identify prey species), an object of same size for number of eaten prey `N_E`, a vector indicating if prey were `Replaced`, and individual experimental duration `Time`.

```
data.stan = list(n=n,
                 m=m,
                 N_0=df.all[, 1:m],
                 N_e=df.all[, (m+1):(2*m)],
                 Replaced = as.integer(df.all$Replaced),
                 Time = df.all$Time
)
```

##### Fit Holling 2

Stan uses Markov-Chain-Monte-Carlo (MCMC) to sample from the posterior distribution of model parameters. Usually, 3–5 MCMC chains are used. These can be run in parallel on multicore processors, when options are specified accordingly (`cores = 3`). Further, we set the total number of iterations and number of warmup iterations for the sampling. The total number of posterior MCMC samples is `chains*(iter-warmup)`.

Starting values `init`for model parameters can provided as a list (in case of multiple chains: a list of lists), but Stan usually converges quickly towards the parameter region that feature a high probability density.

```
chains = 3

init = rep(list(list(a = c(1.0,1.0),
                     h = c(1/50,1/50))
),chains)

fit1 = sampling(
  model.H2,
  data = data.stan,
  chains = chains,
  warmup = 1000,
  iter = 3000,
  cores = 3,
  init = init,
  refresh = 100
)
```

```
print(fit1, pars=c("a","h"), digits=3, probs=c(0.05, 0.5, 0.95))
```

```
Inference for Stan model: anon_model.
```

```
3 chains, each with iter=3000; warmup=1000; thin=1;
post-warmup draws per chain=2000, total post-warmup draws=6000.

      mean se_mean    sd    5%   50%   95% n_eff  Rhat
a[1] 1.033   0.001 0.061 0.934 1.031 1.137  2180 1.001
a[2] 1.569   0.002 0.091 1.424 1.566 1.724  2171 1.001
h[1] 0.018   0.000 0.001 0.016 0.018 0.019  3028 1.000
h[2] 0.016   0.000 0.001 0.015 0.016 0.016  3367 1.000

Samples were drawn using NUTS(diag_e) at Wed Jan 24 09:49:02 2024.
For each parameter, n_eff is a crude measure of effective sample size,
and Rhat is the potential scale reduction factor on split chains (at
convergence, Rhat=1).
```

Convergence is verified by checking the effective sample size `n_eff` and the Gelman-Rubin diagnostics `Rhat` (should be <1.01), indicating that chains have mixed. Additionally, traceplots can be plotted (see below at the end of Model comparison section ).

##### Fit Holling 3

Analog to previous model fit.

```
init = rep(list(list(a = c(0.1,0.1),
                     h = c(1/50,1/50),
                     q = 1)
),chains)

fit2 = sampling(
  model.H3,
  data = data.stan,
  chains = chains,
  warmup = 1000,
  iter = 3000,
  cores = 3,
  init = init,
  refresh = 100
)
```

```
print(fit2, pars=c("a","h","q"), digits=3, probs=c(0.05, 0.5, 0.95))
```

```
Inference for Stan model: anon_model.
3 chains, each with iter=3000; warmup=1000; thin=1;
post-warmup draws per chain=2000, total post-warmup draws=6000.

      mean se_mean    sd    5%   50%   95% n_eff  Rhat
a[1] 0.165   0.001 0.027 0.124 0.163 0.212  1804 1.003
a[2] 0.293   0.001 0.045 0.226 0.291 0.373  1790 1.003
h[1] 0.022   0.000 0.001 0.021 0.022 0.023  3926 1.000
h[2] 0.019   0.000 0.001 0.019 0.019 0.020  3798 1.000
q    0.678   0.001 0.052 0.594 0.677 0.763  1828 1.002

Samples were drawn using NUTS(diag_e) at Wed Jan 24 09:57:32 2024.
For each parameter, n_eff is a crude measure of effective sample size,
and Rhat is the potential scale reduction factor on split chains (at
convergence, Rhat=1).
```

##### Fit Yodzis FR

Analog to previous model fit.

```r
init = rep(list(list(a = c(1,1),
                     h = c(1/50,1/50),
                     w = c(0.5,0.5),
                     r = 1)
),chains)

fit3 = sampling(
  model.Yo,
  data = data.stan,
  chains = chains,
  warmup = 1000,
  iter = 3000,
  cores = 3,
  init = init,
  refresh = 100
)
```

```r
print(fit3, pars=c("a","h","w","r"), digits=3, probs=c(0.05, 0.5, 0.95))
```

```
Inference for Stan model: anon_model.
```

```
3 chains, each with iter=3000; warmup=1000; thin=1;
post-warmup draws per chain=2000, total post-warmup draws=6000.

      mean se_mean    sd    5%   50%   95% n_eff  Rhat
a[1] 1.481   0.003 0.143 1.262 1.470 1.730  2562 1.000
a[2] 2.265   0.004 0.212 1.940 2.253 2.632  2355 1.001
h[1] 0.017   0.000 0.001 0.016 0.017 0.018  3087 1.000
h[2] 0.016   0.000 0.001 0.015 0.016 0.017  3363 1.001
w[1] 0.449   0.001 0.040 0.382 0.449 0.516  2211 1.001
w[2] 0.551   0.001 0.040 0.484 0.551 0.618  2211 1.001
r    0.857   0.001 0.065 0.754 0.855 0.967  5106 1.000

Samples were drawn using NUTS(diag_e) at Wed Jan 24 10:05:45 2024.
For each parameter, n_eff is a crude measure of effective sample size,
and Rhat is the potential scale reduction factor on split chains (at
convergence, Rhat=1).
```

##### Fit Generalized switching FR

Analog to previous model fit.

```r
init = rep(list(list(a = c(0.1,0.1),
                     h = c(1/50,1/50),
                     w = c(0.5,0.5),
                     q = 0.1,
                     r = 0.1)
),chains)

fit4 = sampling(
  model.Gen,
  data = data.stan,
  chains = chains,
  warmup = 1000,
  iter = 3000,
  cores = 3,
  init = init,
  refresh = 100
)
```

```r
print(fit4, pars=c("a","h","w","q","r"), digits=3, probs=c(0.05, 0.5, 0.95))
```

```
Inference for Stan model: anon_model.
3 chains, each with iter=3000; warmup=1000; thin=1;
post-warmup draws per chain=2000, total post-warmup draws=6000.

      mean se_mean    sd    5%   50%   95% n_eff  Rhat
a[1] 1.094   0.006 0.230 0.730 1.087 1.485  1292 1.000
a[2] 1.729   0.009 0.345 1.173 1.718 2.318  1355 1.000
h[1] 0.018   0.000 0.001 0.017 0.019 0.020  1765 1.000
h[2] 0.017   0.000 0.001 0.016 0.017 0.018  1846 1.000
w[1] 0.458   0.001 0.046 0.385 0.456 0.534  2652 1.001
w[2] 0.542   0.001 0.046 0.466 0.544 0.615  2652 1.001
q    0.130   0.002 0.075 0.020 0.122 0.265  1066 1.000
r    0.725   0.003 0.100 0.551 0.729 0.883  1439 1.000

Samples were drawn using NUTS(diag_e) at Wed Jan 24 10:23:58 2024.
For each parameter, n_eff is a crude measure of effective sample size,
and Rhat is the potential scale reduction factor on split chains (at
convergence, Rhat=1).
```

##### Model comparison

The loo package is used for model comparison with an information criterion, that accounts for predictive accuracy as well as model complexity. Log-likelihood values are extracted for all model fits. Then, the criterion (elpd-loo) is computed and compared among models. `loo_compare` lists the models in descending order (best model at the top), including difference to the best model and the associated uncertainty (standard deviation) of that difference.

```
lik1 = extract_log_lik(fit1, merge_chains=FALSE)
reff1 = relative_eff(exp(lik1), cores=4)
loo1 = loo(lik1, r_eff=reff1, cores=4)

lik2 = extract_log_lik(fit2, merge_chains=FALSE)
reff2 = relative_eff(exp(lik2), cores=4)
loo2 = loo(lik2, r_eff=reff2, cores=4)

lik3 = extract_log_lik(fit3, merge_chains=FALSE)
reff3 = relative_eff(exp(lik3), cores=4)
loo3 = loo(lik3, r_eff=reff3, cores=4)

lik4 = extract_log_lik(fit4, merge_chains=FALSE)
reff4 = relative_eff(exp(lik4), cores=4)
```

```r
loo4 = loo(lik4, r_eff=reff4, cores=4)

loo_compare(loo1, loo2, loo3, loo4)
```

```
       elpd_diff se_diff
model4    0.0       0.0  
model3   -0.3       1.7  
model2  -18.5       8.7  
model1 -149.9      21.3  
```

New FR has the best elpd score, but including uncertainty it performs equally to Yodzis FR (difference `0.3` with standard deviation `1.7`). Due to the principle of parsimony, we identify the less complex Yodzis FR as the best fitting model.

Its posterior distributions of all parameters and their multivariate correlation are plotted.

```r
plot(As.mcmc.list(fit3)[, 1:7])
```

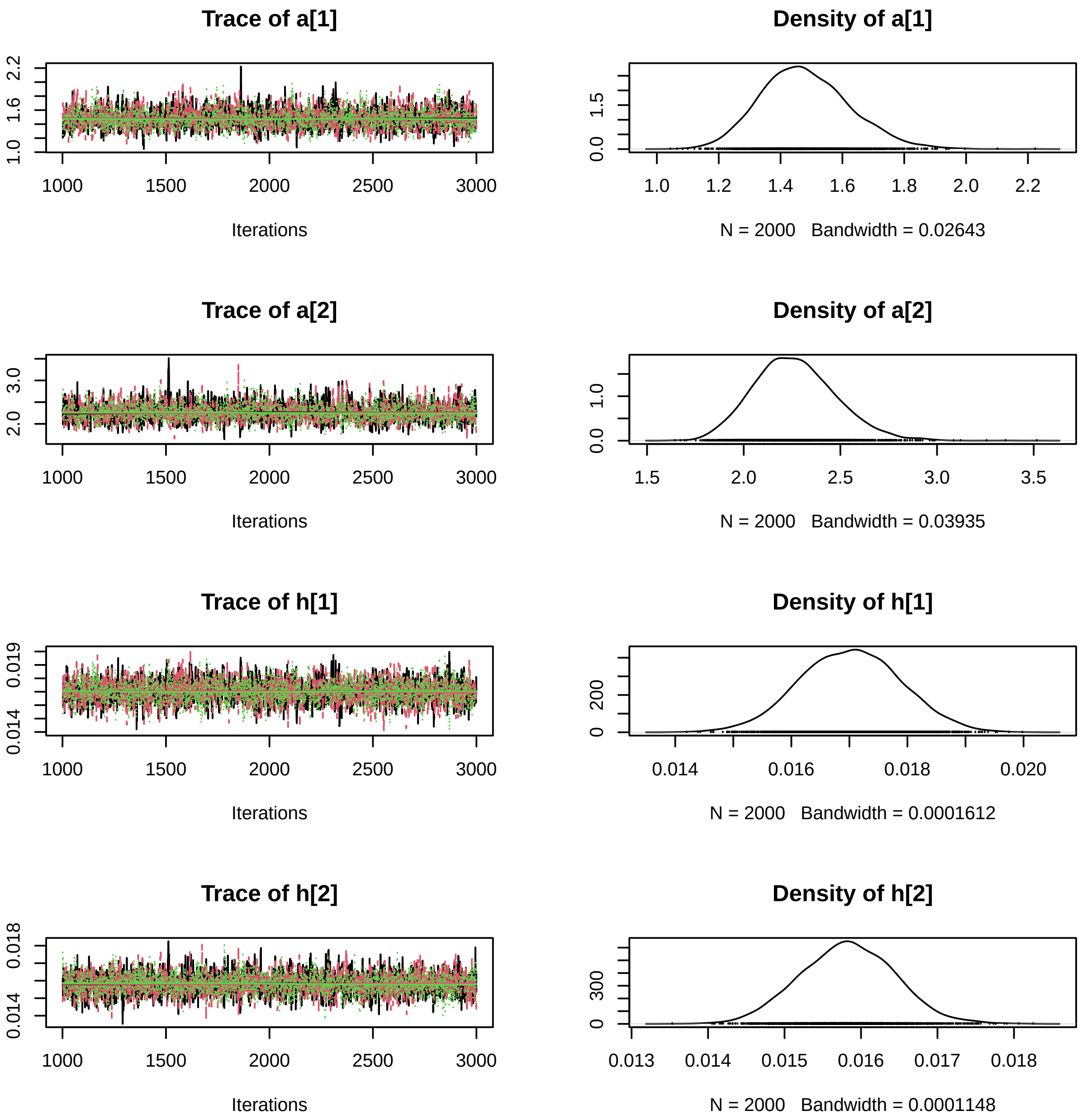
Trace of a[1]
Density of a[1]
Iterations
N = 2000 Bandwidth = 0.02643
Trace of a[2]
Density of a[2]
Iterations
N = 2000 Bandwidth = 0.03935
Trace of h[1]
Density of h[1]
Iterations
N = 2000 Bandwidth = 0.0001612
Trace of h[2]
Density of h[2]
Iterations
N = 2000 Bandwidth = 0.0001148

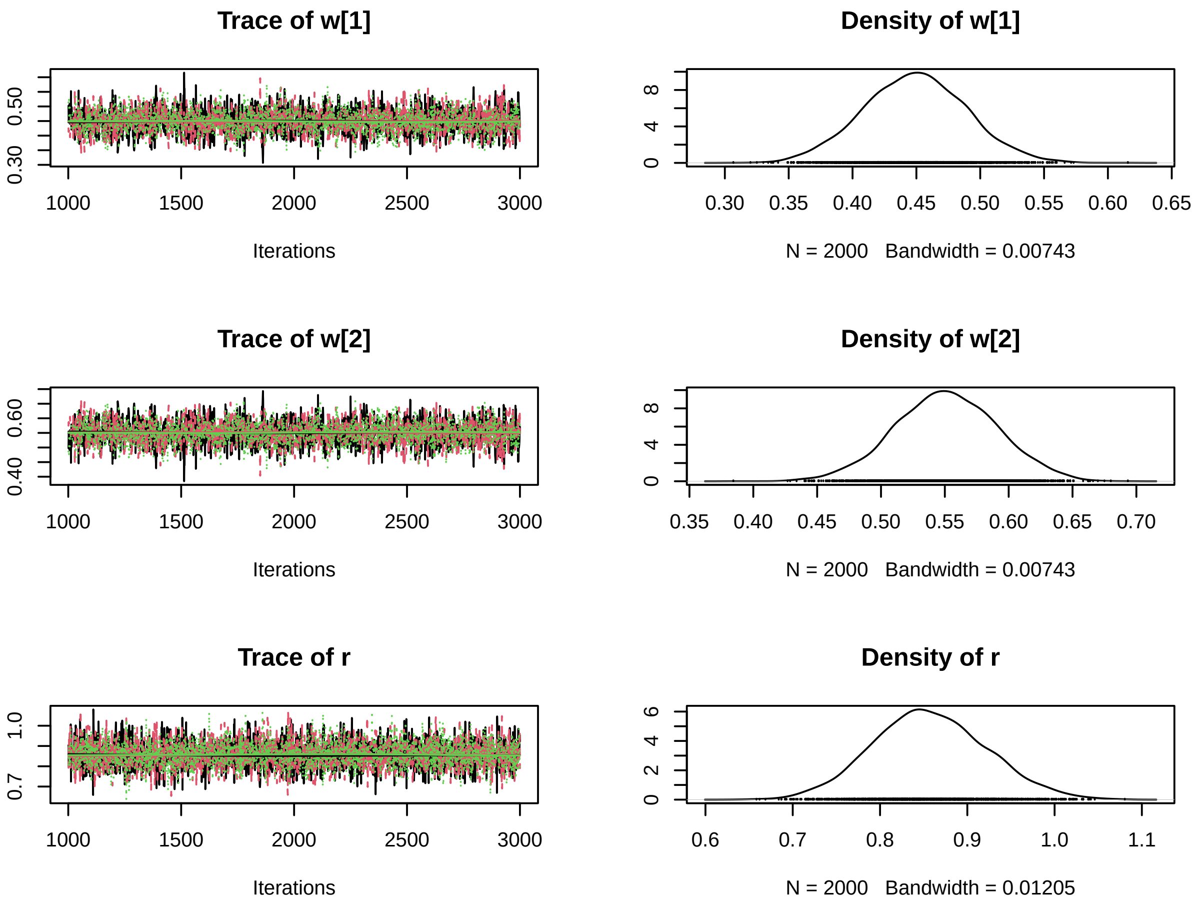

```r
correlationPlot(as.matrix(fit3)[, 1:7])
```

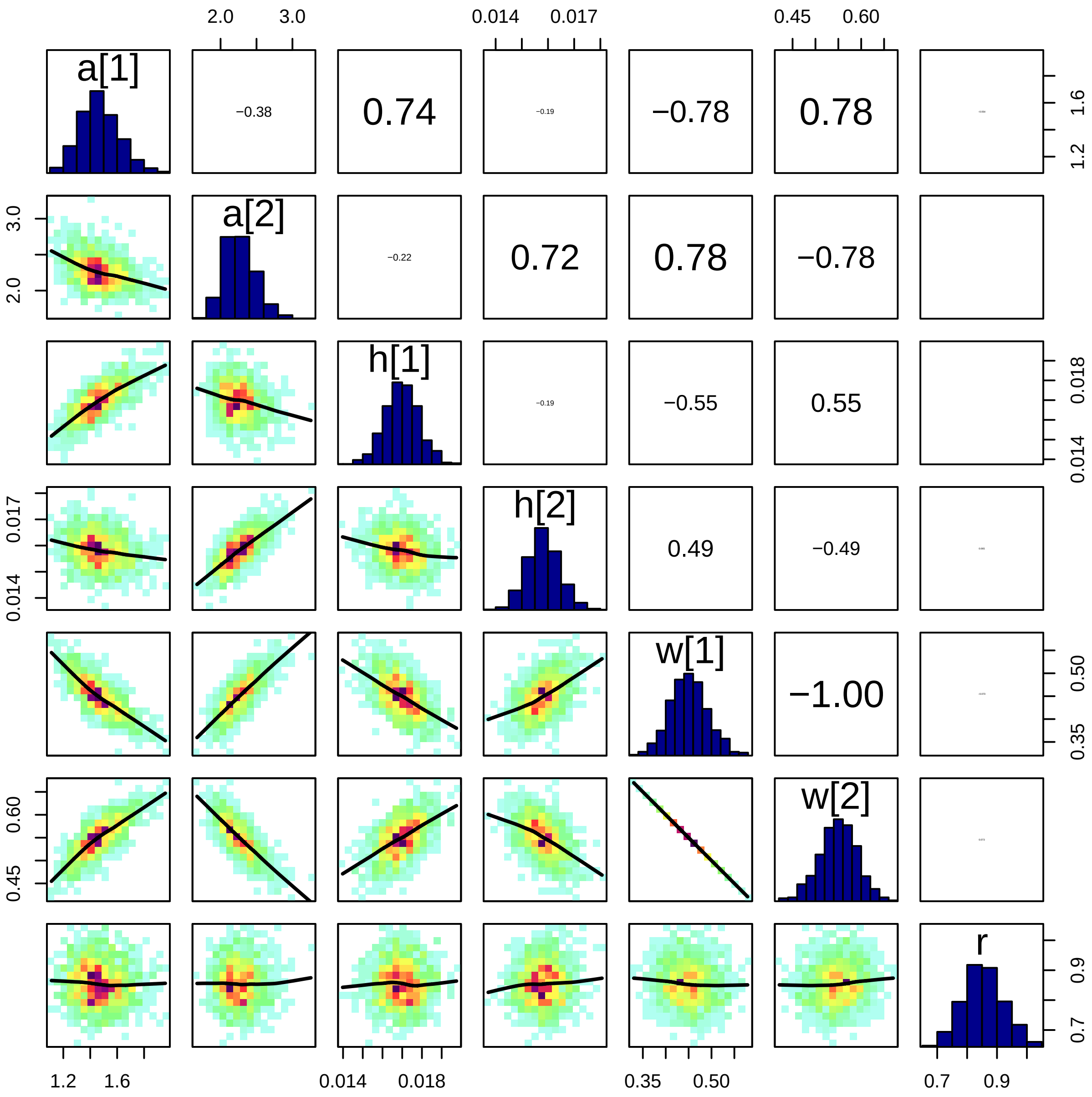

Since `w1=1-w2`, these parameters are perfectly correlated.

#### Posterior predictions for best fitting model

The correct `MSFR` model function is chosen for `deSolve`. 1000 samples from the posterior distribution are chosen randomly. For these, predictions are generated through numerical simulation of ODEs. Their median and 90% credible intervals are plotted against observed data.

```r
source("MSFR_models.R")
MSFR = Yodzis

n.post = 1000
n.pred = 20

post = as.matrix(fit3)[, 1:7]
set.seed(100)
post = post[sample(1:nrow(post),n.post), ]
head(post) |> round(digits=3)
```

```
          parameters
iterations   a[1]  a[2]  h[1]  h[2]  w[1]  w[2]     r
      [1,] 1.649 1.976 0.017 0.015 0.394 0.606 0.816
      [2,] 1.164 2.312 0.016 0.015 0.509 0.491 0.943
      [3,] 1.541 2.194 0.017 0.016 0.432 0.568 0.853
      [4,] 1.522 2.075 0.018 0.015 0.412 0.588 0.861
      [5,] 1.336 2.185 0.017 0.016 0.477 0.523 0.926
      [6,] 1.259 2.419 0.016 0.016 0.504 0.496 0.891
```

##### Single-species prey 1

Predictions for single-species experiments are computed for a gradient of offered prey densities `N10.pred`. Predictions are stored in the matrix `N1E.pred`, Here, Each row will contain a regression line for one sample of the posterior. Each column will contain a distribution of model predictions for a certain level of offered prey.

The outer loop is for each sample of the posterior, and model parameters are assigned from `post[k, ]`. The inner loop is for each number of offered prey `N10.pred[i]`. The second species' density is set to 0 to predict a single-species experiment.

```r
replaced = FALSE
time = 1.0

N10.pred = seq(0.1, Nmax, length.out=n.pred)
N1E.pred = matrix(NA, nrow=n.post, ncol=n.pred)

for(k in 1:n.post){
  params = list(a=post[k, 1:2],
                h=post[k, 3:4],
                w=post[k, 5:6],
```

```
                 r=post[k, 7],
                 m=2)
  for(i in 1:n.pred){
    if(replaced==FALSE){
      out = ode(y = c(N=c(N10.pred[i], 0.0)), times = c(0,time), func = MSFR, parms = para
      N1E.pred[k,i] = N10.pred[i]-out[2,2]
    } else{
      out = (-1)*unlist(MSFR(N=c(N10.pred[i], 0.0), parms=params))
      N1E.pred[k,i] = out[1]*time
    }
  }
}
```

Then, quantiles of the regression lines are extracted column-wise and plotted against observed data.

```
N1E.pred.qs = apply(N1E.pred, 2, function(x) quantile(x, probs=c(0.05, 0.5, 0.95)))

plot(df.single.1$Prey1, df.single.1$Prey1.Eaten,
     xlim=c(0,Nmax), ylim=c(0,max(df.all[, 3:4])),
     xlab="Prey 1 offered", ylab="Prey 1 eaten", main="Single-species feeding: Prey 1")
polygon( c(N10.pred, rev(N10.pred)),
         c(N1E.pred.qs[1, ], rev(N1E.pred.qs[3, ])),
         border=NA,
         col=adjustcolor("blue", alpha.f=0.2)
)
lines(N10.pred, N1E.pred.qs[2, ], col="blue", lwd=2)
```

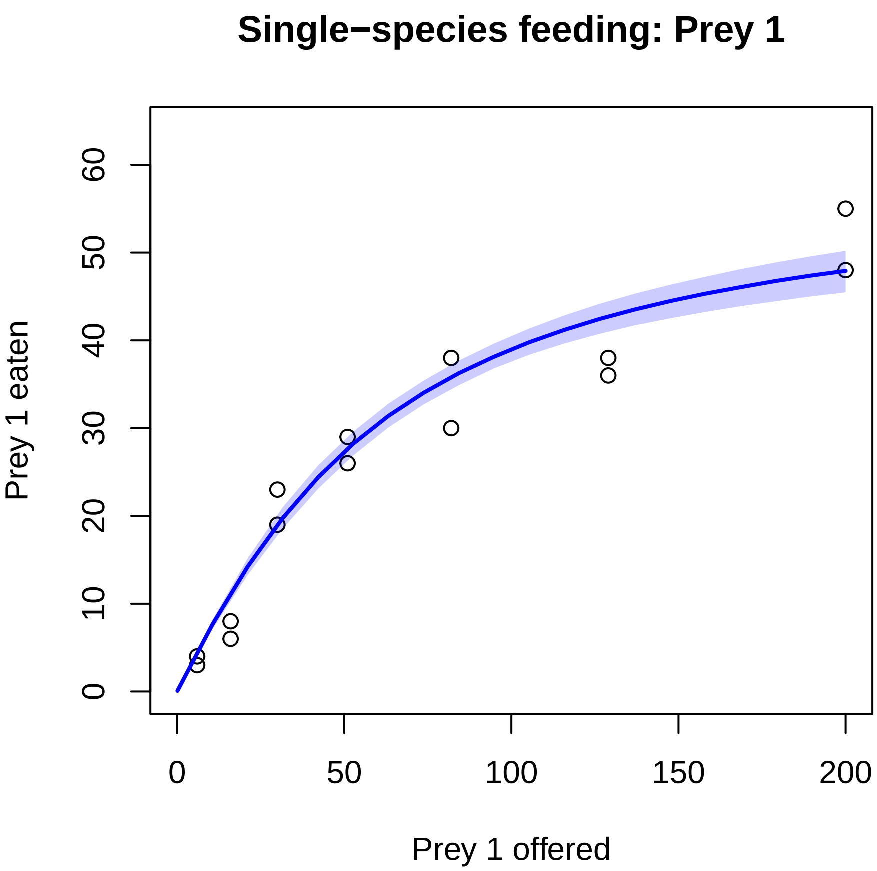

##### Single-species prey 2

Similarly, predictions for the second species' single-prey experiments are computed.

```
replaced = FALSE
time = 1.0

N2O.pred = seq(0.1, Nmax, length.out=n.pred)
N2E.pred = matrix(NA, nrow=n.post, ncol=n.pred)

for(k in 1:n.post){
  params = list(a=post[k, 1:2],
                h=post[k, 3:4],
                w=post[k, 5:6],
                r=post[k, 7],
                m=2)
```

```
  for(i in 1:n.pred){
    if(replaced==FALSE){
      out = ode(y = c(N=c(0.0, N20.pred[i])), times = c(0,time), func = MSFR, parms = para
      N2E.pred[k,i] = N20.pred[i]-out[2,3]
    } else{
      out = (-1)*unlist(MSFR(N=c(0.0, N20.pred[i]), parms=params))
      N2E.pred[k,i] = out[2]*time
    }
  }
}

N2E.pred.qs = apply(N2E.pred, 2, function(x) quantile(x, probs=c(0.05, 0.5, 0.95)))

plot(df.single.2$Prey2, df.single.2$Prey2.Eaten,
     xlim=c(0,Nmax), ylim=c(0,max(df.all[, 3:4])),
     xlab="Prey 2 offered", ylab="Prey 2 eaten", main="Single-species feeding: Prey 2")
polygon( c(N20.pred, rev(N20.pred)),
         c(N2E.pred.qs[1, ], rev(N2E.pred.qs[3, ])),
         border=NA,
         col=adjustcolor("red", alpha.f=0.2)
)
lines(N20.pred, N2E.pred.qs[2, ], col="red", lwd=2)
```

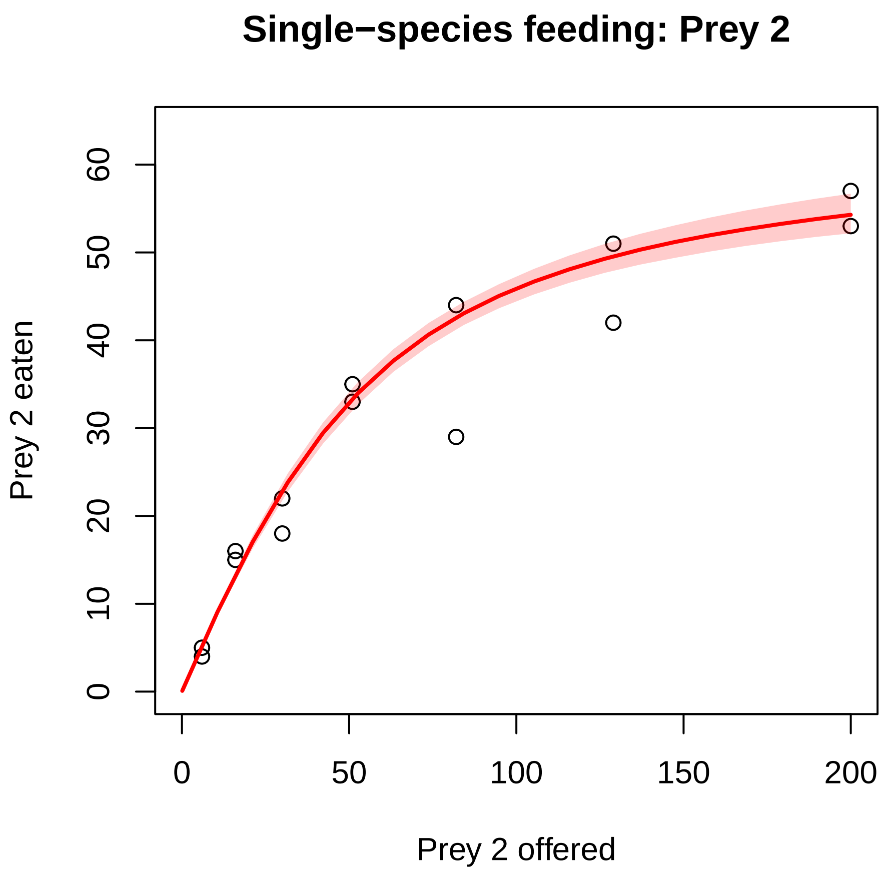

###### Multi-species relative consumption

Multi-species predictions are computed along a gradient of relative composition of $N_{0,1}$ and $N_{0,2}$ with a constant level of total prey offered $N_{0,1} + N_{0,2} = 200$.

```
replaced = FALSE
time = 1.0

N0.total = 200
N10.pred = seq(0, N0.total, length.out=n.pred)
N20.pred = N0.total-N10.pred
N1E.pred = matrix(NA, nrow=n.post, ncol=n.pred)
N2E.pred = matrix(NA, nrow=n.post, ncol=n.pred)

for(k in 1:n.post){
  params = list(a=post[k, 1:2],
```

```
                    h=post[k, 3:4],
                    w=post[k, 5:6],
                    r=post[k, 7],
                    m=2)
    for(i in 1:n.pred){
      if(replaced==FALSE){
        out = ode(y = c(N=c(N10.pred[i], N20.pred[i])), times = c(0,time), func = MSFR, parm
        N1E.pred[k,i] = N10.pred[i]-out[2,2]
        N2E.pred[k,i] = N20.pred[i]-out[2,3]
      }
      else{
        out = (-1)*unlist(MSFR(N=c(N10.pred[i], N20.pred[i]), parms=params))
        N1E.pred[k,i] = out[1]
        N2E.pred[k,i] = out[2]
      }
    }
  }
```

The predictions $N_{E,1}$, $N_{E,2}$ are transformed to proportion of prey 1 eaten feeding $\frac{N_{E1}}{N_{E1}+N_{E2}}$ (relative feeding). Their quantiles are extracted and plotted against proportions of offered prey $\frac{N_{01}}{N_{01}+N_{02}}$.

```
  # make relative
  N1E.pred.rel = matrix(NA, nrow=n.post, ncol=n.pred)
  N2E.pred.rel = matrix(NA, nrow=n.post, ncol=n.pred)
  for(k in 1:n.post){
    N1E.pred.rel[k, ] = N1E.pred[k, ]/(N1E.pred[k, ]+N2E.pred[k, ])
    N2E.pred.rel[k, ] = N2E.pred[k, ]/(N1E.pred[k, ]+N2E.pred[k, ])
  }

  N1E.pred.qs = apply(N1E.pred, 2, function(x) quantile(x, probs=c(0.05, 0.5, 0.95)))
  N2E.pred.qs = apply(N2E.pred, 2, function(x) quantile(x, probs=c(0.05, 0.5, 0.95)))
  N1E.pred.rel.qs = apply(N1E.pred.rel, 2, function(x) quantile(x, probs=c(0.05, 0.5, 0.95))
  N2E.pred.rel.qs = apply(N2E.pred.rel, 2, function(x) quantile(x, probs=c(0.05, 0.5, 0.95))

  plot(df.multi$Prey1/(df.multi$Prey1+df.multi$Prey2),
       df.multi$Prey1.Eaten/(df.multi$Prey1.Eaten+df.multi$Prey2.Eaten),
       xlim=c(0,1), ylim=c(0,1),
       xlab="Proportion prey 1 offered", ylab="Proportion prey 1 eaten", main="Multi-species
  abline(0,1)
  polygon( c(N10.pred/N0.total, rev(N10.pred/N0.total)),
```

```
        c(N1E.pred.rel.qs[1, ], rev(N1E.pred.rel.qs[3, ])),
        border=NA,
        col=adjustcolor("blue", alpha.f=0.2)
)
lines(N10.pred/N0.total, N1E.pred.rel.qs[2, ], col="blue", lwd=2)
```

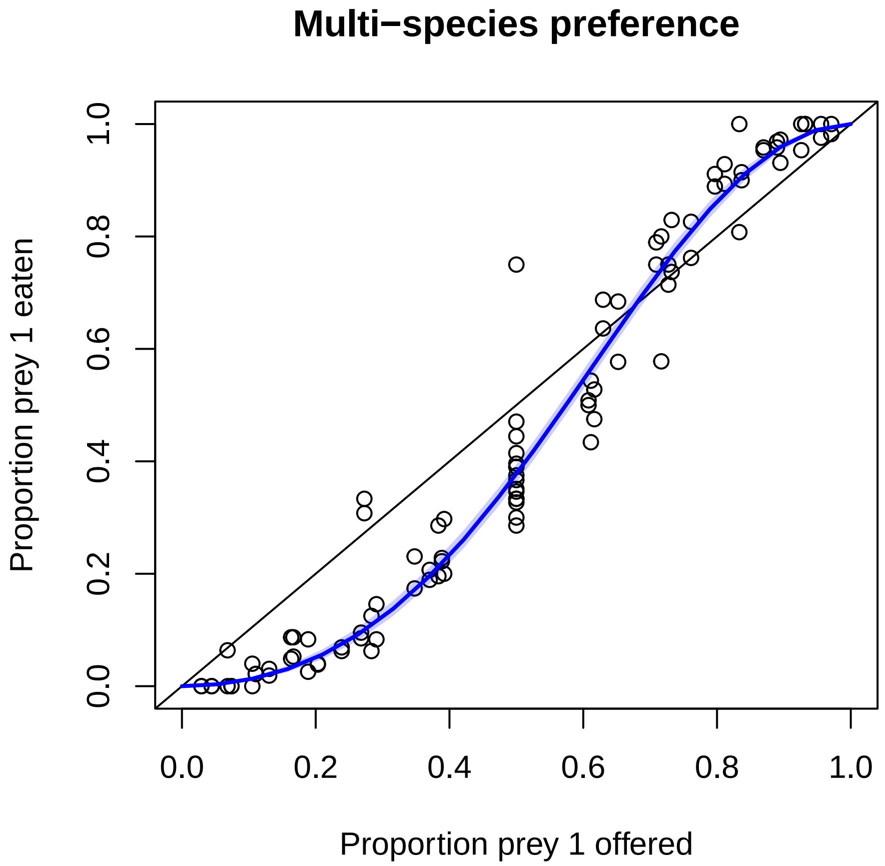

Note however that plotted observations originate from different levels of total offered prey $N_{0,1} + N_{0,2}$, while predictions used only one level of total offered prey $N_{0,1} + N_{0,2} = 200$.

###### Multi-species observed vs. predicted

Additionally, predictions are computed for each (multi-species) feeding trial from the dataset `df.multi`. Medians of predictions are plotted against observed number for eaten prey of both species in a classical observed vs. predicted plot.

```
N1E.pred = matrix(NA, nrow=n.post, ncol=nrow(df.multi))
N2E.pred = matrix(NA, nrow=n.post, ncol=nrow(df.multi))

for(k in 1:n.post){
  params = list(a=post[k, 1:2],
                h=post[k, 3:4],
                w=post[k, 5:6],
                r=post[k, 7],
                m=2)
  for(i in 1:nrow(df.multi)){
    if(df.all$Replaced[i]==FALSE){
      out = ode(y = c(N=c(df.multi$Prey1[i],df.multi$Prey2[i])), times = c(0,df.multi$Time[illegible]
      N1E.pred[k,i] = df.multi$Prey1[i]-out[2,2]
      N2E.pred[k,i] = df.multi$Prey2[i]-out[2,3]
    } else {
      out = (-1)*unlist(MSFR(N=c(df.multi$Prey1[i],df.multi$Prey2[i]), parms=params))
      N1E.pred[k,i] = out[1]*df.multi$Time[i]
      N2E.pred[k,i] = out[2]*df.multi$Time[i]
    }
  }
}

N1E.pred.qs = apply(N1E.pred, 2, function(x) quantile(x, probs=c(0.05, 0.5, 0.95)))
N2E.pred.qs = apply(N2E.pred, 2, function(x) quantile(x, probs=c(0.05, 0.5, 0.95)))

plot(NULL,
     xlim=c(0,max(df.all[, 3:4])), ylim=c(0,max(df.all[, 3:4])),
     xlab="Predicted", ylab="Observed", main="Multi-species obs vs. pred")
points(N1E.pred.qs[2, ], df.multi$Prey1.Eaten, col="blue")
abline(0,1)
points(N2E.pred.qs[2, ], df.multi$Prey2.Eaten, col="red")
abline(0,1)
```

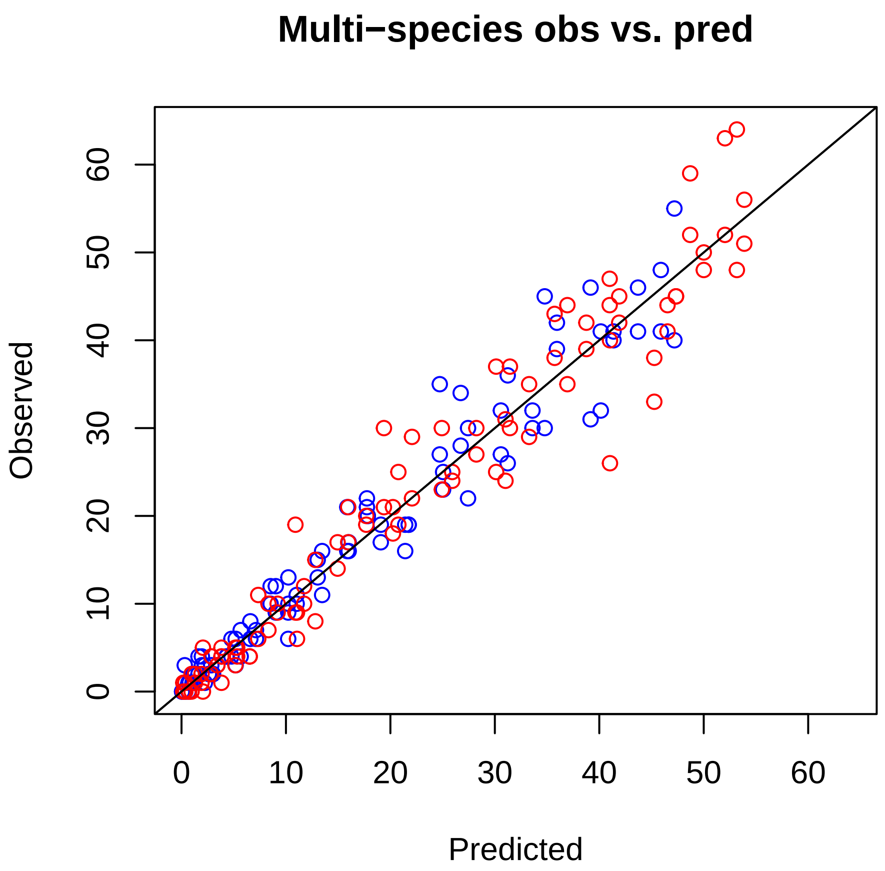

###### Stan code

The `MSFR_H2.stan` files etc. contain raw Stan code for all four models, they were compiled and compiled objects were saved in files `stan_fit_H2.RData` etc. This section contains a technical description, such that users can adapt the model if needed, e.g. to include different MSFR formulations or change parameters' prior distributions. Each single Stan model contains several blocks, which are presented individually below. We here only present the Holling-2 model, other models work analogously. Although the example above included 2 prey species, code works for $m$ species as well.

###### functions{ } block

This function describes the (negative of) functional response $F_i = \frac{a_i N_i}{1+\sum_{j=1}^{m} a_j h_j N_j}$, $i = 1 \dots m$. It is later used to compute predictions of eaten prey $N_{E,i}$ with the ODE method $\frac{dN_i}{dt} = -F_i(N_1 \dots N_m)$ if prey were not replaced (numerical integration in $t \in [0, T]$), or direct predictions $F_i(N_1 \dots N_m) \cdot T$ if prey were replaced during the feeding trials.

Stan requires a specific format for the application of the numerical ODE solver. The output is a vector of length `m`. Input arguments are time `t`, prey abundance `y` ($N_1 \ldots N_m$), model parameters `theta` ($a_1 \ldots a_m, h_1 \ldots h_m$), and other data (here number of species `m`). Additionally, a "fail-safe" sets feeding rates to zero if prey abundances are small enough. This prevents numerical issues of the ODE solver for small densities.

```
functions {
  vector MSFRm(real t, vector y, vector theta, int m) {
    vector[m] dydt;
    real denominator;
    denominator = 1.0;
    for(i in 1:m) denominator = denominator+(theta[i]*theta[m+i]*y[i]);
    for(i in 1:m) dydt[i] = -theta[i]*y[i] / denominator;
    for(i in 1:m){ if(y[i]<1e-6) dydt[i] = 0.0; }
    return dydt;
  }
}
```

##### data{ } block

All data have to be declared including data types and dimensions. Names have to be identical to the named list which was defined in R before.

```
data {
  int n; // number of feeding trials
  int m; // number of species
  array[n,m] int N_0; // offered prey
  array[n,m] int N_e; // eaten prey
  array[n] real Time; // experimantal duration
  array[n] int Replaced; // TRUE / FALSE read as 1 / 0
}
```

##### transformed data{ } block

Data transformations are computed here. `N_0_real` stores the integer numbers of offered prey as real numbers in a matrix. `nLL` is the number of non-zero offered prey which is just used in the `generated quantities{ }` block later.

```
transformed data {
  matrix[n,m] N_0_real;
  int nLL=0;
  for(i in 1:n)
    for(j in 1:m)
      N_0_real[i,j] = 1.0*N_0[i,j];
  for(i in 1:n)
    for(j in 1:m)
      if(N_0[i,j]>0)
        nLL=nLL+1;
}
```

###### parameters{ } block

All free model parameters are declared here. Attack rates and handling times must be positive, and a lower boundary of zero is provided.

```
parameters{
  vector<lower=0>[m] a;
  vector<lower=0>[m] h;
}
```

###### model{ } block

This block defines how the posterior distribution is computed, using prior distributions and the likelihood function. `N_p` is an auxiliary variable to save predictions for a single feeding trial. In case on non-replaced prey, these are the output ODE simulations with the `MSFRm` function. THe ODE solver function `ode_rk45()` requires them to be an array (length 1 for one prediction at time `Time[i]`) containing a vector (of length `m` number of species). All model parameter are coded as one single vector `pars`, as an input for the `MSFRm` function.

```
model{
  array[1] vector[m] N_p; // predictions
  vector[2*m] pars;       // parameters for ODE
```

Next, prior distributions for the model parameters are provided. Here, weak priors are chosen: exponential distributions with mean and standard deviation of 1 for attack rates and 1/10 for handling times. Note that these generic choices might have to altered depending on the units of the data. Here, attack rates are per unit time, and handling times are expressed in unit of time. E.g., transforming experimental duration `Time` from 1 [day] to 24 [hours], attack rates [per hour] will be 24 times smaller, and handling times [in hours] will be 24 times longer.

```
  a ~ exponential(1);
  h ~ exponential(10);

  for (i in 1:m){
    pars[i] = a[i];
    pars[m+i] = h[i];
  }
```

This is a for-loop over all $i = 1 \dots n$ feeding trials. If prey were replaced, predictions are directly computed by $F_j(N_{0,1} \dots N_{0,m}) \cdot T$ (note that the `MSFRm` function has negative feeding rate as output, and is converted to positive here). The observed number of eaten prey $N_{E,1} \dots N_{E,m}$ of all $m$ species is assumed to be Poisson distributed around the mean prediction. If the number of offered prey of a species $N_{0,j}$ is zero (i.e. in single-prey experiments of another prey species), both the observed and predicted number of eaten prey of this species is zero. This datapoint does not contain any information and is not included in the likelihood.

```
  for(i in 1:n){
    if(Replaced[i]>0){
      N_p = { MSFRm(0.0, N_0_real[i, ]', pars, m) };
      for(j in 1:m)
        if(N_0[i,j]>0)
          N_e[i,j] ~ poisson(-Time[i]*N_p[1][j]);
    }
```

If prey were not replaced, predicted numbers of (alive) prey of both species at the end of the experiment $N_{T,j}$ are computed with the numerical ODE solver `ode_rk45()`. Observed numbers of eaten prey are assumed to follow binomial distributions with $N_{0,j}$ trials and success probability $\frac{N_{0,j} - N_{T,j}}{N_{0,j}}$ ($N_{0,j} - N_{T,j}$ being the predicted number of eaten prey).

```
    else {
      N_p = ode_rk45(MSFRm, N_0_real[i, ]', 0.0, {Time[i]}, pars, m);
      for(j in 1:m)
        if(N_0[i,j]>0)
          N_e[i,j] ~ binomial(N_0[i,j], (N_0[i,j]-N_p[1][j])/N_0[i,j] );
    }
  } // end for(i in 1:n)
} // end model
```

##### generated quantities{ } block

This block is **only needed for model comparison** using the loo-package. Above steps of computing predictions are repeated, and log-likelihood values are stored in the vector `log_lik`. Only entries with positive numbers of offered prey are used, therefore its length `nLL` might be smaller than `n*m`.

```
generated quantities{
  array[nLL] real log_lik;{
    int k=0;
    array[1] vector[m] N_p; // predictions
    vector[2*m] pars;       // parameters for ODE
    for (i in 1:m){
      pars[i] = a[i];
      pars[m+i] = h[i];
    }
    for(i in 1:n){
      if(Replaced[i]>0){
        N_p = { MSFRm(0.0, N_0_real[i, ]', pars, m) };
        for(j in 1:m)
          if(N_0[i,j]>0){
            k=k+1;
            log_lik[k] = poisson_lpmf(N_e[i,j] | -Time[i]*N_p[1][j] );
          }
      } else {
        N_p = ode_rk45(MSFRm, N_0_real[i, ]', 0.0, {Time[i]}, pars, m);
        for(j in 1:m)
          if(N_0[i,j]>0){
            k=k+1;
            log_lik[k] = binomial_lpmf(N_e[i,j] | N_0[i,j], (N_0[i,j]-N_p[1][j])/N_0[i,j]
          }
      }
    }
  }
}
```

#### Session info

```
sessionInfo()
```

```
R version 4.3.2 (2023-10-31)
Platform: x86_64-pc-linux-gnu (64-bit)
Running under: Ubuntu 20.04.6 LTS

Matrix products: default
BLAS:   /usr/lib/x86_64-linux-gnu/openblas-pthread/libblas.so.3
LAPACK: /usr/lib/x86_64-linux-gnu/openblas-pthread/liblapack.so.3;  LAPACK version 3.9.0

locale:
 [1] LC_CTYPE=en_US.UTF-8       LC_NUMERIC=C
 [3] LC_TIME=de_DE.UTF-8        LC_COLLATE=en_US.UTF-8
 [5] LC_MONETARY=de_DE.UTF-8    LC_MESSAGES=en_US.UTF-8
 [7] LC_PAPER=de_DE.UTF-8       LC_NAME=C
 [9] LC_ADDRESS=C               LC_TELEPHONE=C
[11] LC_MEASUREMENT=de_DE.UTF-8 LC_IDENTIFICATION=C

time zone: Europe/Berlin
tzcode source: system (glibc)

attached base packages:
[1] stats     graphics  grDevices utils     datasets  methods   base

other attached packages:
[1] randtoolbox_2.0.4   rngWELL_0.10-9      deSolve_1.40
[4] BayesianTools_0.1.8 coda_0.19-4         loo_2.6.0
[7] rstan_2.32.5        StanHeaders_2.32.5

loaded via a namespace (and not attached):
 [1] bridgesampling_1.1-2 utf8_1.2.4           generics_0.1.3
 [4] stringi_1.8.3        lattice_0.22-5       lme4_1.1-35.1
 [7] digest_0.6.34        magrittr_2.0.3       evaluate_0.23
[10] grid_4.3.2           mvtnorm_1.2-4        fastmap_1.1.1
[13] Matrix_1.6-5         jsonlite_1.8.8       pkgbuild_1.4.3
[16] backports_1.4.1      gridExtra_2.3        Brobdingnag_1.2-9
[19] fansi_1.0.6          QuickJSR_1.1.0       scales_1.3.0
[22] codetools_0.2-19     cli_3.6.2            rlang_1.1.3
[25] munsell_0.5.0        splines_4.3.2        yaml_2.3.8
[28] tools_4.3.2          inline_0.3.19        parallel_4.3.2
[31] nloptr_2.0.3         checkmate_2.3.1      minqa_1.2.6
[34] dplyr_1.1.4          colorspace_2.1-0     ggplot2_3.4.4
[37] boot_1.3-28.1        curl_5.2.0           vctrs_0.6.5
[40] R6_2.5.1             matrixStats_1.2.0    stats4_4.3.2
[43] lifecycle_1.0.4      stringr_1.5.1        V8_4.4.1
```

```
[46] MASS_7.3-60.0.1      pkgconfig_2.0.3      RcppParallel_5.1.7
[49] pillar_1.9.0         gtable_0.3.4         glue_1.7.0
[52] Rcpp_1.0.12          xfun_0.41            tibble_3.2.1
[55] tidyselect_1.2.0     rstudioapi_0.15.0    knitr_1.45
[58] nlme_3.1-164         htmltools_0.5.7      rmarkdown_2.25
[61] DHARMa_0.4.6         compiler_4.3.2       IDPmisc_1.1.20
```
